# Nano-scale solution of the Poisson-Nernst-Planck (PNP) equations in a fraction of two neighboring cells reveals the magnitude of intercellular electrochemical waves

**DOI:** 10.1101/2022.09.07.506904

**Authors:** Karoline Horgmo Jæger, Ena Ivanovic, Jan P. Kucera, Aslak Tveito

## Abstract

The basic building blocks of the electrophysiology of cardiomyocytes are ion channels integrated in the cell membranes. Close to the ion channels there are very strong electrical and chemical gradients. However, these gradients extend for only a few nano-meters and are therefore commonly ignored in mathematical models. The full complexity of the dynamics is modelled by the Poisson-Nernst-Planck (PNP) equations but these equations must be solved using temporal and spatial scales of nano-seconds and nano-meters. Here we report solutions of the PNP equations in a fraction of two abuttal cells separated by a tiny extracellular space. We show that when only the potassium channels of the two cells are open, a stationary solution is reached with the well-known Debye layer close to the membranes. When the sodium channels of the left cell are opened, a very strong and brief electrochemical wave emanates from the channels. If the extracellular space is sufficiently small and the number of sodium channels is sufficiently high, the wave extends all the way over to the neighboring cell and may therefore explain cardiac conduction even at very low levels of gap junctional coupling.

## 1 Introduction

At the tissue level (~mm), cardiac electrophysiology is properly represented using the well-established bidomain or monodomain models; see, e.g., [1]. However, a major limitation of the bidomain and monodomain models is that the cardiomyocytes are not explicitly present in the models. This limits the modeling capabilities of these models. At the level of individual cardiomyocytes (~ μm), cell-based (EMI) models representing both the extracellular (E) space, the cell membrane (M) and the intracellular (I) space can be applied; see, e.g., [2, 3, 4, 5]. The EMI models increase the modeling capabilities, allowing parameters to vary between and within individual cardiomyocytes, at the cost of significantly increased computing efforts needed to solve the equations; see, e.g., [6]. Yet, even though individual ion channels can be represented in the EMI model framework, the model does not represent the strong gradients in ion concentrations close to these channels. In order to study the electrodiffusion close to active ion channels placed in the cell membrane, it is necessary to solve the Poisson-Nernst-Planck equations (PNP, see, e.g., [7, 8]) which models electrodiffusion at ~nm level. These equations are challenging to solve numerically because very strong gradients necessitate extremely fine spatial (~0.5 nm) and temporal (~ 0.01 ns) resolutions. The PNP equations can be simplified by assuming electroneutrality (charges sum up to zero everywhere in the computational domain), see, e.g., [9, 10, 11, 12, 13]. However, electroneutrality does not necessarily hold close the outlet (or inlet) of active ion channels and it remains unclear what consequences follow from assuming electroneutrality. A nice summary of the validity of different models close to the cell membranes is provided in [14] and limitations of electroneutrality are analyzed in [15].

Conventionally, electrochemical coupling between neighboring ventricular cardiomyocytes is believed to occur via gap junctions (GJ) that provide low-resistance pathways from cell to cell using the Connexin-43 (Cx43) protein, see, e.g., [16, 17]. An alternative route of conduction from cell to cell is via the extracellular space, referred to as ephaptic coupling. This alternative has been discussed for a very long time (see, e.g., [18, 19]) and recent modeling results, emphasizing the importance of the localization of the ion channels, indicate that ephaptic coupling is a viable alternative way of conduction, see, e.g., [20, 21, 3, 22]. A related discussion goes on in computational neurophysiology where the question is whether or not a neuron can set off an excitation wave in a neighboring cell; see [23, 24, 25, 26, 27]. One inherent difficulty in elucidating these questions is that the space between cells can be very small (~nm), making it very difficult to measure the electrical potential without perturbing the dynamics by the measuring device; see, e.g., [28].

The purpose of our report is to present solutions of the PNP equations between and in a fraction of two neighboring cells. In the simulations, we observe very strong spatial (~ 50 mV/nm) and temporal (~ 25 mV/ns) gradients in the electrical potential and this implies that we need to use an extremely fine mesh and very small time steps. We are thus only able to simulate a very small fraction of the cells and the extracellular space between them (~ 0.2 μm^3^) for a very brief time interval (~ 1 μs). For comparison, the volume of a ventricular myocyte is about 30,000 μm^3^ and the upstroke takes about 1 ms. On the other hand, the cross-sectional area of an ion channel is about 4 nm^2^ (see [29]) and we are able to represent that type of areas in our simulations.

When only the potassium channel cluster is open in both cells, a stationary solution is reached with the well-known Debye layer close to both cell membranes. From this stationary solution, we open the sodium channels in the left cell. This leads to a very strong electrochemical wave emanating from the sodium channels of the left cell. The strength of this wave depends on the number of sodium channels in the cluster of channels under consideration and also on the width of the extracellular space. When the extracellular space is sufficiently small and the number of sodium channels is sufficiently high, the wave emanating from the sodium channels of the left cell, reaches the right cell and can be strong enough to set off an excitation in the right cell.

## 2 Methods

### 2.1 The Poisson-Nernst-Planck (PNP) equations

We perform simulations of a small part of an intercalated disc. We consider a computational domain consisting of a fraction of two intracellular domains, with two associated membrane domains and an extracellular domain located between the membrane domains, as illustrated in Figure 1A. In the membrane domains, clusters of K^+^ and Na^+^ channels are embedded, as illustrated for a single K^+^ and Na^+^ channel in Figure 1B.

**Figure 1:**
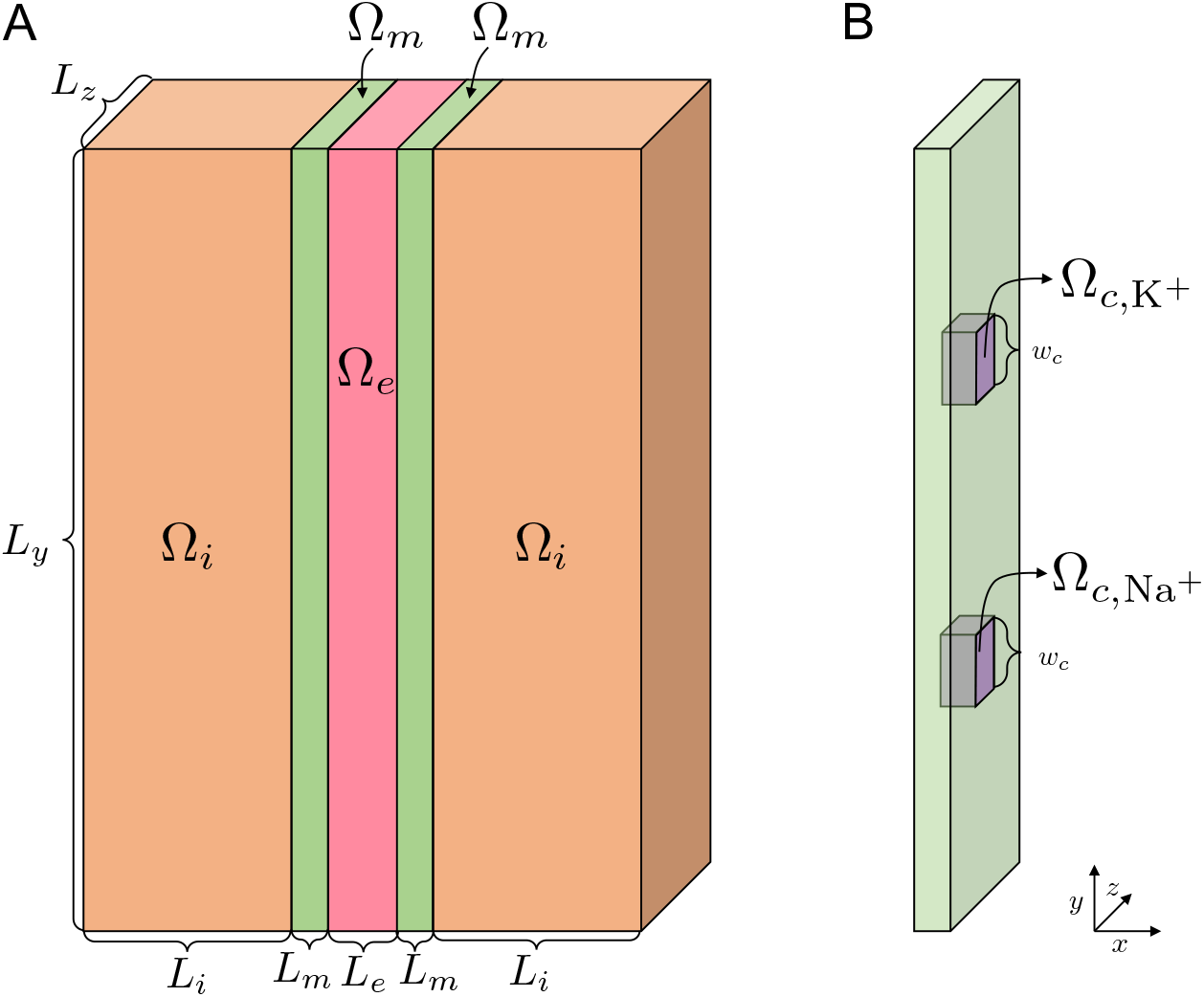
**A**. Illustration of the computational domain. We consider two intracellular domains Ω_*i*_ (orange), two membrane domains Ω_m_ (green) and an extracellular domain Ω_*e*_ (pink). **B**. Illustration of the channels embedded in the membrane, Ω_*c*,K^+^_ and Ω_*c*,Na^+^_ (purple). The width of the channels is *w_c_* in the *y*- and *z*-directions. Note that in our simulations, we typically consider clusters of channels, as illustrated in Figure 2. Note also that the illustrations in this figure are not drawn in scale, but the values of the lengths defining the geometry used in the simulations are specified in Table 2.

In this domain, the electric potential and the concentration of the four ionic species Na^+^, K^+^, Ca^2+^, and Cl^-^ are modeled using the Poisson-Nernst-Planck (PNP) model:

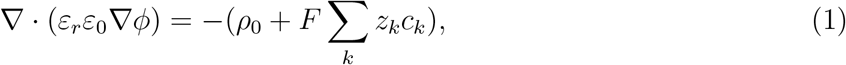

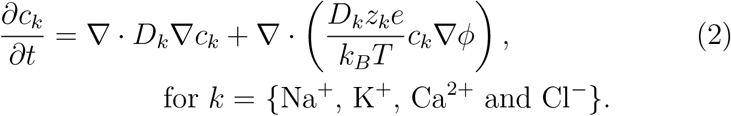

Here, *ϕ* is the electric potential (in mV) and *c_k_* are the ion concentrations (in mM). Furthermore, *F* is Faraday’s constant (in C/mol), *ε*_0_ is the vacuum permittivity (in fF/m), *ε_r_* is the (unitless) relative permittivity of the medium, *ρ*_0_ is the background charge density (in C/m^3^), *D_k_* are the diffusion coefficients for each ion species (in nm^2^/ms), *e* is the elementary charge (in C), *k_B_* is the Boltzmann constant (in mJ/K), *T* is the temperature (in K), and *z_k_* are the (unitless) valences of the ion species. Furthermore, time is given in ms and length is given in nm. Note that in the vizualizations below, we use ns as the temporal unit because of the extreme swiftness of the dynamics involved.

Defining

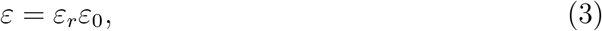

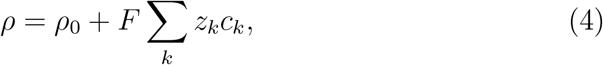

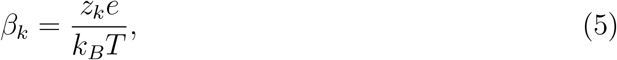

the system can be rewritten more compactly as

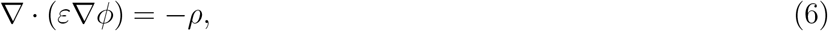

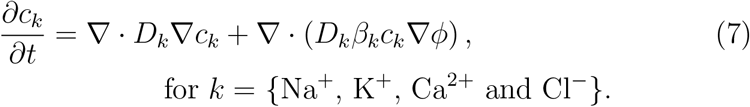

#### 2.1.1 Initial conditions and background charge density, *ρ*_0_

The initial conditions for the ion concentrations in the intracellular and extracellular domains are specified in Table 1. In the membrane (outside ion channels), all ion concentrations are set to zero. Furthermore, in the ion channels, the concentration of the ion species that is able to move through the channel is initially set up to vary linearly from the intracellular to the extracellular part of the channel, and the remaining ion concentrations are set to zero.

**Table 1:** Initial conditions for the ion concentrations, based on [11, 12].

| Ion | Intracellular | Extracellular |
| --- | --- | --- |
| Na<sup>+</sup> | 12 mM | 100 mM |
| K<sup>+</sup> | 125 mM | 5 mM |
| Ca<sup>2+</sup> | 0.0001 mM | 1.4 mM |
| Cl<sup>-</sup> | 137.0002 mM | 107.8 mM |

The background charge density, *ρ*_0_, is set up such that the entire domain is electroneutral at *t* = 0. This means that *ρ*_0_ = 0 everywhere except in the ion channels, where it is determined by the initial conditions for the ion species able to move through the channel.

In the supplementary information, we show the simulation results for a different choice of *ρ*_0_ than the default linear profile. More specifically, we consider initial conditions in the channels specified by a jump from the intracellular to the extracellular concentration in the center of the channel. The main results concerning the magnitude of the changes of the extracellular potential between the cells seem to be quite similar for this other choice of *ρ*_0_ in the channels. However, some differences between the solutions are observed (see the Supplementary Information).

#### 2.1.2 Parameter values

The default geometry used in the simulations is specified in Figure 1 and Table 2. In the purely intracellular and extracellular spaces, the relative permittivity, *ε_r_*, is set to *ε*_1_ = 80. In the membrane and the ion channels, *ε_r_* is set to *ε_m_* = 2. Furthermore, in the purely intracellular and extracellular domains, Ω_*i*_ and Ω_*e*_, the diffusion coefficients for the ions are as specified in Table 3. In the membrane domains, Ω_*m*_, all diffusion coefficients are set to zero. In the K^+^ channels, Ω_*c*,K^+^_, the diffusion coefficient for K^+^ is set to *d*_K^+^_ × *D*_K^+^_, where *d*_K^+^_ is a channel scaling factor. The diffusion coefficient for the remaining ions are set to zero in the K^+^ channels. Similarly, in the Na^+^ channels, Ω_*c*,Na^+^_, all diffusion coefficients are set to zero when the channel is closed and the diffusion coefficient for Na^+^ is set to the value *d*_Na^+^_ × *D*_Na^+^_ when the channel is open. The justification for the choice of scaling factors *d*_K^+^_ and *d*_Na^+^_ is specified below.

**Table 2:** Default geometry parameter values used in the simulations (see Figure 1).

| Parameter | Value |
| --- | --- |
| $L_i$ | 1000 nm |
| $L_m$ | 5 nm |
| $L_e$ | 5–30 nm |
| $L_y$ | 300 nm |
| $L_z$ | 300 nm |
| $w_c$ | 1 nm |

**Table 3:** Parameter values used in the simulations.

| Parameter | Description | Value | Ref. |
| --- | --- | --- | --- |
| $F$ | Faraday's constant | 96485.3365 C/mol | [30] |
| $\varepsilon_0$ | Vacuum permittivity | 8854 fF/m | [30] |
| $\varepsilon_1$ | Relative permittivity, $\varepsilon_r$, in $\Omega_i$ and $\Omega_e$ | 80 | [31] |
| $\varepsilon_m$ | Relative permittivity, $\varepsilon_r$, in $\Omega_m$, $\Omega_{c,Na^+}$ and $\Omega_{c,K^+}$ | 2 | [31] |
| $D_{Na^+}$ | Diffusion coefficient for $Na^+$ in $\Omega_i$ and $\Omega_e$ | $1.33 \cdot 10^6$ nm<sup>2</sup>/ms | [11] |
| $D_{K^+}$ | Diffusion coefficient for $K^+$ in $\Omega_i$ and $\Omega_e$ | $1.96 \cdot 10^6$ nm<sup>2</sup>/ms | [11] |
| $D_{Ca^{2+}}$ | Diffusion coefficient for $Ca^{2+}$ in $\Omega_i$ and $\Omega_e$ | $0.71 \cdot 10^6$ nm<sup>2</sup>/ms | [11] |
| $D_{Cl^-}$ | Diffusion coefficient for $Cl^-$ in $\Omega_i$ and $\Omega_e$ | $2.03 \cdot 10^6$ nm<sup>2</sup>/ms | [11] |
| $d_{K^+}$ | Scaling factor for $D_{K^+}$ in $\Omega_{c,K^+}$ | 0.025 | |
| $d_{Na^+}$ | Scaling factor for $D_{Na^+}$ in $\Omega_{c,Na^+}$ | 0.5 | |
| $z_{Na^+}$ | Valence of $Na^+$ | 1 | |
| $z_{K^+}$ | Valence of $K^+$ | 1 | |
| $z_{Ca^{2+}}$ | Valence of $Ca^{2+}$ | 2 | |
| $z_{Cl^-}$ | Valence of $Cl^-$ | -1 | |
| $e$ | Elementary charge | $1.60217662 \cdot 10^{-19}$ C | [30] |
| $k_B$ | Boltzmann constant | $1.380649 \cdot 10^{-20}$ mJ/K | [30] |
| $T$ | Temperature | 310 K | |

#### 2.1.3 Parameterization of the channels

##### Channel area and diffusion

We let the cross sectional area of each ion channel be 1 nm × 1 nm. In addition, we adjust the value of the scaling factor *d*_Na^+^_ for the diffusion through an open Na^+^ channel so that the peak current through an open channel is approximately 2 pA, based on single channel measurements from [32]. Furthermore, based on estimation from [33] that the conductance of *I*_K1_ channels is about 4 times smaller than the conductance of *I*_Na_ channels and that the number of *I*_Na_ channels is about 5 times larger than the number of *I*_K1_ channels, the scaling factor for the K^+^ channel diffusion, *d*_K^+^_, is set to be 20 times smaller than *d*_Na^+^_.

##### Representation of channel clusters

Na^+^ channels have been found to form clusters on the membrane of cardiomyocytes (see, e.g., [34, 35, 36]). In order to investigate how such channel clustering affects the simulation results, we consider some cases of different numbers of open Na^+^ channels in a cluster. The clusters are represented in the model by increasing the width, *w_c_* (and consequently the area) of the Na^+^ channels that are embedded in the membrane (see Figure 1B). The scaling factor for the Na^+^ channel diffusion, *d*_Na^+^_ is kept constant when the channel cluster size is increased. In our simulations, we consider Na^+^ channel clusters consisting of 4, 36, 196 or 900 Na^+^ channels (see Figure 2).

**Figure 2:**
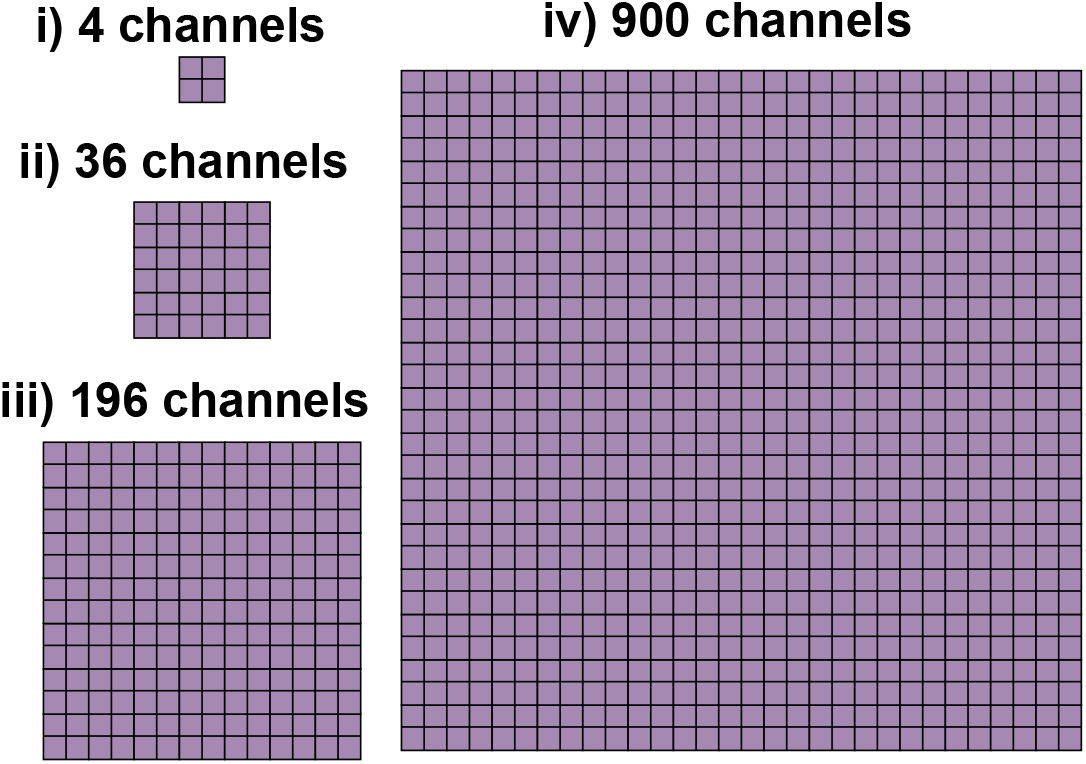
Illustration of the four different Na^+^ channel cluster sizes considered in our simulations. In the case of 4 channels, the channel cluster width is 2 nm, whereas in the case of 36, 196 and 900 channels, the width of the channel clusters are 6 nm, 14 nm and 30 nm, respectively.

For the K^+^ channel clusters, we consider a cluster of 36 channels for each of the two cells. When the size of the Na^+^ channel clusters are adjusted, we adjust the scaling factor for the K^+^ channel diffusion, *d*_K^+^_, by a factor 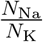, where *N*_Na_ is the number of Na^+^ channels and *N_K_* = 36 is the number of K^+^ channels. This is done to maintain the ratio of the magnitude of the *I*_Na_ and *I*_K1_ currents.

### 2.2 Simulation setup

Using the model setup described in Section 2.1, we perform simulations to investigate possible ephaptic coupling between the two cells. The simulations are separated into two parts:

#### Part 1: Closed Na^+^ channels

We start the simulation from electroneutral conditions (see Section 2.1.1) with physiological values for the extracellular and intracellular ion concentrations (see Table 1). We then let the model approach a resting state by letting only the K^+^ channels be open between the intracellular and extracellular spaces on the membrane of both cells. For the initial conditions (electroneutral), *ρ* is zero, and *ϕ* is constant (see (6) and (10)).

#### Part 2: Open Na^+^ channel cluster on the membrane of the left cell

After the simulation in Part 1 seems to have reached a steady state, we simulate the opening of the Na^+^ channel cluster on the membrane of the left cell and observe how this affects the ion concentrations and electric potential in the extracellular space between the cells. In particular, we wish to investigate whether the Na^+^ channel opening on the left cell seems to affect the electric potential outside the right cell enough to potentially initiate Na^+^ channel opening on the membrane of the right cell. The K^+^ channels are kept open in this part of the simulation. The Na^+^ channels on the left cell are kept open until the transmembrane potential, *v* (see Figure 5), across the Na^+^ channel cluster of the left cell reaches a value of 30 mV.

**Figure 3:**
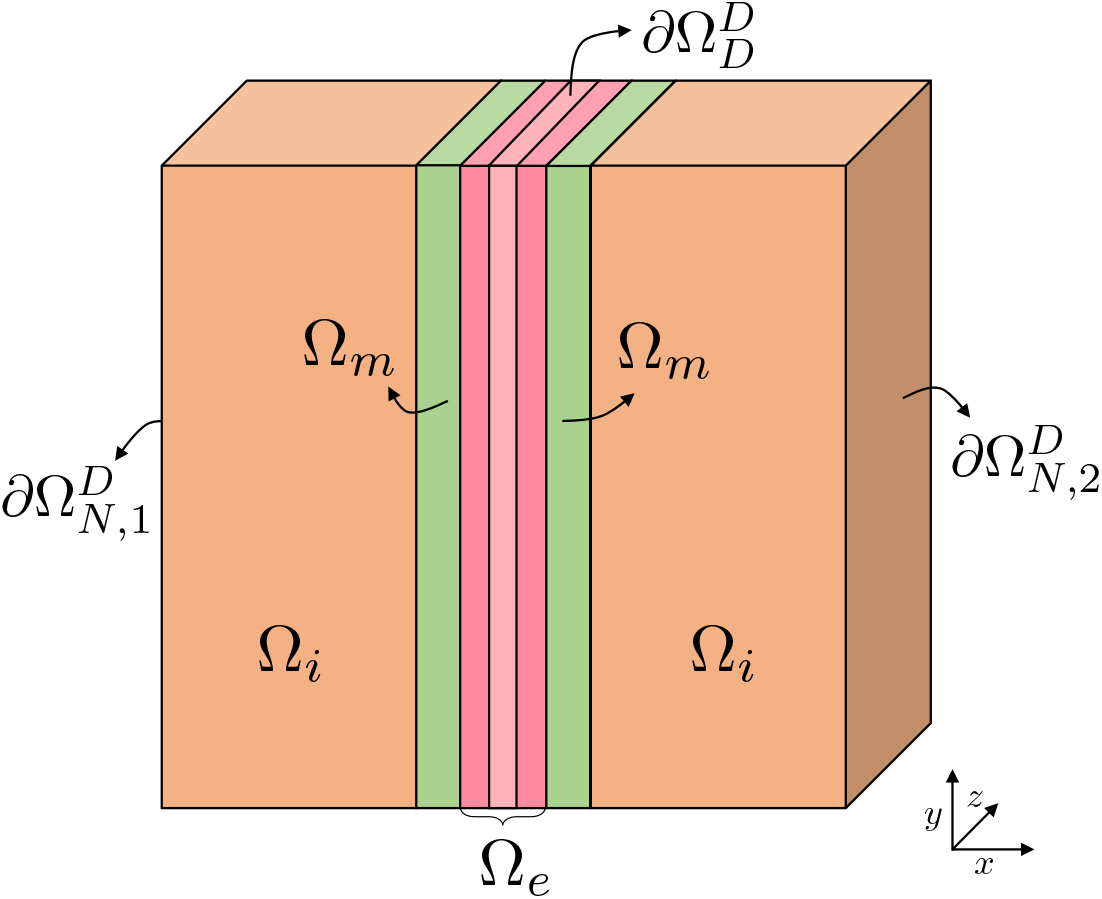
Illustration of different parts of the domain boundary involved in the physiologically motivated boundary conditions (see Section 2.3.2) used in most of our simulations. In 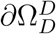, i.e., in the boundary in the *y*- and *z*-directions for the *x*-values corresponding to the middle third of the extracellular cleft, we apply Dirichlet boundary conditions for both *ϕ* and all the ionic concentrations. In the simulations used to find the resting state of the system (see Section 2.2), we apply Neumann boundary conditions for *ϕ* and Dirichlet boundary conditions for the concentrations at 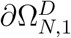 and 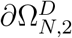. In the simulations with open Na^+^ channels (see Section 2.2), we apply Neumann boundary conditions for *ϕ* and Dirichlet boundary conditions for the concentrations at 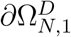 and Dirichlet boundary conditions for both the concentrations and *ϕ* at 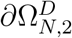. In the remaining boundary 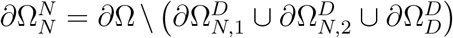, we apply Neumann boundary conditions for both ϕ and the ionic concentrations.

**Figure 4:**
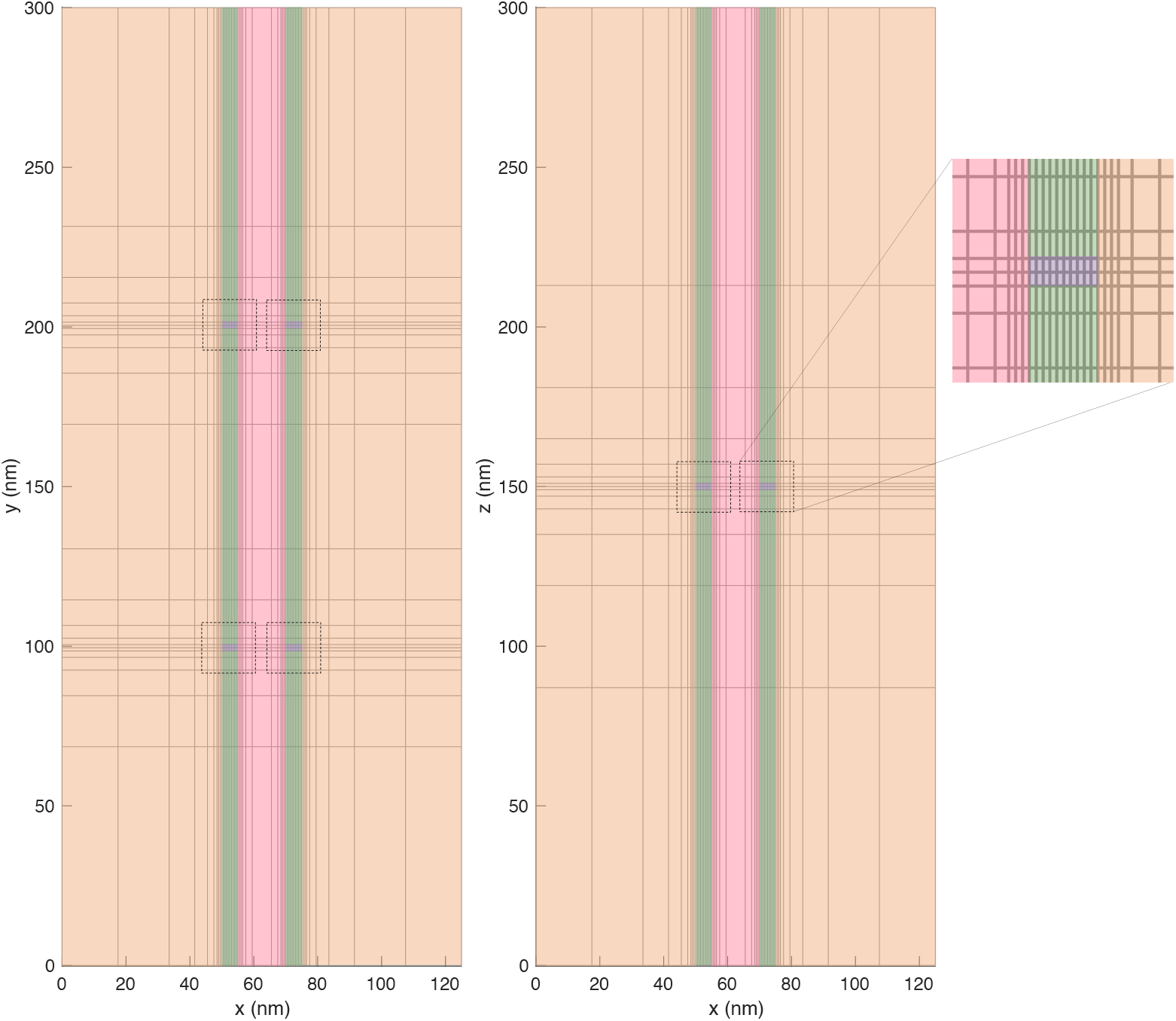
Illustration of an example mesh for *L_e_* = 15 nm and *L_i_* = 50 nm in the (*x, y*)- and the (*x, z*)-planes. The intracellular domain is colored orange, the extracellular domain is colored pink, the membrane is colored green and the ion channels are colored purple. In this figure, we consider K^+^ and Na^+^ channel clusters consisting of 2×2 channels of each type in the membrane of both cells. In the (*x, y*)-plane, the two types of ion channel are located as different locations (different *y*-values), whereas in the (*x, z*)-plane the two types of ion channel overlap because they are located at the same *z*-values. The mesh in the area close to the ion channels (indicated by dashed lines) is shown in more detail on the right side of the figure.

**Figure 5:**
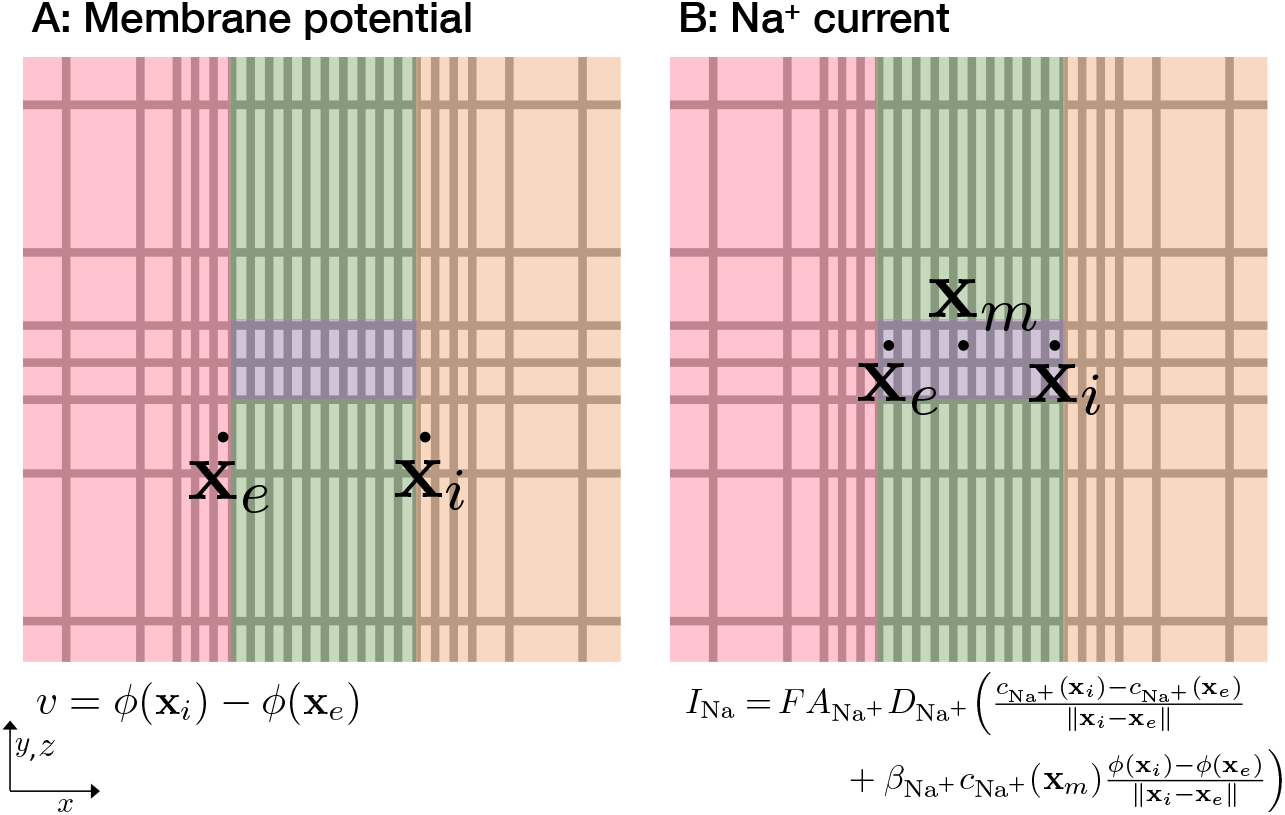
Illustration of the definition of the transmembrane potential, *v*, and the Na^+^ current, *I*_Na_, used in our computations. The illustrations show a small part of the mesh in the (*x, y*)- or (*x, z*)-planes close to the Na^+^ channel (as illustrated on the right side of Figure 4).

### 2.3 Boundary conditions

In order to simulate possible ephaptic coupling between two neighboring cells, we would like to solve the PNP system in a volume consisting of two cells and the surrounding extracellular space (i.e., about 60,000 *μ*m^3^). If we, for simplicity, assume that every computational mesh block is 1 nm^3^, this would lead to a computational problem with 6×10^13^ blocks, which is currently out of reach. We therefore need to restrict the problem spatially and that means that we need to define boundary conditions at locations where there are no actual physical boundaries. Of course, we want to define such boundary conditions in a way that provides the best possible model of the physiology under consideration. We will deal with combinations of Neumann and Dirichlet conditions and present results for two alternative sets of boundary conditions (one alternative will be shown only in the Supplementary Information).

#### 2.3.1 Natural boundary conditions for the PNP system

For the Poisson equation (6), a compatibility condition needs to be satisfied in order to obtain proper solutions of the equation, see, e.g., [37, 38, 39]. This condition follows from Gauss’ theorem after integration over the entire computational domain,

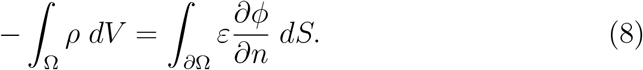

In the case of *natural* boundary conditions, i.e., when the normal derivatives of the both the electrical potential and the all the concentrations vanishes at the boundary, this condition holds. First, the right hand side is clearly zero (by the boundary condition on the electrical potential). Also, if we assume that the ions initially are in an electroneutral state in the whole domain, then the integral of *ρ*(·, *t* = 0) is zero. Furthermore, this integral does not change in time because the integral of each specie is constant,

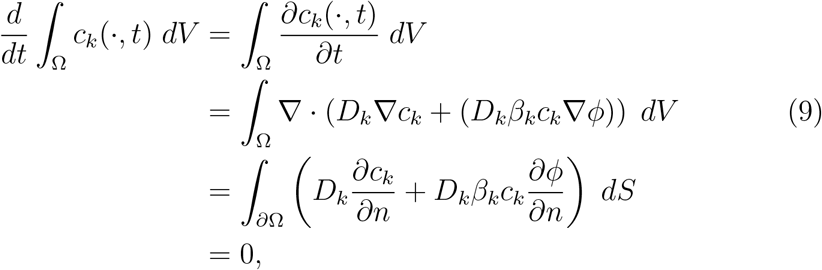

for any *k* = {Na^+^, K^+^, Ca^2+^ Cl^-^}. Since both the left hand side and the right hand side of (8) are zero, the compatibility condition is satisfied for natural boundary conditions.

#### 2.3.2 Physiologically motivated boundary conditions

The set of boundary conditions considered in most of our simulations is a physiologically motivated and relatively complex mix of Dirichlet and Neumann type boundary conditions. These boundary conditions are applied in all simulations reported in the Results section of the paper, and all simulations in the Supplementary Information, unless otherwise stated.

In the physiologically motivated boundary conditions, we apply a Dirichlet boundary condition fixing the electric potential, *ϕ*, at zero in an area of the boundary in the *y-* and *z*-directions in the extracellular cleft. More specifically, this area, 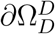 is defined for the *x*-values corresponding to the middle third of the extracellular cleft, as illustrated in Figure 3. In 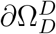, we apply Dirichlet boundary conditions for the all the ionic concentrations as well. The concentrations are in this area fixed at the initial conditions for the extracellular space (see Table 1). At the leftmost boundary of the left cell, marked as 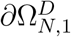 in Figure 3, we fix the concentrations at the initial conditions for the intracellular space see Table 1), and apply no-flux Neumann boundary conditions for *ϕ*. At the rightmost boundary of the right cell, marked as 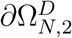 in Figure 3, we use the same boundary conditions as for 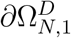 in the simulations used to find the resting state of the system (see Section 2.2). However, when the Na^+^ channels of the left cell are opened, we apply Dirichlet boundary conditions for *ϕ* on the right boundary of the right cell 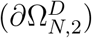, fixing the potential at this boundary at the value of *ϕ* found at the end of the resting state simulations. On the remaining part of the domain boundary, 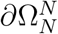, we apply no-flux Neumann boundary conditions for both *ϕ* and the concentrations.

In summary, the boundary conditions are given by

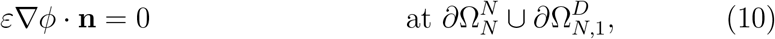

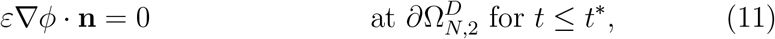

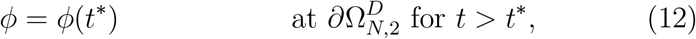

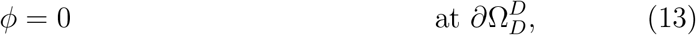

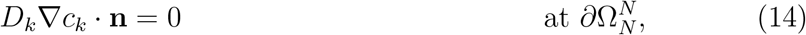

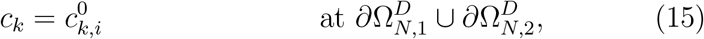

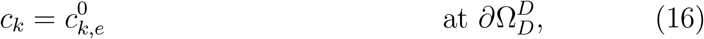

for *k* = {Na^+^, K^+^, Ca^2+^ and Cl^-^}, where 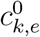 and 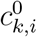 are the extracellular and the intracellular concentrations, respectively, of the ion species *k* specified in Table 1 and *t*\* is the point in time when the Na^+^ channels of the left cell are opened (see Section 2.2). Furthermore, 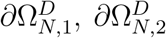 and 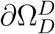 are illustrated in Figure 3, and 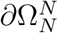 is the remaining part of the domain boundary, defined by 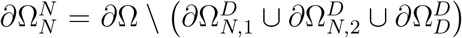, where *∂*Ω is the entire domain boundary.

Note that in the Supplementary Information, we show the results of simulations where Neumann boundary conditions are applied everywhere for all ionic concentrations and everywhere except at the right boundary for *ϕ*. The results of the simulations with this different choice of boundary condition seem to be very close to the results obtained using the boundary conditions described above.

### 2.4 Numerical solution of the PNP equations

We solve the PNP model equations using an implicit finite difference discretization of a splitting scheme in which the two equations (6) and (7) are solved separately. That is, for each time step n, we assume that the concentrations 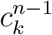 (and consequently *ρ*^*n*-1^) are known for *t*_*n*-1_ and the PNP model equations are solved in two steps:

**Step 1:** Solve

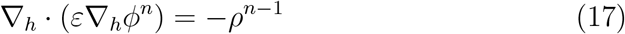

**Step 2:** Solve

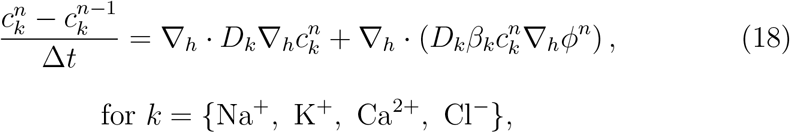

where ∇_*h*_ refers to a finite difference discretization of ∇. The finite difference discretizations used in Step 1 and Step 2 are described in detail in the Supplementary Information.

#### 2.4.1 Mesh

The spatial domain is discretized using a simple adaptive mesh in the *x*-, *y*- and *z*-directions. Close to the membrane and in the membrane, Δ*x* is set to 0.5 nm. Further away from the membrane, Δ*x* is doubled for each grid point. Similarly, Δ*y* and Δ*z* are set to 1 nm close to the ion channels and doubled for each grid point further away from the channels. An example grid is illustrated in the (*x, y*)- and the (*x, z*)-planes in Figure 4. This simple approach to create an adaptive mesh considerably reduces the number of mesh points and thus also the CPU efforts. For example, for the default domain with *L_y_* = *L_z_* = 300 nm, *L_i_* = 1000 nm, and *L_e_* = 10 nm, the number of grid points is reduced from about 364 · 10^6^ for a uniform mesh with Δ*x* = 0.5 nm and Δ*y* = Δ*z* = 1 nm to about 18,000 for the adaptive mesh. In other words, the number of unknowns to be computed for each time point is reduced by a factor of about 20,000.

In order to avoid unstable numerical solutions, we use the time step Δ*t* = 2 · 10^-8^ ms = 0.02 ns.

#### 2.4.2 Definition of the membrane potential and Na^+^ current

In our simulations, the transmembrane potential is defined as

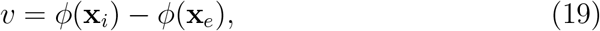

where **x**_*i*_ and **x**_*e*_ are the first grid points (in the *x*-direction) outside of the membrane in the intracellular and extracellular spaces, respectively, as illustrated in Figure 5A. Furthermore, the sodium current, *I*_Na_, is defined by the associated flux,

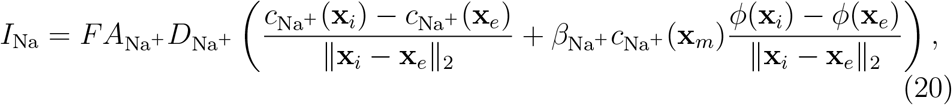

where **x**_*i*_ and **x**_*e*_ are the first grid points (in the *x*-direction) inside of the Na^+^ channel, close to the intracellular and extracellular spaces, respectively, and **x**_*m*_ is a point in the center of the channel as illustrated in Figure 5B. Moreover, *A*_Na^+^_ is the area of the membrane covered by the Na^+^ channel cluster, and || · ||_2_ is the Euclidean norm.

## 3 Results

### 3.1 The resting state of the PNP equations

Before investigating the dynamics involved with the opening of Na^+^ channels on the membrane one of the two cells included in the simulations, we run simulations to obtain a resting state for the model, as described in Section 2.2. We run this type of simulation for three different lengths of the extracellular space between the cells (*L_e_* = 5 nm, *L_e_* = 10 nm, and *L_e_* = 30 nm).

#### 3.1.1 The resting state depends on the size of the extracellular space

Figure 6 shows the extracellular potential, *ϕ*, the concentration of Na^+^, K^+^, Ca^2+^, and Cl^-^ ions and the charge density, *ρ*, in the extracellular space between the cells when the solutions seem to have reached steady state in the simulations performed to obtain the resting state of the system. In these simulations, only the K^+^ channels are open on the membrane of the two cells. We show the solutions in a plane in the *x*- and *y*-directions in the center of the domain in the *z*-direction. We consider the solutions close to the K^+^ channel, with the K^+^ channels located in the center of the plots in the *y*-direction. Close to the membrane of both cells, we observe a boundary layer with slightly different values of the potential, *ρ* and the concentrations than in the remaining extracellular space. This boundary layer extends further into the domain in the area close to the K^+^ channels.

**Figure 6:**
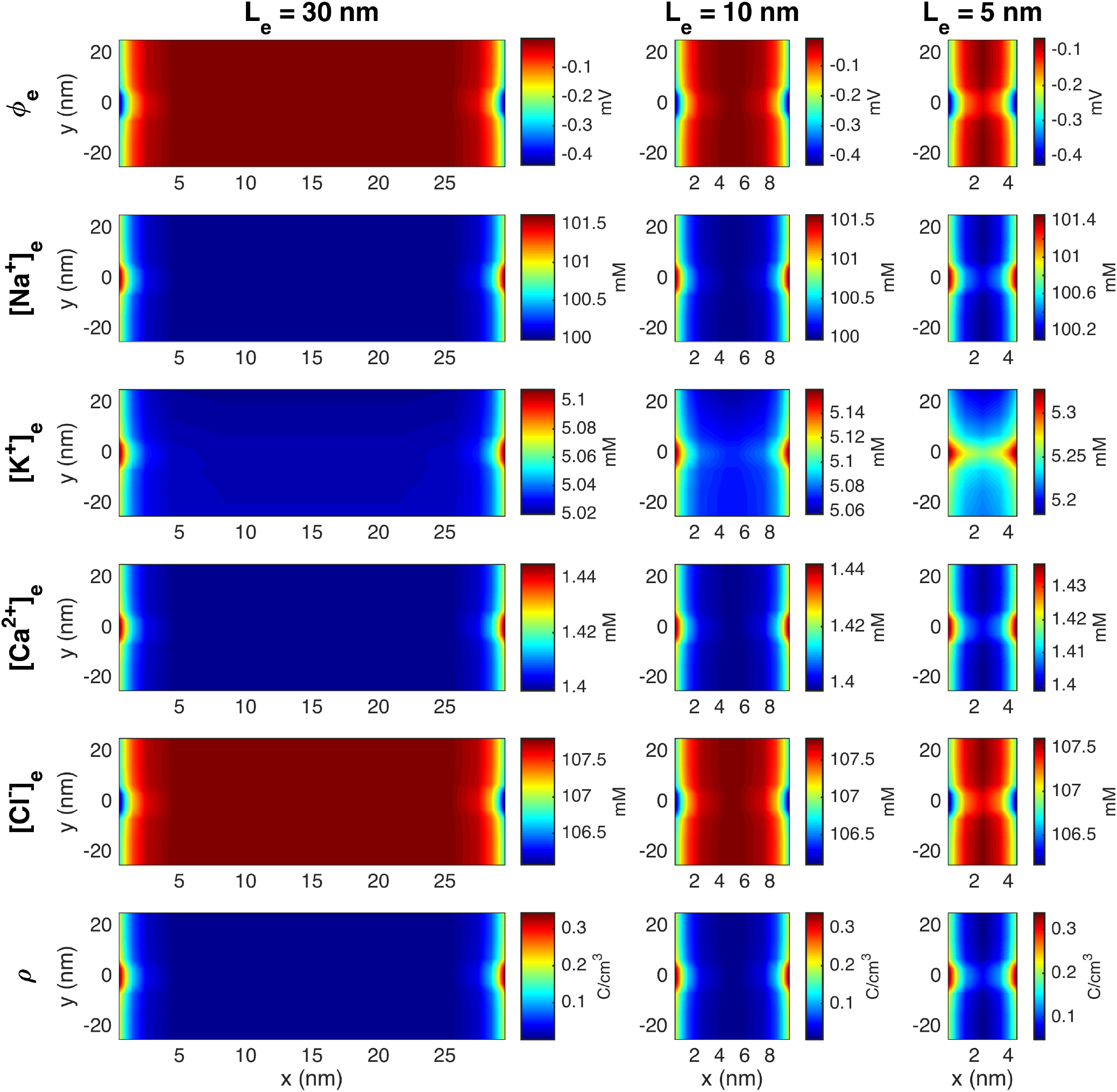
Stationary solution of the potential, *ϕ*, the concentration of Na^+^, K^+^, Ca^2+^, and Cl^-^ ions, and the charge density, *ρ* in the extracellular space between the two cells in simulations with open K^+^ channels, but closed Na^+^ channels. The width of the extracellular space, *L_e_*, is varied in the columns. The plots show the solution in the (*x, y*)-plane for the center of the domain in the *z*-direction at a point in time when steady state is reached. In the *y*-direction, we focus on the 50 nm closest to the K^+^ channels. The coordinates on the axes are shifted so that *x* = 0 marks the end of the membrane of the left cell and *y* = 0 marks the center of the K^+^ channels. Note that to improve the visibility of the boundary layer, the scaling of the colorbar is different for the different cases.

In Figure 7, we further illustrate the resting state solutions as lines in the x-direction through the K^+^ channels and through a line across the main membrane. We observe that the electric potential *ϕ* is about −80 mV in the intracellular space and about 0 mV in the extracellular space, resulting in a transmembrane potential of about −80 mV. We can also observe some small boundary layers in the ion concentrations close to the membrane. In the lower row of Figure 7, we show the charge density, *ρ*, computed from the ionic concentrations using (4). For the main membrane (dotted line), we clearly see that this value is close to zero everywhere, except in the boundary layer in both the intracellular and the extracellular spaces close to the membrane. In the line crossing the K^+^ channel cluster, we also observe a non-zero charge density inside and close to the channel cluster. The profile of *ρ* in the K^+^ channel cluster is a consequence of small deviations from the linear profile used as initial condition for the K^+^ concentration in the cluster and is discussed further in the Supplementary Information.

**Figure 7:**
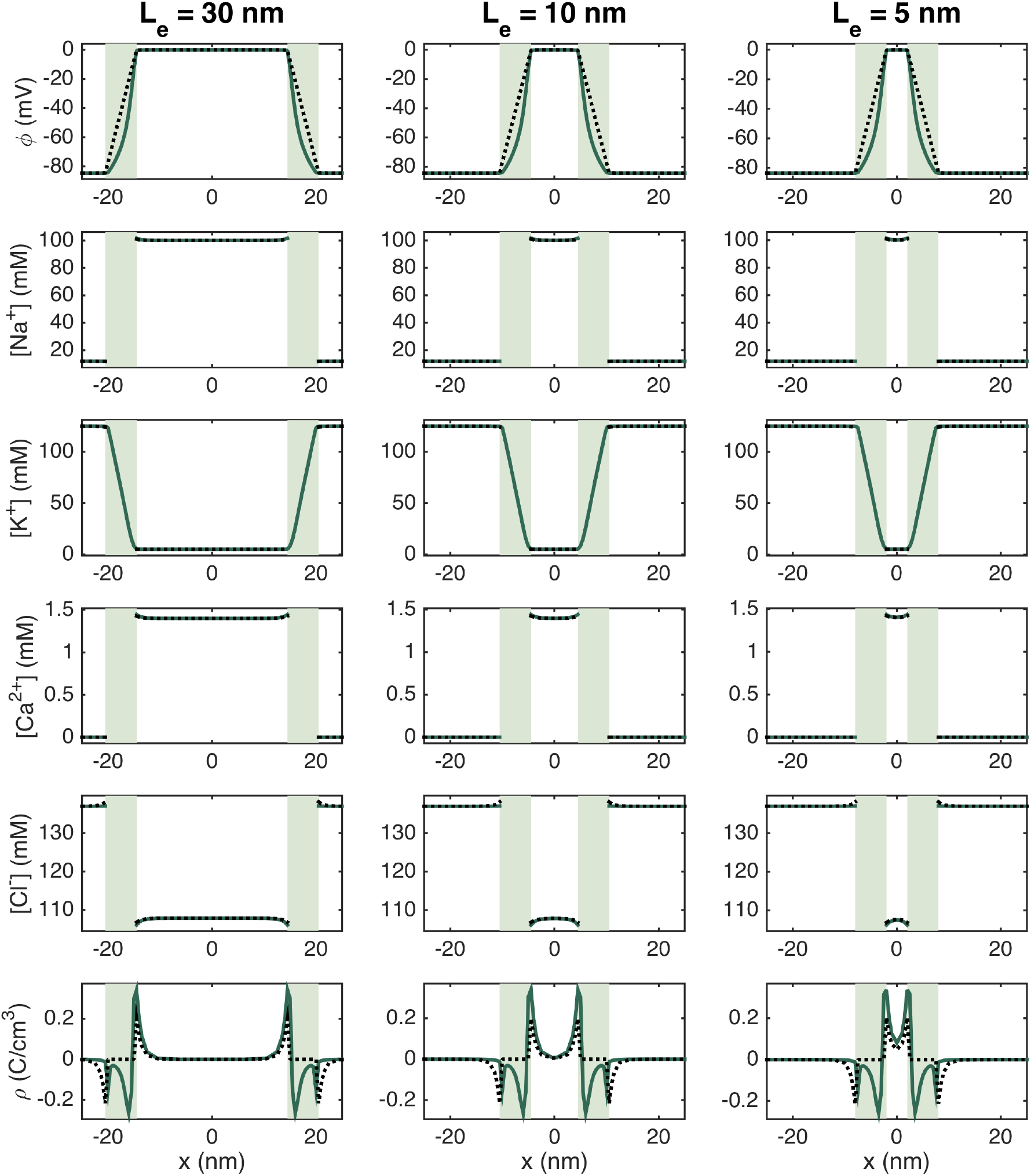
Stationary solution of the potential, *ϕ*, the concentration of Na^+^, K^+^, Ca^2+^, and Cl^-^ ions, and the charge density, *ρ*, along lines in the *x*-direction for open K^+^ channels and closed Na^+^ channels. The full green line represents the solution along a line crossing through the K^+^ channels and the dotted black line represents the solution along a line about 100 nm below the K^+^ channel cluster. The light green areas mark the membrane. Note that all ion concentrations are zero in the membrane, except for K^+^ ions in the K^+^ channel. The coordinates on the axes are shifted so that *x* = 0 marks the center of the extracellular cleft, and to improve the visibility, the plots only focus on a small part of the intracellular space, closest to the membrane.

### 3.2 Opening the Na^+^ channels on the left cell

Starting from the solution at the end of the resting state simulations, we run simulations with open Na^+^ channels in the membrane of the left cell.

#### 3.2.1 A strong and rapid transient emanates from the open Na^+^ channels

In Figure 8, we show the solution of a simulation with an extracellular space width of *L_e_* = 10 nm and a Na^+^ channel cluster consisting of 196 channels. We show the solutions in a plane in the *x*- and *y*-directions in the extracellular space close to the Na^+^ channel cluster at five different points in time. The first column of Figure 8 shows the solution 1 ns after the Na^+^ channel cluster of the left cell is opened. We observe that the Na^+^ concentration is somewhat decreased outside of the channel cluster, as Na^+^ ions have started to move into the cell through the open channel cluster. The charge density, ρ, is also negative outside of the Na^+^ channel cluster, and the extracellular potential is more negative outside of the channel cluster than at rest. Furthermore, the K^+^ and Ca^2+^ concentrations are slightly increased and the Cl^-^ concentration is slightly decreased in the vicinity of the open Na^+^ channel cluster.

**Figure 8:**
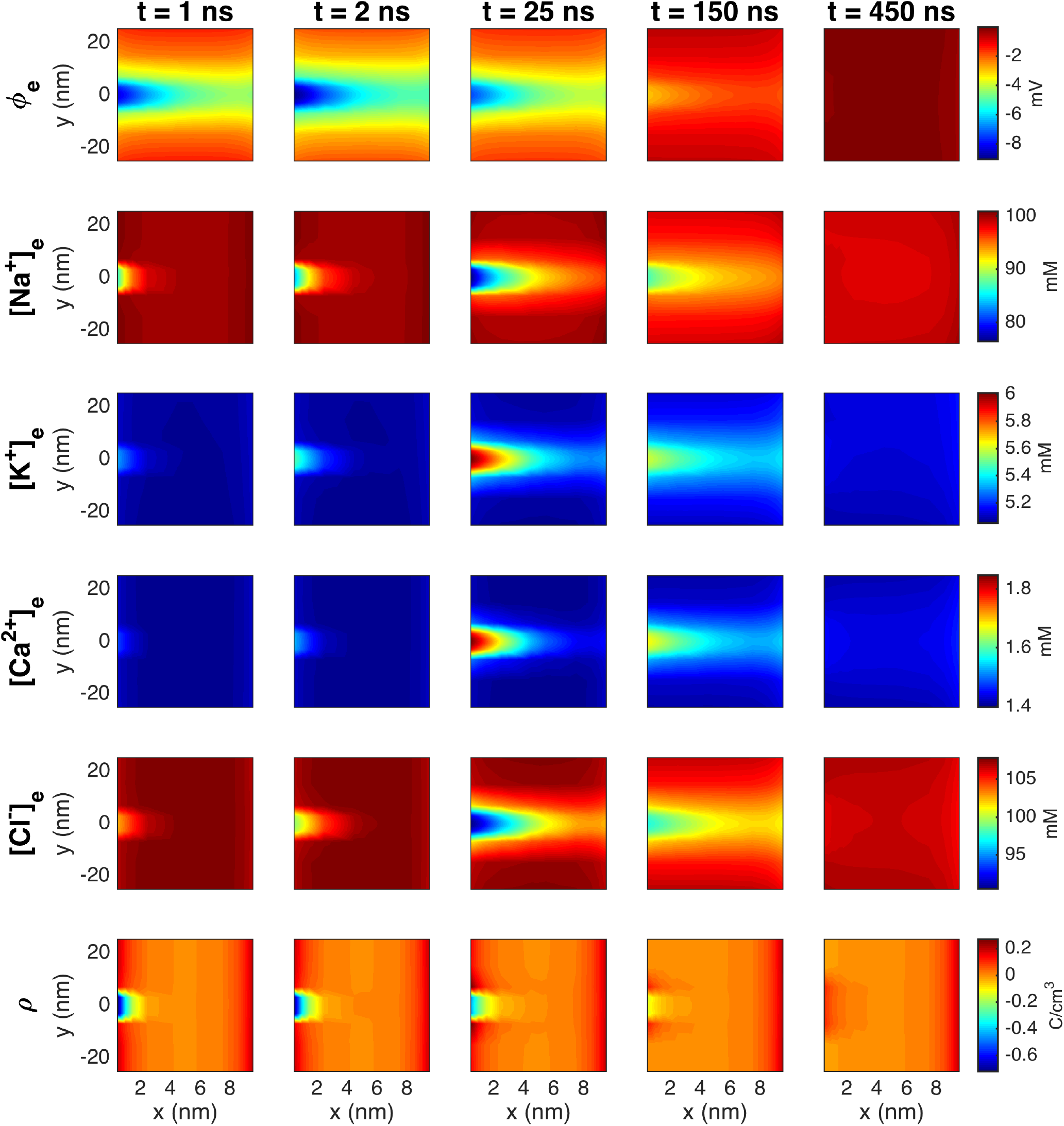
The PNP model solution in the extracellular space between two cells in a simulation with an Na^+^ channel cluster of 196 channels on the membrane of the left cell (the channels are opened at *t* = 0). The width of the extracellular space, *L_e_*, is 10 nm. The plots show the solution in the (*x, y*)-plane for the center of the domain in the *z*-direction at five different points in time (specified in the column titles). In the *y*-direction, we focus on the 50 nm closest to the Na^+^ channel clusters. The coordinates on the axes are shifted so that *x* = 0 marks the end of the membrane of the left cell and *y* = 0 marks the center of the Na^+^ channel cluster. Note that the short duration of the dynamics is an artefact caused by the small membrane area associated with the Na^+^ channel cluster in the simulation (see Sections 3.2.3–3.2.5 for more details and an estimation of a more realistic duration).

In the next column, at *t* = 2 ns, the changes to the ion concentrations are more pronounced and extend further into the extracellular space between the cells. In addition, the extracellular potential is at its most negative, with a value of about −9 mV in an area outside of the open Na^+^ channel cluster of the left cell. In the extracellular space outside of the membrane of the right cell, the potential is about −5 mV. The charge density outside of the channel cluster appears to be similar to that observed at 1 ns. The next column shows the solutions at the time when the changes to the ion concentrations peak, at *t* = 25 ns. Now, the ion concentration changes extend even further into the extracellular space. The changes in extracellular potential and the charge density, on the other hand, are decreasing. In the final two columns, the changes in both the extracellular potential, *ρ* and the ion concentrations decreases. Note that the duration of these dynamics in this simulation might not be physiologically realistic because of the small size of the simulated domain and the small size of the simulated membrane. This is investigated in Sections 3.2.3–3.2.5. In Section 3.2.5 a more realistic duration is estimated.

#### 3.2.2 The magnitude of the electrochemical wave depends strongly on the number of open sodium channels and the size of the extracellular space

Figure 9 shows the extracellular potential, *ϕ*, between the two cells at the point in time when the largest deviation from rest occurs after the Na^+^ channel cluster of the left cell has been opened for three different lengths of the extracellular space (*L_e_* = 5 nm, *L_e_* = 10 nm and *L_e_* = 30 nm) and four different sizes of the Na^+^ channel clusters (see Figure 2). We observe that for all the different Na^+^ channel cluster sizes, the magnitude (absolute value) of the most negative extracellular potential increases when the width of the extracellular space in decreased. Furthermore, we observe that for *L_e_* = 30 nm or *L_e_* = 10 nm, the magnitude of the extracellular potential is considerably lower outside of the right cell than outside of the left cell (with open Na^+^ channels), while for *L_e_* = 5 nm, the magnitude of the extracellular potential appears to be quite similar across the extracellular space between the cells. This indicates that the strong negative extracellular potential generated when the Na^+^ channels open will reach the the neighboring cell when the cells are sufficiently close.

We also observe that the magnitude of the extracellular potential increases when the size of the Na^+^ channel cluster is increased. For *L_e_* = 30 nm, the most negative extracellular potential goes from about −0.5 mV for a cluster of 4 Na^+^ channels to about −15 mV for 900 channels, and for *L_e_* = 5 nm, the most negative potential goes from about −0.7 mV for 4 Na^+^ channels to about −30 mV for 900 channels.

**Figure 9:**
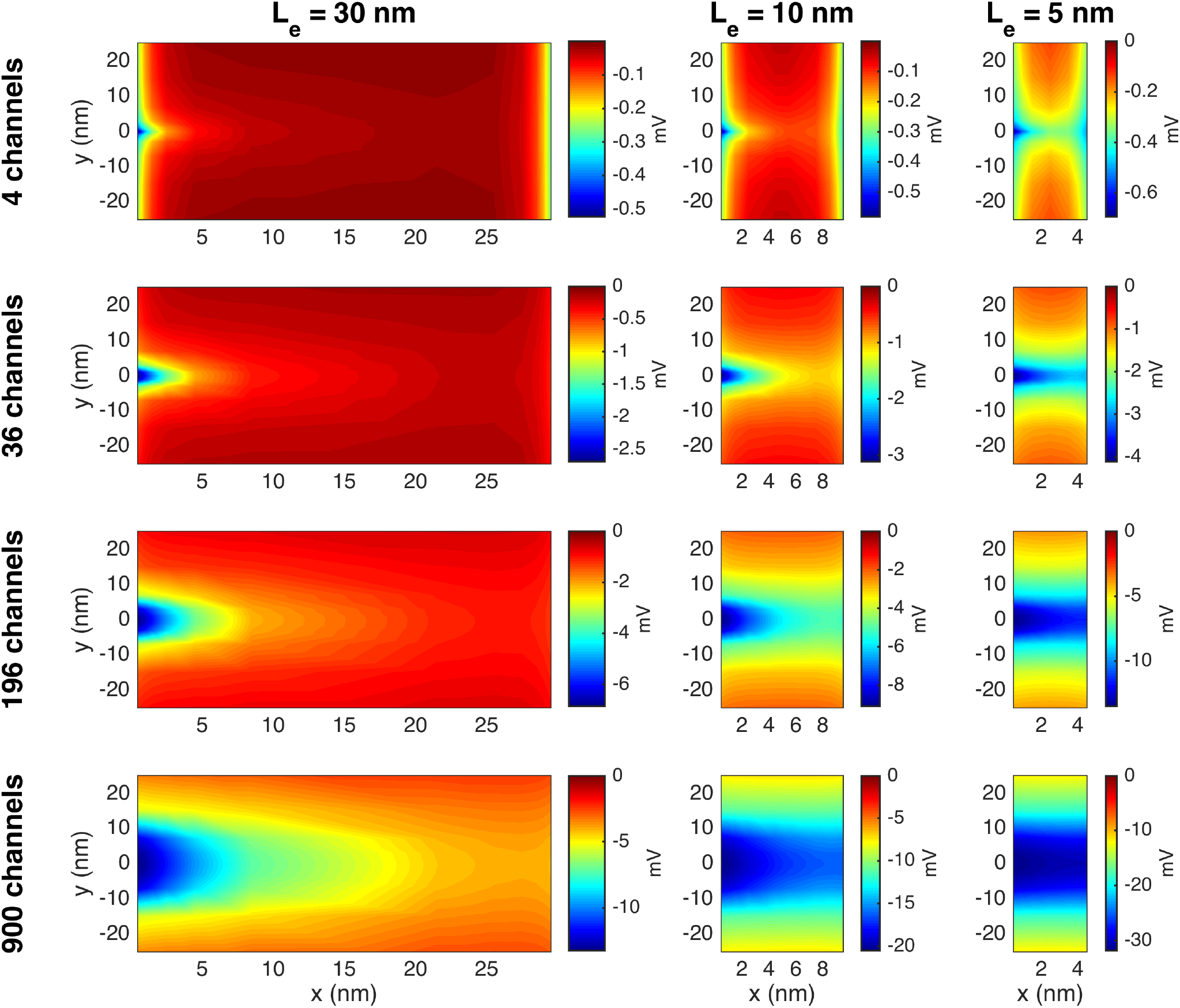
The potential, *ϕ*, in the extracellular space between the two cells in simulations with open Na^+^ channel clusters on the membrane of the left cell. The width of the extracellular space, *L_e_*, and the size of the Na^+^ channel cluster is varied in the columns and rows of the figure, respectively. The plots show the solution in the (*x, y*)-plane for the center of the domain in the *z*-direction at the point in time when the deviation from rest is largest. In the *y*-direction, we focus on the 50 nm closest to the Na^+^ channel clusters. The coordinates on the axes are shifted so that *x* = 0 marks the end of the membrane of the left cell and *y* = 0 marks the center of the Na^+^ channel cluster. Note that the scaling of the colorbar is different for the different cases.

Figures 10 and 11 show similar plots of the extracellular Na^+^ and K^+^ concentrations. In Figure 10, we observe that the extracellular Na^+^ concentration is locally decreased outside of the Na^+^ channel cluster. For *L_e_* = 30 nm, this does not seem to affect the Na^+^ concentration outside of the right cell, but for *L_e_* = 5 nm, there seems to be some effect outside of the right cell in the cases of large Na^+^ channel clusters. In Figure 11, we observe that the K^+^ concentration is locally increased outside of the Na^+^ channel cluster. In addition, we observe a general increase in extracellular K^+^ concentration as *L_e_* is decreased. Similar plots for the extracellular Ca^2+^ and Cl^-^ concentrations are found in the Supplementary Information.

**Figure 10:**
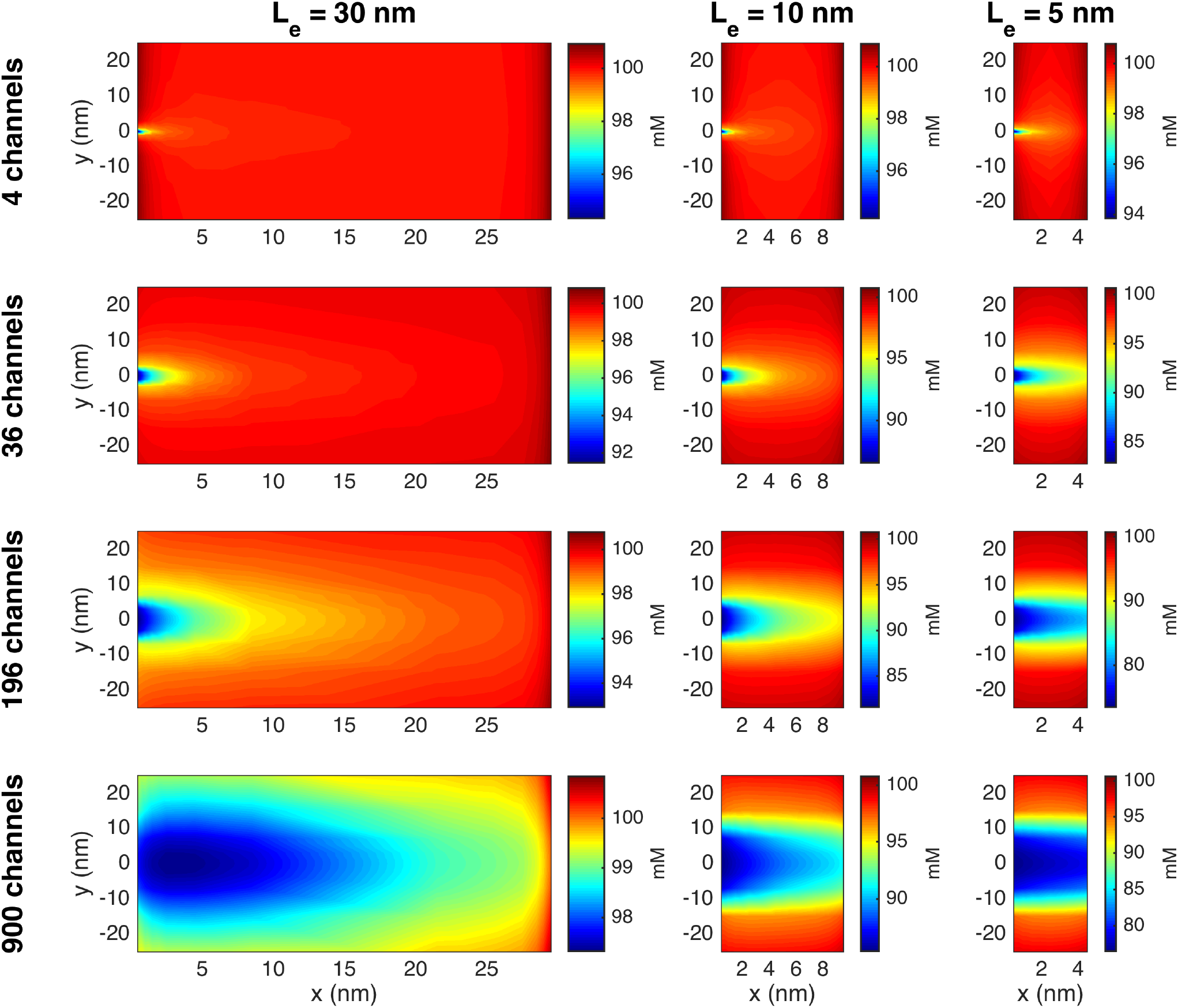
The Na^+^ concentration in the extracellular space between the two cells in simulations with open Na^+^ channel clusters on the membrane of the left cell. The figure setup is the same as for Figure 9.

**Figure 11:**
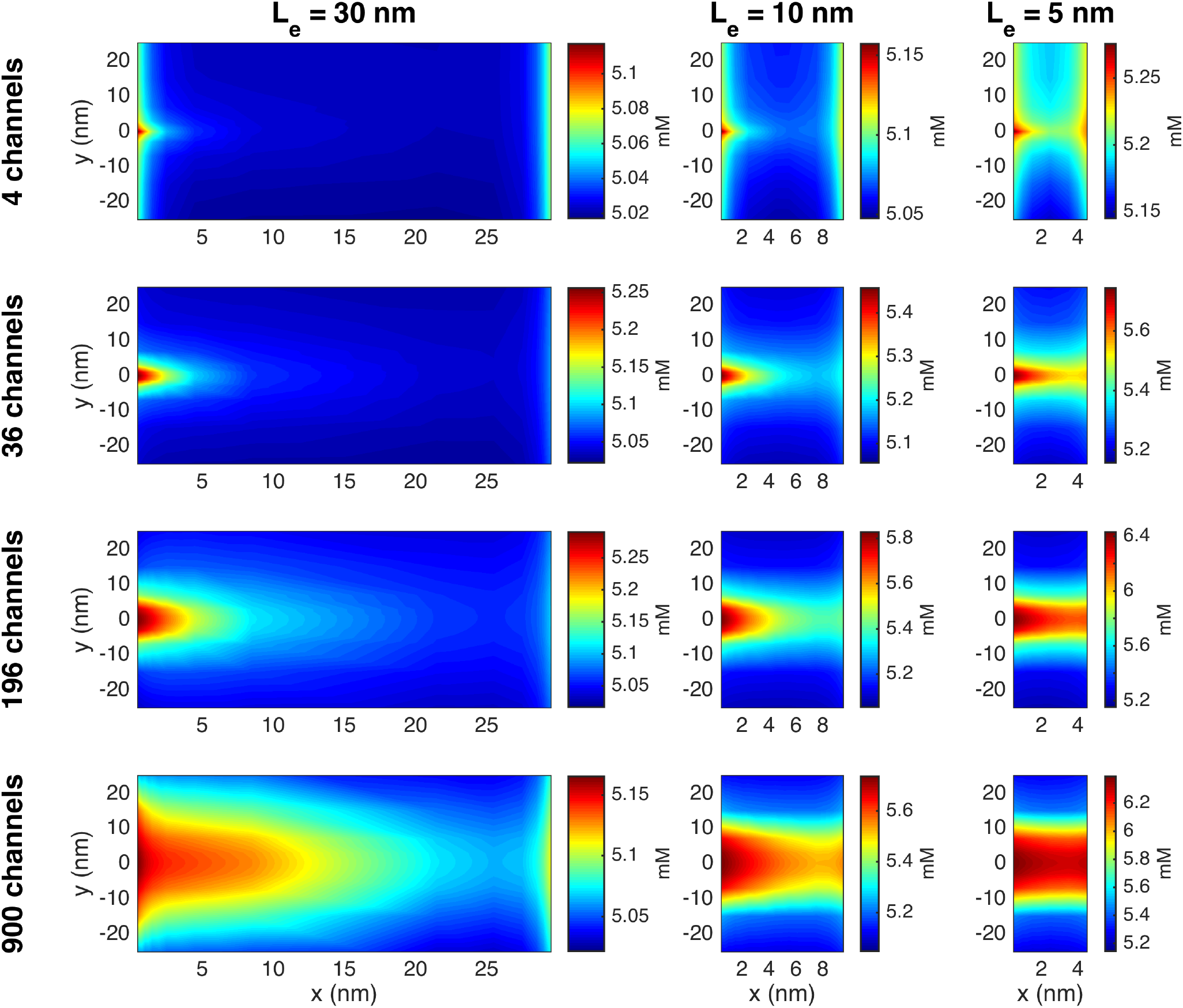
The K^+^ concentration in the extracellular space between the two cells in simulations with open Na^+^ channel clusters on the membrane of the left cell. The figure setup is the same as for Figure 9.

Figure 12 summarizes the results of adjusting the extracellular width and the number of Na^+^ channels on the magnitude of the potential in the extracellular space between the cells. Again, we observe that the magnitude of the negative extracellular potential increases as the distance between the cells (*L_e_*) is decreased. The extracellular potential also increases when the number of Na^+^ channels in the channel clusters increases, and for *L_e_* = 5 nm, the effect is almost identical outside of the right cell as outside of the left cell. For the extracellular widths of *L_e_* = 30 nm and *L_e_* = 10 nm, the extracellular potential is not as negative outside of the right cell as outside of the left cell.

**Figure 12:**
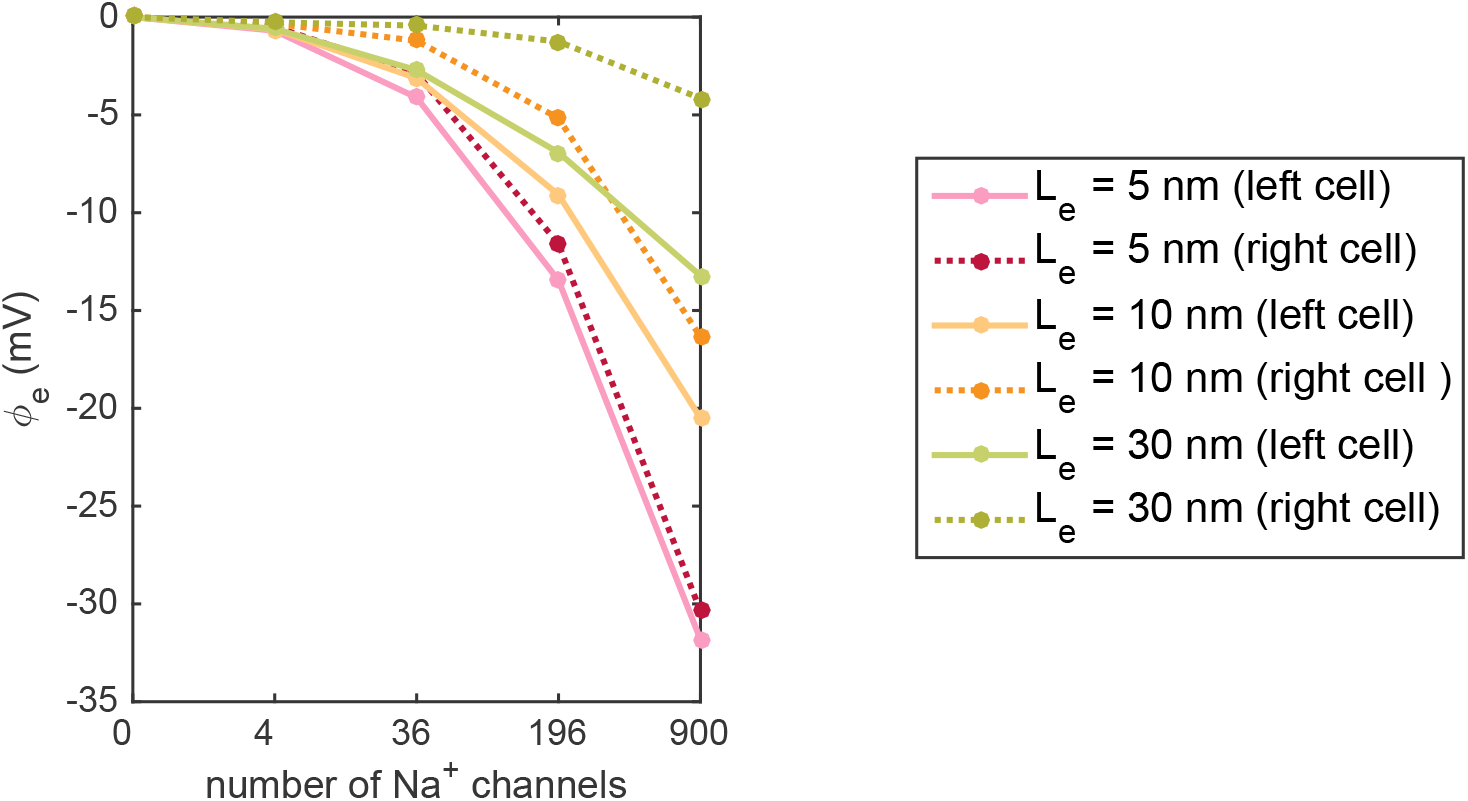
Most negative extracellular potential outside of the Na^+^ channel clusters in simulations with different widths of the extracellular space between the cells, *L_e_*, and different sizes of the Na^+^ channel clusters. The full lines show the solution outside of the Na^+^ channel cluster of the left cell, which has open Na^+^ channels. The dotted lines show the solution on the other side of the extracellular space, outside of the right cell, which has closed Na^+^ channels.

#### 3.2.3 The duration of the electrochemical wave depends on the membrane area included in the simulation

In the simulations reported above, we observe a strong electrochemical wave emanating from the left cell when the Na^+^ channels open. The wave reaches towards the cell on the right-hand side and the strength depends on the number of ion channels available and the width of the extracellular space between the cells (see Figures 9–12). Also, we have seen that the wave is extremely brief. The changes in the potential and the ion concentrations in the simulation reported in Figure 8 last for only about 500 ns. The duration of this wave is critical for understanding ephaptic coupling. If the wave is extremely brief, it is not clear that it will be able to open the Na^+^ channels in the next cell, but if the wave lasts for a sufficiently long time, it may be strong enough to excite the next cell. We therefore need to obtain a firm understanding of the duration of the wave.

Figure 13 shows the membrane potential, the extracellular potential, the extracellular Na^+^ and K^+^ concentrations, the charge density and the total Na^+^ current in simulations with four different values of *L_y_* and *L_z_*. The remaining parameter values are the same in the different simulations. Mimicking inactivation of the sodium channels, the *I*_Na_ channels are forced to close when the membrane potential reaches 30 mV (resulting in the dents in the solutions observed towards the end of the simulations). The scaling of the time axis is different in the different columns of the figure, and we observe that as the membrane area is increased, the duration of the dynamics is increased.

**Figure 13:**
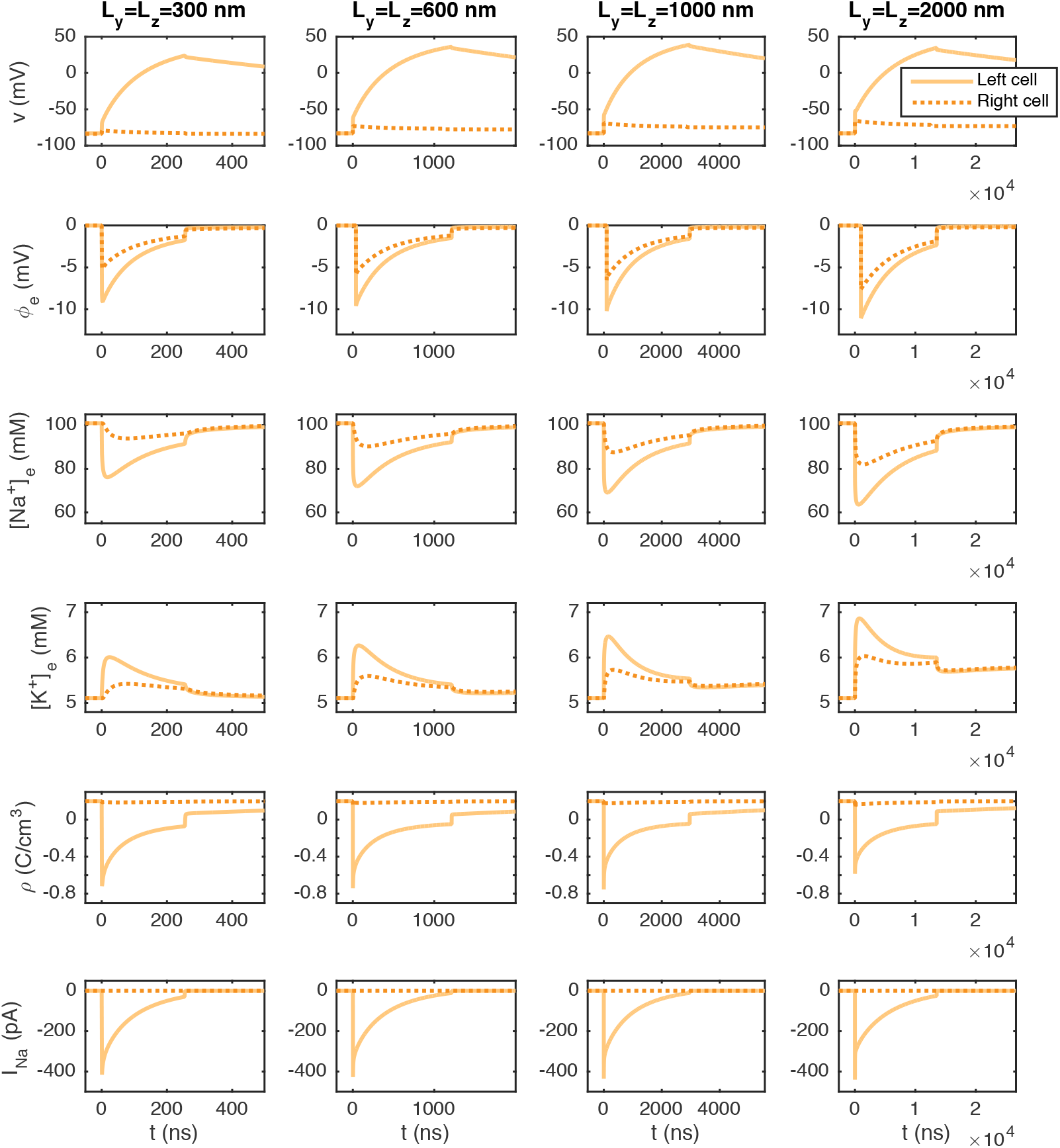
The membrane potential, *v*, the extracellular potential, *ϕ*, the extracellular Na^+^ and K^+^ concentrations, the charge density, ρ, and the total *I*_Na_ current of the left and right cells as functions of time. The membrane potential and *I*_Na_ are measured as described in Figure 5, and *ϕ* and the ionic concentrations are recorded in the first grid point (in the *x*-direction) outside of the center (in the *y*- and *z*-directions) of the Na^+^ channel clusters. In the first column, the total membrane area included for each cell is 300 nm × 300 nm, and in the next columns, the membrane area is 600 nm × 600 nm, 1000 nm × 1000 nm, and 2000 nm × 2000 nm. In the simulations, the cell distance is *L_e_* = 10 nm and the Na^+^ channel cluster consists of 196 Na^+^ channels (see Figure 2) and the remaining parameter values are as specified in Tables 2 and 3. The Na^+^ channels of the left cell are opened at *t* = 0, and the Na^+^ channels of the right are closed in the entire simulations. Note that the scaling of the time axis (*x*-axis) is different in the different columns.

The results of the simulations are summarized in Figure 14. In the left panel, we observe that the upstroke duration, i.e., the time for the membrane potential to increase from the resting potential to 30 mV appears to be proportional to the membrane area included in the simulation. From Figure 13, we also see that this duration seems to correspond to the duration of the change in extracellular potential and concentrations. The magnitude of the *I*_Na_ peak, on the other hand, appears to be very similar for the different membrane areas. In addition, as the membrane area increases, the extracellular potential seems to become a bit more negative, and the minimum value of the Na^+^ concentration decreases and the maximum value of the K^+^ concentration increases.

**Figure 14:**
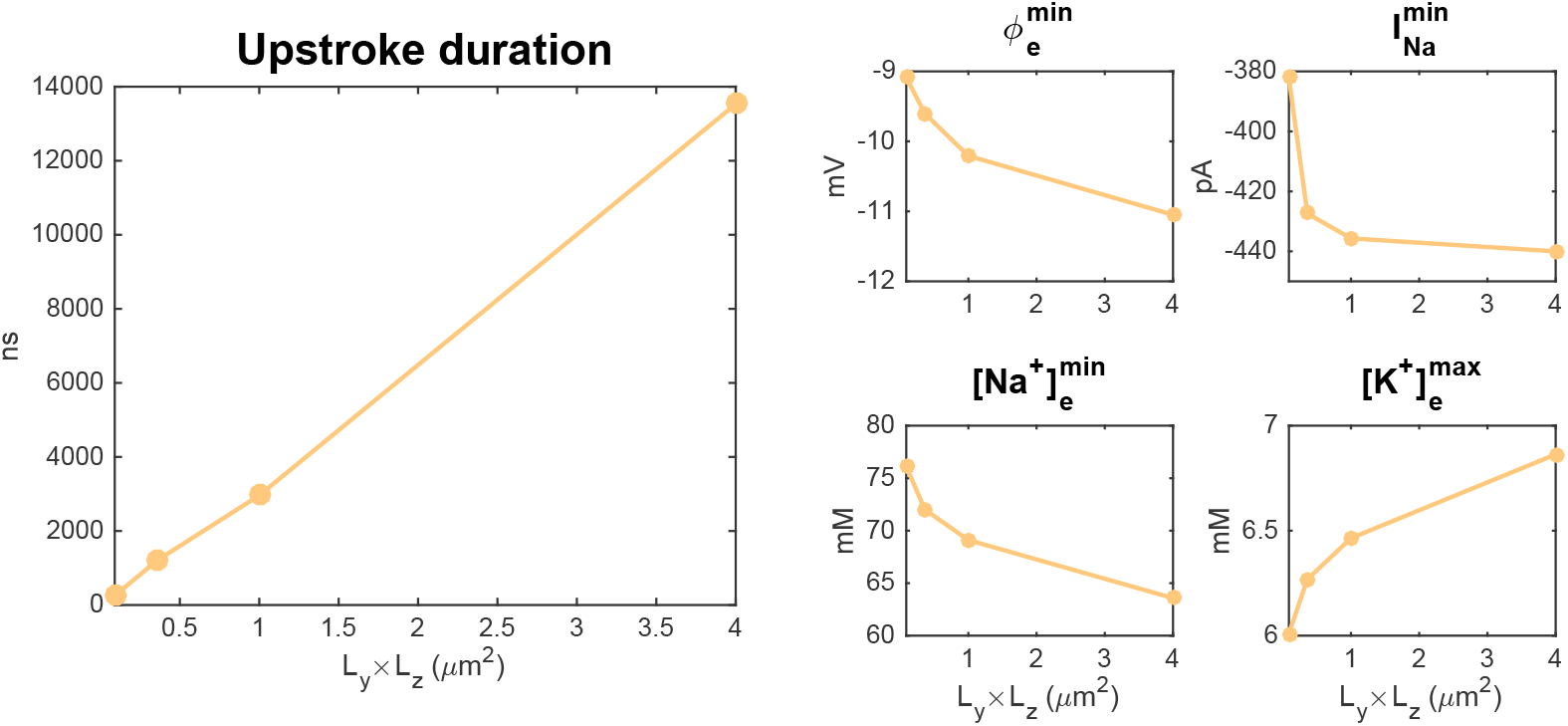
Summary of the results in Figure 13. Left panel: The upstroke duration for the membrane potential of the left cell as a function of the membrane area *L_y_* × *L_z_*. This upstroke duration corresponds to the time for the membrane potential of the left cell to increase from the resting potential to *v* = 30 mV after the *I*_Na_ channel cluster has been opened. Right panels: The minimum value of *ϕ* and [Na^+^] and the maximum value of [K^+^] outside of the Na^+^ channel cluster of the left cell, as well as the minimum value of *I*_Na_ (the I_Na_ peak) of the left cell as functions of the membrane area.

#### 3.2.4 Upstroke duration in a simplified model

To further investigate the relationship between the membrane area included in the simulation and the duration of the upstroke of the membrane potential and the duration of the *I*_Na_ current, we consider the following classical simplified model for how the membrane potential of excitable cells changes with time as a result of the ionic currents through the ion channels of the membrane:

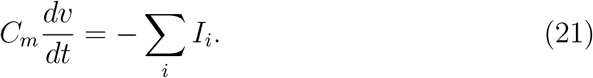

Here, *C_m_* is the capacitance of the membrane (in pF), *v* is the membrane potential (in mV), and *I_i_* are currents (in pA) through different types of membrane ion channels. The capacitance, *C_m_*, can be expressed as

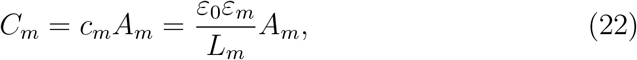

where

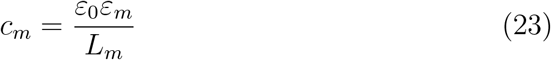

is the specific membrane capacitance (in pF/nm^2^), *ε*_0_ is the vacuum permittivity (in pF/nm), *ε_m_* is the (unitless) relative permittivity of the medium, *L_m_* is the width of the membrane, and *A_m_* is the area of the membrane (in nm^2^) [40]. Furthermore, the current through a type of ion channel can be expressed by the simplified model

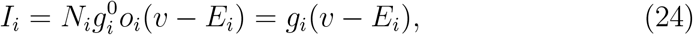

where *N_i_* is the number of ion channels of type *i*, 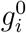 is the conductance through a single open channel of type *i* (in nS), *o_i_* is the open probability of the channels of type *i*, *E_i_* is the Nernst equilibrium potential of the ion type *i*, and

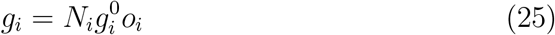

(in nS) is the total conductance of channels of type *i* [41]. In our case, we consider two types of ion currents (*I*_Na_ and *I*_K_), and assume that both types of channels are open (i.e., *o_i_* = 1). Table 4 reports parameterizations of simplified formulations of these currents of the form (24), fitted to the currents observed in the PNP simulations in Figure 13.

**Table 4:** Parameter values for simplified versions of the *I*_Na_ and *I*_K_ currents of the form (24) fitted to the currents in the PNP simulations in Figure 13. In these simulations, *N*_Na_ is 196, *N*_K_ is 36 and *o*_Na_ = *o*_K_ = 1.

| Parameter | Value |
| --- | --- |
| $g_{\text{Na}}^0$ | $20 \cdot 10^{-3} \text{ nS}$ |
| $g_{\text{K}}^0$ | $4.0 \cdot 10^{-3} \text{ nS}$ |
| $E_{\text{Na}}$ | 36 mV |
| $E_{\text{K}}$ | -80 mV |

In the case of the two currents *I*_Na_ and *I*_K_ of the form (24) with o_K_ = *o*_Na_ = 1, the simplified model (21) has the analytical solution

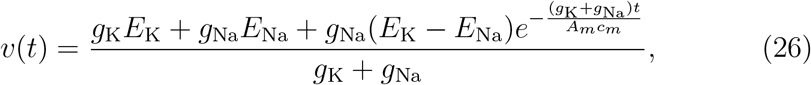

assuming that *v*(0) = *E*_K_. In Figure 15, we compare the analytical solution of the simplified model to the solution of the PNP model for the simulations considered in Figure 13. We observe that the simplified model seems to be a relatively good approximation for the PNP solution for different sizes of the membrane area.

**Figure 15:**
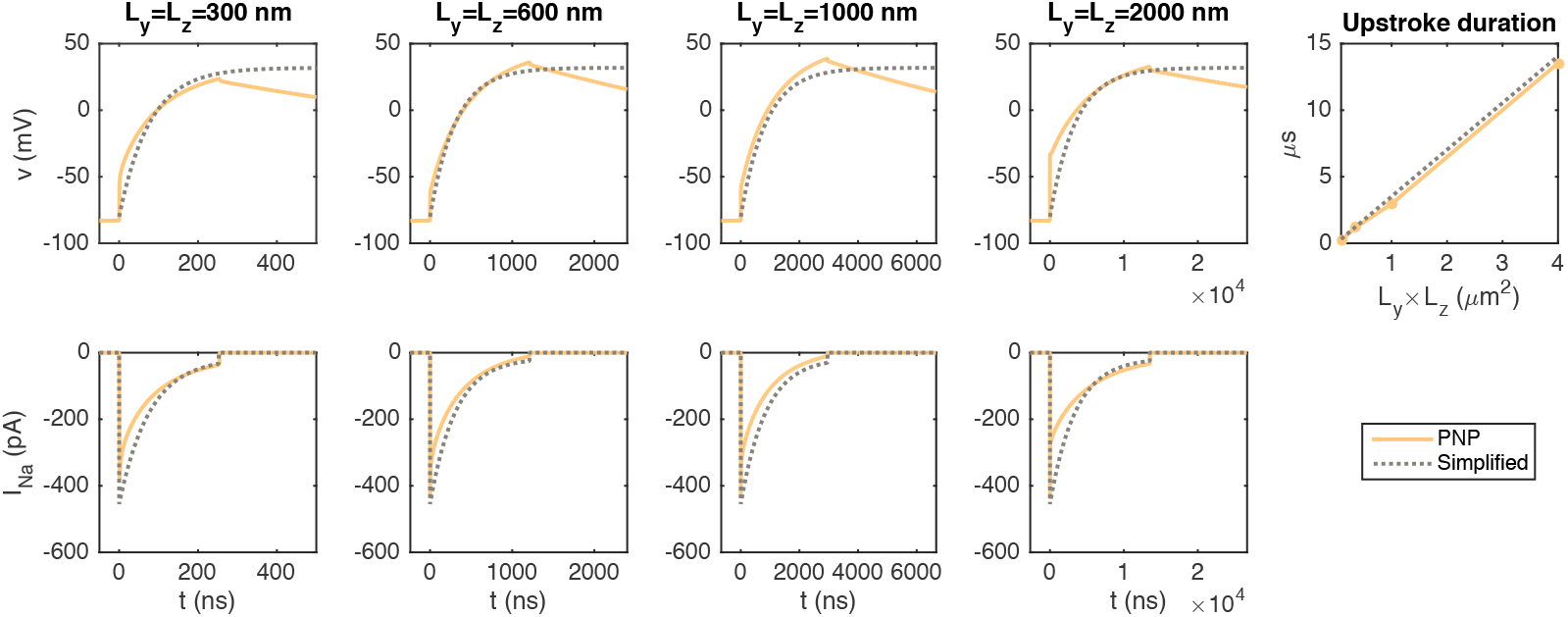
The membrane potential, *v*, and the total *I*_Na_ current of the left cell as functions of time in the PNP simulations reported in Figure 13 and in the solution of the simplified model (21)–(26). The Na^+^ channel cluster consists of 196 open Na^+^ channels. In the PNP simulations, the membrane potential and *I*_Na_ are measured as described in Figure 5, the cell distance is *L_e_* = 10 nm, and the remaining parameter values are as specified in Tables 2 and 3. In the simplified model, the parameter values are as specified in Tables 2, 3 and 4. Note that the scaling of the time axis (*x*-axis) is different in the different columns. The rightmost column reports the time for the membrane potential to increase from the resting potential to *v* = 30 mV as a function of the membrane area in the PNP simulations and as computed by the analytical formula (27).

From the formula (26), the duration from *t* = 0 (i.e., when the Na^+^ channels open) to when the membrane potential has reached the value 30 mV is given by

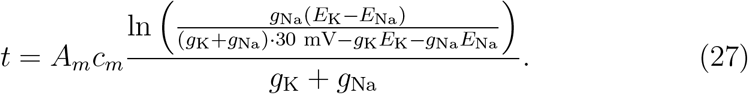

In this formula, it is clear that the time for the membrane potential to reach 30 mV depends linearly of the membrane area included, *A_m_*. In the rightmost panel of Figure 15, we observe that this approximation of the duration of the upstroke is in very good agreement with the durations observed in the PNP simulations.

#### 3.2.5 The extracellular wave will last for about 0.4 ms

In the simulations reported in Figure 13, we consider 196 open Na^+^ channels, and obtain a peak total *I*_Na_ current of about −400 pA. For a membrane area of 300 nm × 300 nm (like in Figure 8), the peak current density for the *I*_Na_ current is 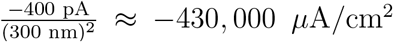. For comparison, in typical models of the cardiac action potential, the peak current density is about −400 *μ*A/cm^2^ (see, e.g., [41, 42]). This indicates that the Na^+^ channel cluster of 196 channels should have been associated with a membrane area that is about 1000 times larger than the 300 nm × 300 nm area used here in order to represent a realistic duration of the dynamics. For example, an area of 10000 nm × 10000 nm could be suitable. We are not, however, currently able to perform such large simulations due to the computational load of running long simulations.

On the other hand, since the simplified model (22) provided results that almost coincide with the PNP results, and the linear relation between the area of the membrane and the duration of the wave is approximated very well, we can use the simplified model to estimate the duration of the wave for any membrane area. By using (27) and the parameters given in Table 4, we find that the wave will last for about 0.35 ms for the 10000 nm × 10000 nm membrane area. This is most likely sufficient to initiate a depolarization of the next cell when the wave reaches across the extracellular space. Adjusting the membrane area similarly to obtain a current density of about −400 *μ*A/cm^2^ for the other Na^+^ channel cluster sizes, this duration will be the same for the different cluster sizes.

We also note that the results in Figures 13 and 14 indicate that the magnitude of the change in the extracellular potential and the ion concentrations increases as the membrane area is increased. This suggests that the changes in potential and concentrations in a simulation with a more realistic membrane area would be at least as large as those observed in our small simulations with a membrane area of 300 nm × 300 nm (see Figures 9–12). Nevertheless, this estimate is associated with a level of uncertainty, since the membrane areas currently investigated are considerably smaller than the membrane area needed for realistic durations of the dynamics.

### 3.3 The gradients of the solutions of the PNP equations

As mentioned above, the gradients involved in the solution of the full PNP model are substantial. The maxmimum gradients in an example simulation are listed in Table 5. These gradients imply that extremely fines meshes and time steps must be applied and this may be the reason why the model is often simplified by assuming electroneutrality. Numerical instabilities arise if the spatial or temporal resolution are too coarse, see, e.g., [11].

**Table 5:** The maximum of the absolute value of the derivative of the potential and the concentrations with respect to time and with respect to *x* in the PNP simulation reported in Figure 8. In this simulation, the K^+^ channels are open for both cells, and a Na^+^ channel cluster consisting of 196 channels are opened for the left cell. The extracellular space width is *L_e_* = 10 nm. As a comparison, the transmembrane electric field is in the range of 15 mV/nm for a membrane potential of −90 mV.

| | $\max \left\vert \frac{\partial}{\partial t} \right\vert $ | $\max \left\vert \frac{\partial}{\partial x} \right\vert $ |
| --- | --- | --- |
| $\phi$ | 26 mV/ns | 51 mV/nm |
| $c_{\text{Na}^+}$ | 13 mM/ns | 22 mM/nm |
| $c_{\text{K}^+}$ | 0.3 mM/ns | 28 mM/nm |
| $c_{\text{Ca}^{2+}}$ | 0.05 mM/ns | 0.06 mM/nm |
| $c_{\text{Cl}^-}$ | 5.5 mM/ns | 6 mM/nm |

## 4 Discussion

Using the Poisson-Nernst-Planck equations, we developed a computational modelling framework to investigate electric potentials and ion concentration changes in the vicinity of ion channel clusters in two opposing cell membranes, as found, e.g., in cardiac intercalated discs. For the present study, we focused on Na^+^ and K^+^ channel clusters that vary in size, corresponding to different numbers of channels in the clusters. To develop the model and solve the mathematical problem, a very fine temporal and spatial discretization was required, leading to a very large computational effort. We limited the computational expenses by reducing the represented membrane areas and modelling only a small part of the intercalated disc membranes. To additionally reduce the number of nodes, an adaptive mesh was used for spatial discretization.

When modelling complex dynamical processes in a small part of two adjacent cells, the choice of appropriate boundary conditions is challenging and of great importance. Therefore, we have tested several alternatives before defining the most suitable boundary conditions for our problem. If only no-flux Neumann boundary conditions are applied on intracellular and extracellular domains, neither current nor ions can enter or exit the modelled region. It is worth noting that Neumann boundary conditions are compatible with the Poisson equation, but corresponds to an unrealistic sealing off of the extracellular intercalated disc domain from the extracellular bulk space. With the aim to simulate the junction between the intercalated disc and bulk space, a Dirichlet condition for ion concentrations and zero potential was therefore applied at the periphery of the extracellular domain. To ensure the formation of the Debye layers close to the cell membranes at the periphery of the simulated extra- and intracellular domains, this Dirichlet condition was applied only in the central part of the extracellular boundary surface (see Figure 3). Given that only a small part of the intercalated disc was modeled, a combined choice of Dirichlet and Neumann boundary conditions is, in our opinion, the most appropriate approach to realistically represent cardiomyocytes joined by an intercalated disc.

### 4.1 Main results

The first main finding in our simulations is the clear and distinct formation of boundary layers, known as Debye layers, for the extracellular potential, the charge density, *ρ*, and the concentrations of Na^+^, K^+^, Ca^2+^ and Cl^-^ when, in a first step, only K^+^ channels are open and the resting state of the model is reached. These Debye layers arise naturally from Gauss’ law of electrostatics [43] and the dielectric property of the membrane. For the small membrane area applied in our simulations, these Debye layers form at the microsecond timescale (see Figure S1 in the Supplementary Information). However, the time scale associated with the formation of the Debye layers may depend on the membrane area associated with the K^+^ channels (as observed for the duration of the dynamics following Na^+^ channel opening in Section 3.2.3; Figure 13.)

Our second main finding was obtained in the second part of the study, when the Na^+^ channels in the cluster were opened. The simulations show that near the opening Na^+^ channel cluster, large concentration changes occur. When Na^+^ ions enter the cytoplasm through open Na^+^ channels, the extracellular Na^+^ concentration at the inlet of the channel cluster decreases. With the large extracellular Na^+^ depletion resulting from the influx of Na^+^ into the left (pre-junctional) cell, a strong negative extracellular potential builds up in the extracellular cleft. Concomitantly, in the extracellular domain, K^+^ ions and Ca^2+^ ions are attracted by the electrochemical gradient towards the Na^+^ channel cluster, while Cl^-^ anions are repelled away from it. Indeed, near the extracellular side of the Na^+^ channel cluster, [K^+^] and [Ca^2+^] transiently rise while [Cl^-^] decreases (see Figure 8). Thus, substantial concentration gradients are generated. In the simulation shown in Figure 8, the large ion concentration gradients dissipate quickly and almost completely on a time scale of ~500 ns, however this time scale appears to be proportional to the membrane area associated with the Na^+^ channel cluster (see Figures 13–15), and a more physiologically realistic duration of the dynamics is estimated to be about 0.4 ms (see Section 3.2.5). More specifically, the changes in the extracellular potential, the charge density and the ionic concentrations in the cleft all seem to last for about as long as the duration of *I*_Na_ current, which self-limitates as the membrane potential, *v*, approaches the Nernst equilibrium potential for Na^+^, E_Na_ (see Section 3.2.4).

It should be noted that even though the charge density, *ρ*, is non-zero close to the membrane in our simulations, and in particular close to the open Na^+^ channel cluster during the upstroke of the membrane potential (see, e.g., Figure 6 and Figure 8), we believe that for larger scale models (e.g., individual cardiomyocytes), electroneutrality is a reasonable and necessary assumption to reduce the large computational effort, as was done in other modeling frameworks examining biological membranes and channels, such as the Kirchhoff-Nernst-Planck formalism [9, 12].

Our third main finding is that an increasing number of Na^+^ channels in a cluster and reduced extracellular cleft width lead to more negative extracellular potentials in the cleft. The number of Na^+^ channels was modelled by an increased Na^+^ channel cluster size. The question arises whether the size or the number of Na^+^ channels leads to the more negative extracellular potential in the cleft. According to simulations with the computational model of the intercalated disc by Hichri et al. [21], when the number of Na^+^ channels at the intercalated disc membrane remains constant but the cluster size is decreased, the extracellular potential becomes more negative and thus ephaptic effects are accentuated. Therefore, we conclude that the number of Na^+^ channels rather than the cluster size is the key factor in our simulation results. Moreover, the reduction of the extracellular cleft width (from 30 nm to 5 nm), leading to more negative extracellular potentials in the cleft is consistent with previously published computational studies at the cellular level [9, 21, 3]. With decreasing cleft width, negative extracellular potentials in the local neighborhood of the Na^+^ channel cluster of the left cell are transmitted to the other side of the cleft (close to the right cell), where it is likely to affect Na^+^ channel gating, thus forming the basis for ephaptic coupling.

### 4.2 Limitations and perspectives

The strength of our modelling framework is the full incorporation of the Poisson-Nernst-Planck equations at the level of ion channel clusters. However, during the development of the model, some highly simplifying assumptions were necessary.

A first simplifying assumption was taken at the level of the ion channel pores, in terms of the free energy profile that a permeating ion is subjected to as it passes through a channel. This energy profile is determined by the distribution of charges within the protein forming the channel, especially in the selectivity filter, which is typically lined up with amino acid residues bearing a charge that is opposite to that of the permeating ion [44]. As the background charge density *ρ*_0_ of ion channels (e.g., Na^+^ and K^+^ channels) is not precisely known, we assumed for our main results a linear *ρ*_0_ profile from the intracellular to the extracellular part of the Na^+^ and K^+^ channels. However, as shown in the supplementary materials, different concentration and *ρ*_0_ profiles (i.e., an abrupt discontinuity in the center of the channel) affect permeation [44], and thus the simulation results. Moreover, ion channel permeation kinetics are strongly influenced by the dehydration and rehydration of permeating ions inside the channels, which further affect the free energy profile [44]. We did not incorporate these channel features into our model. Another aspect that we did not consider in our framework is the negative charge at the surface of the phospholipid bilayer membrane [44], that may have an additional effect on the simulation results. However, the incorporation of such details into the model was not in the scope of our study. A full model integrating ion channel permeation kinetics would require a mixed framework with molecular dynamics of channel permeation, which would add complexity and computational effort to our already computationally intensive modelling framework.

Because our simulations corresponded to nano- and microsecond time scales, we also assumed in our model that clustered ion channels open at the same time, while in reality they exhibit asynchronous stochastic openings and closings resulting from voltage gating. However, for Na^+^ channels, this gating occurs at the much longer millisecond scale. In future work, we plan to incorporate the gating of channels, which will allow the examination of action potential propagation via ephaptic coupling. The incorporation of channel gating will provide insight into possible cooperative gating within the channel clusters [45, 46].

## 5 Conclusion

In conclusion, our new computational model allows a detailed investigation of dynamic ion concentration changes around single Na^+^ channel and K^+^ channel clusters in a small fraction of two opposing intercalated disc membranes. The features of the full Poisson-Nernst-Planck modelling framework can be in the future incorporated into larger scale models. Our simulation results highlight the importance of incorporating dynamic ion concentration changes into existing models to obtain a full picture of the interaction between ion channels, ion concentrations, and electric potentials at the spatial and temporal nanoscales. Particularly, this will be crucial to gain further insights into the mechanisms of ephaptic coupling as a complementary mechanism to gap junctional coupling for action potential propagation in the heart. This knowledge may be useful, in the future, to develop more suitable treatments for cardiac arrhythmias.

## References

[1] Piero C Franzone, Luca F Pavarino, and Simone Scacchi. Mathematical Cardiac Electrophysiology, volume 13. Springer, 2014.

[2] Aslak Tveito, Karoline H Jæger, Miroslav Kuchta, Kent-Andre Mardal, and Marie E Rognes. A cell-based framework for numerical modeling of electrical conduction in cardiac tissue. Frontiers in Physics, 5:48, 2017.

[3] Ena Ivanovic and Jan P Kucera. Localization of Na+ channel clusters in narrowed perinexi of gap junctions enhances cardiac impulse transmission via ephaptic coupling: a model study. The Journal of Physiology, 599(21):4779–4811, 2021.

[4] Sebastián Domínguez, Joyce Reimer, Kevin R Green, Reza Zolfaghari, and Raymond J Spiteri. A simulation-based method to study the lqt1 syndrome remotely using the emi model. In Emerging Technologies in Biomedical Engineering and Sustainable TeleMedicine, pages 179–189. Springer, 2021.

[5] Karoline Horgmo Jæger and Aslak Tveito. Deriving the bidomain model of cardiac electrophysiology from a cell-based model; properties and comparisons. Frontiers in Physiology, page 2439, 2022.

[6] Karoline Horgmo Jæger, Andrew G Edwards, Wayne R Giles, and Aslak Tveito. From millimeters to micrometers; re-introducing myocytes in models of cardiac electrophysiology. Frontiers in Physiology, 12, 2021.

[7] Carl L Gardner and Jeremiah R Jones. Electrodiffusion model simulation of the potassium channel. Journal of Theoretical Biology, 291:10–13, 2011.

[8] Jerzy J Jasielec. Electrodiffusion phenomena in neuroscience and the nernst–planck–poisson equations. Electrochem, 2(2):197–215, 2021.

[9] Yoichiro Mori, Glenn I Fishman, and Charles S Peskin. Ephaptic conduction in a cardiac strand model with 3d electrodiffusion. Proceedings of the National Academy of Sciences, 105(17):6463–6468, 2008.

[10] Yoichiro Mori and Charles Peskin. A numerical method for cellular electrophysiology based on the electrodiffusion equations with internal boundary conditions at membranes. Communications in Applied Mathematics and Computational Science, 4(1):85–134, 2009.

[11] Andreas Solbrå, Aslak Wigdahl Bergersen, Jonas van den Brink, Anders Malthe-Sørenssen, Gaute T Einevoll, and Geir Halnes. A Kirchhoff-Nernst-Planck framework for modeling large scale extracellular electrodiffusion surrounding morphologically detailed neurons. PLoS Computational Biology, 14(10):e1006510, 2018.

[12] Ada J Ellingsrud, Andreas Solbrå, Gaute T Einevoll, Geir Halnes, and Marie E Rognes. Finite element simulation of ionic electrodiffusion in cellular geometries. Frontiers in Neuroinformatics, 14:11, 2020.

[13] Ada J Ellingsrud, Cécile Daversin-Catty, and Marie E Rognes. A cell-based model for ionic electrodiffusion in excitable tissue. In Modeling Excitable Tissue, pages 14–27. Springer, Cham, 2021.

[14] Jurgis Pods. A comparison of computational models for the extracellular potential of neurons. Journal of Integrative Neuroscience, 16(1):19–32, 2017.

[15] Edmund JF Dickinson, Juan G Limon-Petersen, and Richard G Compton. The electroneutrality approximation in electrochemistry. Journal of Solid State Electrochemistry, 15(7):1335–1345, 2011.

[16] Rengasayee Veeraraghavan, Robert G Gourdie, and Steven Poelzing. Mechanisms of cardiac conduction: a history of revisions. American Journal of Physiology-Heart and Circulatory Physiology, 306(5):H619–H627, 2014.

[17] André G Kléber and Yoram Rudy. Basic mechanisms of cardiac impulse propagation and associated arrhythmias. Physiological reviews, 84(2):431–488, 2004.

[18] Nick Sperelakis and James E Mann Jr. Evaluation of electric field changes in the cleft between excitable cells. Journal of theoretical biology, 64(1):71–96, 1977.

[19] M Suenson. Ephaptic impulse transmission between ventricular myocardial cells in vitro. Acta Physiologica Scandinavica, 120(3):445–455, 1984.

[20] Seth H Weinberg. Ephaptic coupling rescues conduction failure in weakly coupled cardiac tissue with voltage-gated gap junctions. Chaos: An Interdisciplinary Journal of Nonlinear Science, 27(9):093908, 2017.

[21] Echrak Hichri, Hugues Abriel, and Jan P Kucera. Distribution of cardiac sodium channels in clusters potentiates ephaptic interactions in the intercalated disc. The Journal of Physiology, 596(4):563–589, 2018.

[22] Karoline Horgmo Jæger, Andrew G Edwards, Andrew McCulloch, and Aslak Tveito. Properties of cardiac conduction in a cell-based computational model. PLoS Computational Biology, 15(5):e1007042, 2019.

[23] Jonathan Bell. Modeling parallel, unmyelinated axons: Pulse trapping and ephaptic transmission. SIAM Journal on Applied Mathematics, 41(1):168–180, 1981.

[24] Gary R Holt and Christof Koch. Electrical interactions via the extracellular potential near cell bodies. Journal of Computational Neuroscience, 6(2):169–184, 1999.

[25] Costas A Anastassiou and Christof Koch. Ephaptic coupling to endogenous electric field activity: why bother? Current Opinion in Neurobiology, 31:95–103, 2015.

[26] Costas A Anastassiou, Rodrigo Perin, Henry Markram, and Christof Koch. Ephaptic coupling of cortical neurons. Nature Neuroscience, 14(2):217–223, 2011.

[27] Aslak Tveito, Karoline H Jæger, Glenn T Lines, Lukasz Paszkowski, Joakim Sundnes, Andrew G Edwards, Tuomo Māki-Marttunen, Geir Halnes, and Gaute T Einevoll. An evaluation of the accuracy of classical models for computing the membrane potential and extracellular potential for neurons. Frontiers in Computational Neuroscience, 11:27, 2017.

[28] Alessio Paolo Buccino, Miroslav Kuchta, Karoline Horgmo Jæger, Torbjørn Vefferstad Ness, Pierre Berthet, Kent-Andre Mardal, Gert Cauwenberghs, and Aslak Tveito. How does the presence of neural probes affect extracellular potentials? Journal of Neural Engineering, 16(2):026030, 2019.

[29] Yong Wang, Yanxin Liu, Hannah A DeBerg, Takeshi Nomura, Melinda Tonks Hoffman, Paul R Rohde, Klaus Schulten, Boris Martinac, and Paul R Selvin. Single molecule FRET reveals pore size and opening mechanism of a mechano-sensitive ion channel. Elife, 3:e01834, 2014.

[30] Peter J. Mohr, David B. Newell, Barry N. Taylor, and E. Tiesinga. NIST reference on constants, units, and uncertainty. https://physics.nist.gov/cuu/Constants/index.html, Fundamental Constants Data Center of the NIST Physical Measurement Laboratory, 2018. Accessed: 2022-04-03.

[31] Maria G Kurnikova, Rob D Coalson, Peter Graf, and Abraham Nitzan. A lattice relaxation algorithm for three-dimensional poisson-nernst-planck theory with application to ion transport through the gramicidin a channel. Biophysical Journal, 76(2):642–656, 1999.

[32] Jérôme Clatot, Malcolm Hoshi, Xiaoping Wan, Haiyan Liu, Ankur Jain, Krekwit Shinlapawittayatorn, Céline Marionneau, Eckhard Ficker, Taekjip Ha, and Isabelle Deschênes. Voltage-gated sodium channels assemble and gate as dimers. Nature Communications, 8(1):1–14, 2017.

[33] Mathieu Lemay, Enno de Lange, and Jan P Kucera. Effects of stochastic channel gating and distribution on the cardiac action potential. Journal of Theoretical Biology, 281(1):84–96, 2011.

[34] Alejandra Leo-Macias, Esperanza Agullo-Pascual, Jose L Sanchez-Alonso, Sarah Keegan, Xianming Lin, Tatiana Arcos, Yuri E Korchev, Julia Gorelik, David Fenyö, Eli Rothenberg, et al. Nanoscale visualization of functional adhesion/excitability nodes at the intercalated disc. Nature Communications, 7(1):1–12, 2016.

[35] Sarah H Vermij, Jean-Sébastien Rougier, Esperanza Agulleó-Pascual, Eli Rothenberg, Mario Delmar, and Hugues Abriel. Single-molecule localization of the cardiac voltage-gated sodium channel reveals different modes of reorganization at cardiomyocyte membrane domains. Circulation: Arrhythmia and Electrophysiology, 13(7):e008241, 2020.

[36] Rengasayee Veeraraghavan, Gregory S Hoeker, Anita Alvarez-Laviada, Daniel Hoagland, Xiaoping Wan, D Ryan King, Jose Sanchez-Alonso, Chunling Chen, Jane Jourdan, Lori L Isom, et al. The adhesion function of the sodium channel beta subunit (β1) contributes to cardiac action potential propagation. Elife, 7:e37610, 2018.

[37] Dongdong He, Kejia Pan, and Xiaoqiang Yue. A positivity preserving and free energy dissipative difference scheme for the poisson–nernst–planck system. Journal of Scientific Computing, 81(1):436–458, 2019.

[38] Jingwei Hu and Xiaodong Huang. A fully discrete positivity-preserving and energy-dissipative finite difference scheme for poisson–nernst–planck equations. Numerische Mathematik, 145(1):77–115, 2020.

[39] A Tricot, IM Sokolov, and D Holcman. Modeling the voltage distribution in a non-locally but globally electroneutral confined electrolyte medium: applications for nanophysiology. Journal of Mathematical Biology, 82(7):1–31, 2021.

[40] Karoline Horgmo Jæger and Aslak Tveito. Derivation of a cell-based mathematical model of excitable cells. In Modeling Excitable Tissue, pages 1–13. Springer, Cham, 2021.

[41] Karoline Horgmo Jæger, Verena Charwat, Bérénice Charrez, Henrik Finsberg, Mary M Maleckar, Samuel Wall, Kevin E Healy, and Aslak Tveito. Improved computational identification of drug response using optical measurements of human stem cell derived cardiomyocytes in microphysiological systems. Frontiers in Pharmacology, 10:1648, 2020.

[42] Eleonora Grandi, Francesco S Pasqualini, and Donald M Bers. A novel computational model of the human ventricular action potential and ca transient. Journal of Molecular and Cellular Cardiology, 48(1):112–121, 2010.

[43] John David Jackson. Classical electrodynamics. Wiley, New York, 1998. 3rd edition.

[44] Bertil Hille. Ion channels of excitable membranes. Sinauer Associates, Sunderland, MA, U.S.A, 1992.

[45] Kee-Hyun Choi. Cooperative gating between ion channels. General Physiology and Biophysics, 33(1):1–12, 2013.

[46] Rose E Dixon, Manuel F Navedo, Marc D Binder, and L Fernando Santana. Mechanisms and physiological implications of cooperative gating of clustered ion channels. Physiological Reviews, 102(3):1159–1210, 2022.

