## Supplementary Information for "Nano-scale solution of the Poisson-Nernst-Planck (PNP) equations in a fraction of two neighboring cells reveals the magnitude of intercellular electrochemical waves"

September 7, 2022

**S2 Temporal development from electroneutrality to the resting state** **4**

**S3 Profile of $\rho$ in the $K^+$ channel cluster at rest** **6**

**S4 $Ca^{2+}$ and $Cl^-$ concentrations when $Na^+$ channels are open** **7**

**S5 Different choice of $\rho_0$ in the channels** **7**

**S6 A different set of boundary conditions** **10**

### S1 Numerical scheme

In this section, we describe the numerical scheme used to solve the PNP model equations in three dimensions.

#### S1.1 Temporal splitting scheme

For every timestep $n$, we assume that the concentrations $c_k^{n-1}$ (and consequently $\rho^{n-1}$) are known for $t_{n-1}$, and we solve the PNP system in two steps:

**Step 1:** Solve

$$\nabla_h \cdot (\varepsilon \nabla_h \phi^n) = -\rho^{n-1} \tag{1}$$

**Step 2:** Solve

$$\frac{c_k^n - c_k^{n-1}}{\Delta t} = \nabla_h \cdot D_k \nabla_h c_k^n + \nabla_h \cdot \left( D_k \beta_k c_k^n \nabla_h \phi^n \right), \tag{2}$$

$$\text{for } k = \{\mathrm{Na}^+,\ \mathrm{K}^+,\ \mathrm{Ca}^{2+},\ \mathrm{Cl}^-\},$$

where $\nabla_h$ refers to a finite difference discretization of $\nabla$.

We consider the solution in a 3D grid of $N_x \times N_y \times N_z$ points, at $N_x$ positions in the $x$-direction, $N_y$ positions in the $y$-direction and $N_z$ positions in the $z$-direction. The distance between the points in the $x$-direction are denoted by $\Delta x_1$, ..., $\Delta x_{N_x-1}$, and the distance between the points in the $y$- and $z$-directions are similarly denoted by $\Delta y_1$, ..., $\Delta y_{N_y-1}$ and $\Delta z_1$, ..., $\Delta z_{N_z-1}$, respectively.

#### S1.2 Finite difference scheme for Step 1

The finite difference scheme for (1) is given by

$$\begin{aligned}
&\frac{\varepsilon_{i+1/2,j,q} \frac{\partial}{\partial x} \phi^n_{i+1/2,j,q} - \varepsilon_{i-1/2,j,q} \frac{\partial}{\partial x} \phi^n_{i-1/2,j,q}}{0.5(\Delta x_{i-1} + \Delta x_i)} \\
&+ \frac{\varepsilon_{i,j+1/2,q} \frac{\partial}{\partial y} \phi^n_{i,j+1/2,q} - \varepsilon_{i,j-1/2,q} \frac{\partial}{\partial y} \phi^n_{i,j-1/2,q}}{0.5(\Delta y_{j-1} + \Delta y_j)} \\
&+ \frac{\varepsilon_{i,j,q+1/2} \frac{\partial}{\partial z} \phi^n_{i,j,q+1/2} - \varepsilon_{i,j,q-1/2} \frac{\partial}{\partial z} \phi^n_{i,j,q-1/2}}{0.5(\Delta z_{q-1} + \Delta z_q)} = -\rho^{n-1}_{i,j,q},
\end{aligned}$$

or written out more completely,

$$\frac{\varepsilon_{i+1/2,j,q} \left( \phi^n_{i+1,j,q} - \phi^n_{i,j,q} \right)}{0.5(\Delta x_{i-1} + \Delta x_i)\Delta x_i} + \frac{\varepsilon_{i-1/2,j,q} \left( \phi^n_{i-1,j,q} - \phi^n_{i,j,q} \right)}{0.5(\Delta x_{i-1} + \Delta x_i)\Delta x_{i-1}}$$

$$+\frac{\varepsilon_{i,j+1/2,q}\left(\phi^n_{i,j+1,q}-\phi^n_{i,j,q}\right)}{0.5(\Delta y_{j-1}+\Delta y_j)\Delta y_j}+\frac{\varepsilon_{i,j-1/2,q}\left(\phi^n_{i,j-1,q}-\phi^n_{i,j,q}\right)}{0.5(\Delta y_{j-1}+\Delta y_j)\Delta y_{j-1}}$$
$$+\frac{\varepsilon_{i,j,q+1/2}\left(\phi^n_{i,j,q+1}-\phi^n_{i,j,q}\right)}{0.5(\Delta z_{q-1}+\Delta z_q)\Delta z_q}+\frac{\varepsilon_{i,j,q-1/2}\left(\phi^n_{i,j,q-1}-\phi^n_{i,j,q}\right)}{0.5(\Delta z_{q-1}+\Delta z_q)\Delta z_{q-1}}=-\rho^{n-1}_{i,j,q}.$$

Here, as an example, $\varepsilon_{i-1/2,j,q}$ is the value of $\varepsilon$ evaluated in the point

$$\left(\sum_{m=1}^{i-1}(\Delta x_m)-\Delta x_{i-1}/2,\sum_{m=1}^{j-1}(\Delta y_m),\sum_{m=1}^{q-1}(\Delta z_m)\right).$$

#### S1.3 Finite difference scheme for Step 2

The finite difference scheme follows the same structure for the different ionic species, $k$. In order to avoid confusion related to the index $k$ for the different ionic species, we therefore describe the scheme for an arbitrary concentration $c$, with diffusion coefficient $D$ and $\beta$-value $\beta$, where $c$, $D$ and $\beta$ can be replaced by $c_k$, $D_k$ and $\beta_k$ for any of the ionic species, $k$.

The scheme reads

$$\frac{c^n_{i,j,q}-c^{n-1}_{i,j,q}}{\Delta t}=\frac{D_{i+1/2,j,q}\frac{\partial}{\partial x}c^n_{i+1/2,j,q}-D_{i-1/2,j,q}\frac{\partial}{\partial x}c^n_{i-1/2,j,q}}{0.5(\Delta x_{i-1}+\Delta x_i)}$$
$$+\frac{D_{i,j+1/2,q}\frac{\partial}{\partial y}c^n_{i,j+1/2,q}-D_{i,j-1/2,q}\frac{\partial}{\partial y}c^n_{i,j-1/2,q}}{0.5(\Delta y_{j-1}+\Delta y_j)}$$
$$+\frac{D_{i,j,q+1/2}\frac{\partial}{\partial z}c^n_{i,j,q+1/2}-D_{i,j,q-1/2}\frac{\partial}{\partial z}c^n_{i,j,q-1/2}}{0.5(\Delta z_{q-1}+\Delta z_q)}$$
$$+\frac{D_{i+1/2,j,q}\beta c^n_{i+1/2,j,q}\frac{\partial}{\partial x}\phi^n_{i+1/2,j,q}-D_{i-1/2,j,q}\beta c^n_{i-1/2,j,q}\frac{\partial}{\partial x}\phi^n_{i-1/2,j,q}}{0.5(\Delta x_{i-1}+\Delta x_i)}$$
$$+\frac{D_{i,j+1/2,q}\beta c^n_{i,j+1/2,q}\frac{\partial}{\partial y}\phi^n_{i,j+1/2,q}-D_{i,j-1/2,q}\beta c^n_{i,j-1/2,q}\frac{\partial}{\partial y}\phi^n_{i,j-1/2,q}}{0.5(\Delta y_{j-1}+\Delta y_j)}$$
$$+\frac{D_{i,j,q+1/2}\beta c^n_{i,j,q+1/2}\frac{\partial}{\partial z}\phi^n_{i,j,q+1/2}-D_{i,j,q-1/2}\beta c^n_{i,j,q-1/2}\frac{\partial}{\partial z}\phi^n_{i,j,q-1/2}}{0.5(\Delta z_{q-1}+\Delta z_q)}.$$

Written out more completely, and using approximations of the type $c_{i+1/2,j,q}\approx 0.5(c_{i+1,j,q}+c_{i,j,q})$,

$$\frac{c^n_{i,j,q}-c^{n-1}_{i,j,q}}{\Delta t}=\frac{D_{i+1/2,j,q}\left(c^n_{i+1,j,q}-c^n_{i,j,q}\right)}{0.5(\Delta x_{i-1}+\Delta x_i)\Delta x_i}+\frac{D_{i-1/2,j,q}\left(c^n_{i-1,j,q}-c^n_{i,j,q}\right)}{0.5(\Delta x_{i-1}+\Delta x_i)\Delta x_{i-1}}$$
$$+\frac{D_{i,j+1/2,q}\left(c^n_{i,j+1,q}-c^n_{i,j,q}\right)}{0.5(\Delta y_{j-1}+\Delta y_j)\Delta y_j}+\frac{D_{i,j-1/2,q}\left(c^n_{i,j-1,q}-c^n_{i,j,q}\right)}{0.5(\Delta y_{j-1}+\Delta y_j)\Delta y_{j-1}}$$

$$
\begin{aligned}
&+ \frac{D_{i,j,q+1/2}\left(c^n_{i,j,q+1} - c^n_{i,j,q}\right)}{0.5(\Delta z_{q-1} + \Delta z_q)\Delta z_q} + \frac{D_{i,j,q-1/2}\left(c^n_{i,j,q-1} - c^n_{i,j,q}\right)}{0.5(\Delta z_{q-1} + \Delta z_q)\Delta z_{q-1}} \\
&+ \frac{D_{i+1/2,j,q}\beta\left(c^n_{i+1,j,q} + c^n_{i,j,q}\right)\left(\phi^n_{i+1,j,q} - \phi^n_{i,j,q}\right)}{(\Delta x_{i-1} + \Delta x_i)\Delta x_i} \\
&+ \frac{D_{i-1/2,j,q}\beta\left(c^n_{i-1,j,q} + c^n_{i,j,q}\right)\left(\phi^n_{i-1,j,q} - \phi^n_{i,j,q}\right)}{(\Delta x_{i-1} + \Delta x_i)\Delta x_{i-1}} \\
&+ \frac{D_{i,j+1/2,q}\beta\left(c^n_{i,j+1,q} + c^n_{i,j,q}\right)\left(\phi^n_{i,j+1,q} - \phi^n_{i,j,q}\right)}{(\Delta y_{j-1} + \Delta y_j)\Delta y_j} \\
&+ \frac{D_{i,j-1/2,q}\beta\left(c^n_{i,j-1,q} + c^n_{i,j,q}\right)\left(\phi^n_{i,j-1,q} - \phi^n_{i,j,q}\right)}{(\Delta y_{j-1} + \Delta y_j)\Delta y_{j-1}} \\
&+ \frac{D_{i,j,q+1/2}\beta\left(c^n_{i,j,q+1} + c^n_{i,j,q}\right)\left(\phi^n_{i,j,q+1} - \phi^n_{i,j,q}\right)}{(\Delta z_{q-1} + \Delta z_q)\Delta z_q} \\
&+ \frac{D_{i,j,q-1/2}\beta\left(c^n_{i,j,q-1} + c^n_{i,j,q}\right)\left(\phi^n_{i,j,q-1} - \phi^n_{i,j,q}\right)}{(\Delta z_{q-1} + \Delta z_q)\Delta z_{q-1}}.
\end{aligned}
$$

Here, like for $\varepsilon_{i-1/2,j,q}$, $D_{i-1/2,j,q}$ is the value of $D$ evaluated in the point

$$
\left(\sum_{m=1}^{i-1}(\Delta x_m) - \Delta x_{i-1}/2, \sum_{m=1}^{j-1}(\Delta y_m), \sum_{m=1}^{q-1}(\Delta z_m)\right).
$$

### S2 Temporal development from electroneutrality to the resting state

In Figure S1, we show how the potential, charge density and ion concentrations close to the membrane change with time in the simulation used to find the resting state of the system for $L_e$= 10 nm. Initially, the potential, $\phi$, and the charge density, $\rho$ are zero, but as the $K^+$ channels open, the intracellular potential approaches a value of about $-80$ mV, and the charge density approaches about $-0.2$ C/cm$^3$ on the intracellular side of the membrane and about 0.2 C/cm$^3$ on the extracellular side.

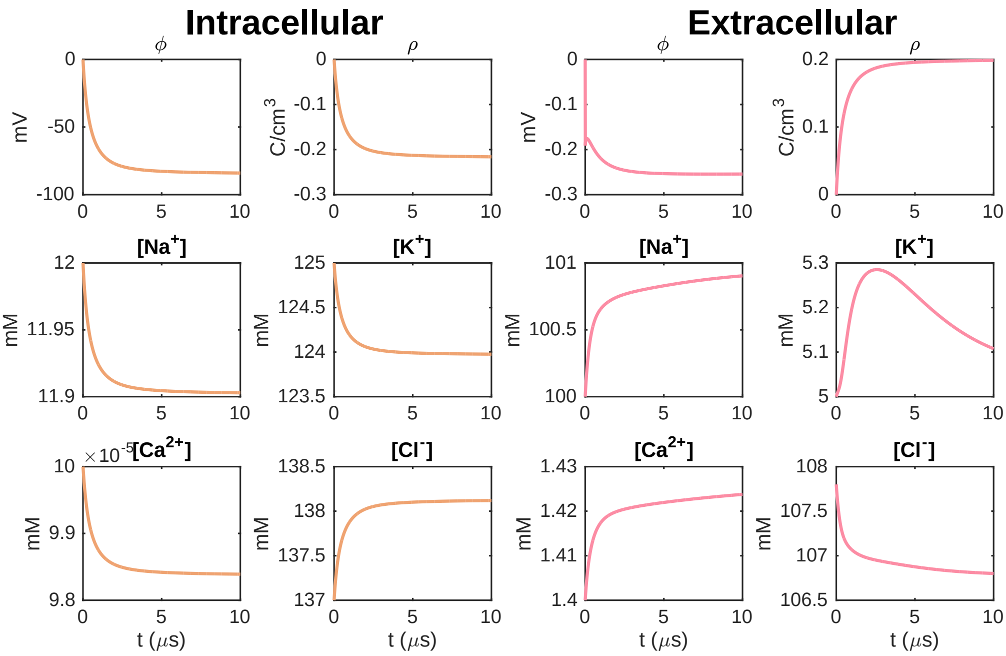

Figure S1: Illustration of how the Debye layer and the resting membrane potential evolves in the simulations with open $K^+$ channels used to find the resting state of the system. We plot the potential, charge density and ion concentrations in the first points outside of the membrane in the $x$-direction, on the intracellular and extracellular sides. In the $y$- and $z$-directions, the plotted points are located in the center of the $Na^+$ channel cluster, which is closed and thus acts as a normal membrane. We use $L_y = L_z = 300$ nm, $L_e = 10$ nm and a $K^+$ channel cluster consisting of 36 channels. In order to speed up the dynamics, we have used a scaling factor of $d_{\mathrm{K}^+}$=1 for the $K^+$ channel diffusion instead of the default value of $d_{\mathrm{K}^+}$=0.5.

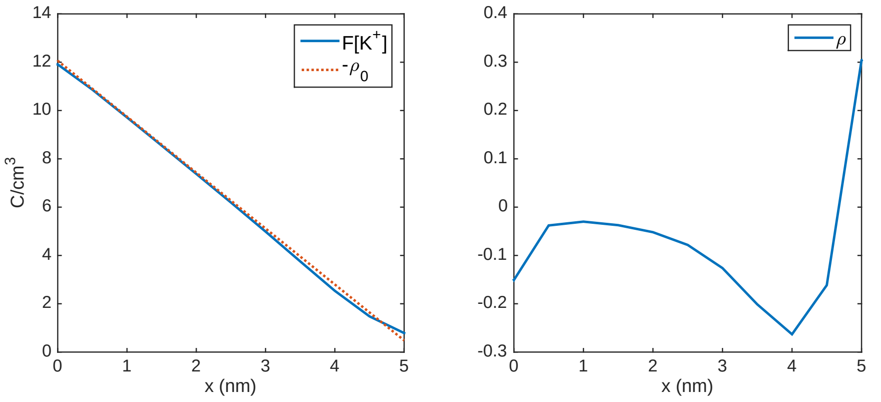

Figure S2: Illustration of the complex profile of $\rho$ in the $K^+$ channel clusters at rest. In the left panel, we show $-\rho_0$ and $F[K^+]$ along a line in the center of the $K^+$ channel cluster of the left cell at the end of the resting state simulation for $L_e = 10$ nm. We observe some small deviations between the two curves, giving rise to a non-zero $\rho = \rho_0 + F[K^+]$, illustrated in the right panel. The $x$-axis is shifted so that the $K^+$ channel cluster starts (from the intracellular side) at $x = 0$.

### S3 Profile of $\rho$ in the $K^+$ channel cluster at rest

In Figure 7 in the main paper, we observed that the charge density, $\rho$, inside of the $K^+$ channel cluster had a rather complex profile. In this section, we wish to illustrate how this profile arises as a consequence of small deviations from the completely linear profile defined as initial condition for the $K^+$ concentration in the $K^+$ channels. Recall from (4) in the main paper, that the charge density, $\rho$, is defined as

$$\rho = \rho_0 + F \sum_k z_k c_k, \tag{3}$$

where $F$ is Faraday's constant, $z_k$ is the charge of the different ion species present, and $c_k$ is the concentration of each of these ion species. Furthermore, $\rho_0$ is defined such that $\rho = 0$ in the entire domain at the beginning of the simulation. In the $K^+$ channel cluster, the concentrations of all ion species except for $K^+$ are initially set to zero and remain zero in the entire simulation, because the diffusion coefficient of all other ions that $K^+$ is set to zero in the $K^+$ channels. This means that in the $K^+$ channel cluster, $\rho$ is given by

$$\rho = \rho_0 + F[K^+], \tag{4}$$

where $[K^+]$ is the concentration of $K^+$ ions. In the left panel of Figure S2, we plot $-\rho_0$ and $F[K^+]$. We observe that the two terms are very similar, making $\rho$ close to zero. However, there are some small differences between the terms, caused by small changes in the $K^+$ concentration occurred during the simulation, and this gives rise to the profile of $\rho$ (see the right panel).

### S4 $Ca^{2+}$ and $Cl^-$ concentrations when $Na^+$ channels are open

Figures S3 and S4 show the $Ca^{2+}$ and $Cl^-$ concentrations in the simulations reported in the main paper with an open $Na^+$ channel cluster in the membrane of the left cell. The figures show the solutions in a plane in the $x$- and $y$-directions in the extracellular space close to the $Na^+$ channel clusters at the time when the change from the resting values is largest, similarly to Figures 10 and 11 in the paper, which shows the $Na^+$ and $K^+$ concentrations instead of the $Ca^{2+}$ and $Cl^-$ concentrations.

### S5 Different choice of $\rho_0$ in the channels

In order to investigate how the choice of the initial conditions and $\rho_0$ in the channel clusters affect the results, we conducted simulations with a different choice than the default linear transition used in the simulations reported in the main paper. Recall here that $\rho_0$ is set up such that $\rho$ is zero in the entire domain at $t = 0$. Moreover, the concentration of all ions are set to zero in the channel clusters except for the ion that is able to move through the channel cluster. Therefore, the profile of $\rho_0$ in the channel clusters follows directly from the initial conditions in the channel clusters.

Figures S5–S13 show the results of simulations using a piecewise constant concentration as initial conditions in the channel clusters. More specifically, the initial concentration is set equal to the intracellular concentration in the half of the channel channel that is closest to the intracellular space and equal to the extracellular concentration in the half that is closest to the extracellular space. The results of the simulations using this set up seem to be quite close to the results obtained using the linear profiles used in the simulations shown in the main paper, except for the profile of $\phi$, $\rho$ and the ion concentrations in and close to the $K^+$ channel clusters at rest (see Figures S5 and S6). In addition, the maximum changes in the extracellular potential seem to be somewhat smaller for this case (compare, e.g., Figure 12 in the main paper and Figure S13).

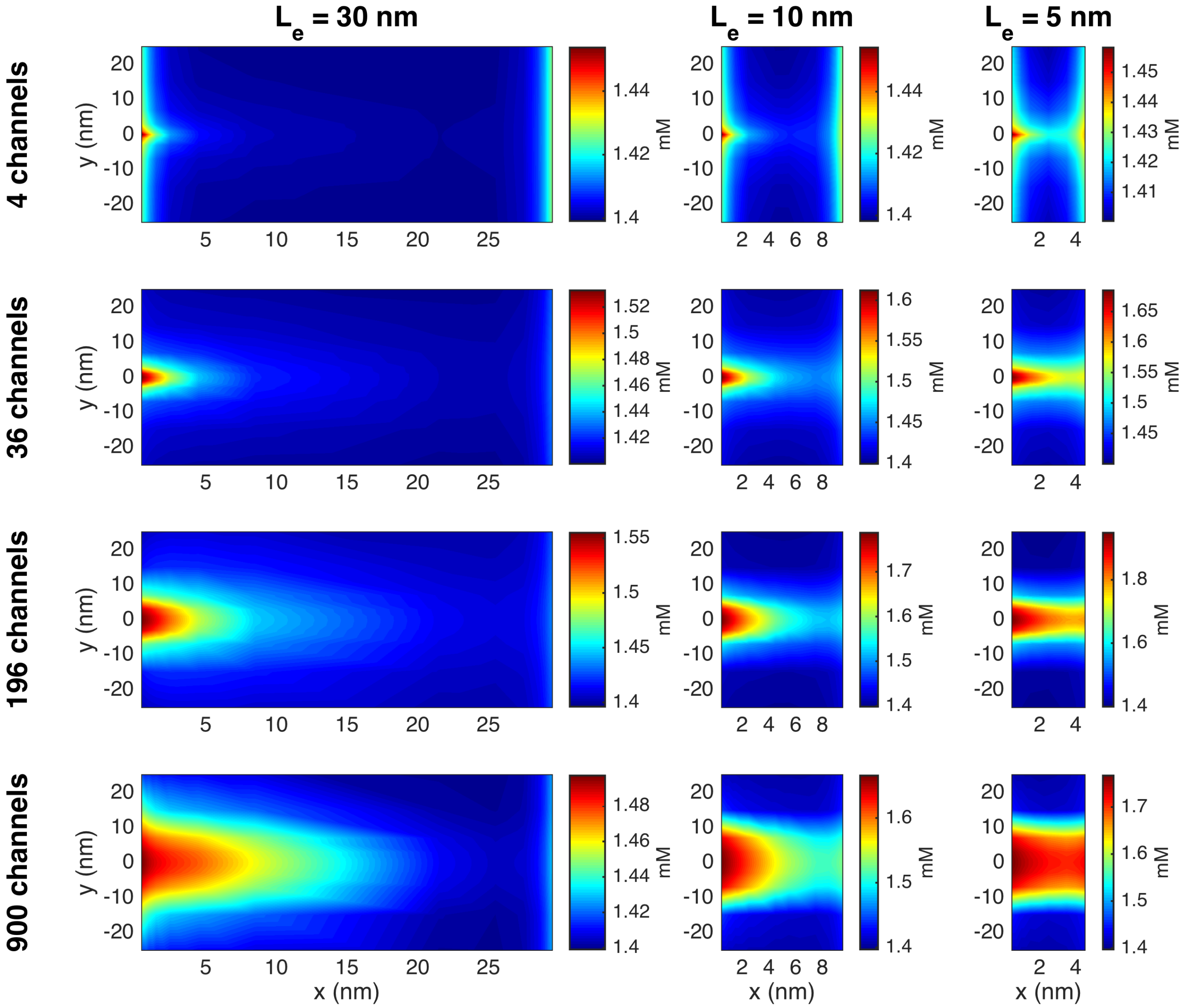

Figure S3: The $Ca^{2+}$ concentration in the extracellular space between two cells in simulations with an open $Na^+$ channel clusters on the membrane of the left cell. The width of the extracellular space, $L_e$, and the size of the $Na^+$ channel cluster is varied in the columns and rows of the figure, respectively. The plots show the solution in the $(x, y)$-plane for the center of the domain in the $z$-direction at the point in time when the deviation from rest is largest. In the $y$-direction, we focus on the 50 nm closest to the $Na^+$ channel clusters. The coordinates on the axes are shifted so that $x = 0$ marks the end of the membrane of the left cell and $y = 0$ marks the center of the $Na^+$ channels. Note that the scaling of the colorbar is different for the different cases.

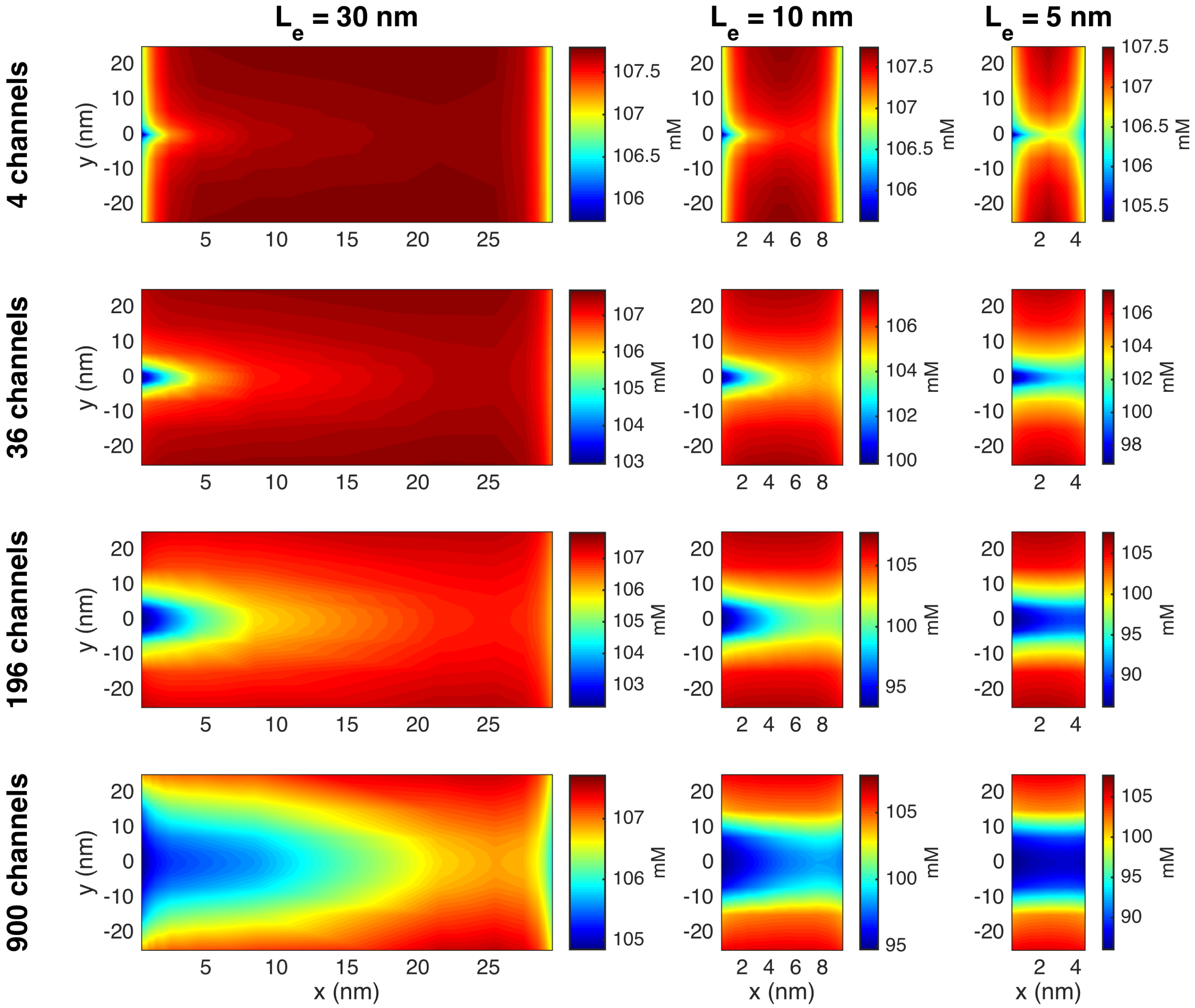

Figure S4: The $Cl^-$ concentration in the extracellular space between two cells in simulations with an open $Na^+$ channel cluster on the membrane of the left cell. The figure setup is the same as for Figure S3.

### S6 A different set of boundary conditions

In order to investigate how the choice of boundary conditions affect the results, Figures S14–S22 show the results of simulations using a different set of boundary conditions than those used in the remaining simulations of this study. These boundary conditions are described in Section 2.3.3 in the main paper.

More specifically, we apply a Dirichlet boundary condition of

$$\phi = E_{\mathrm{K}} \tag{5}$$

on the rightmost boundary of the right cell. Here, $E_{\mathrm{K}}$ is the Nernst equilibrium potential for $\mathrm{K}^+$ ions, i.e.,

$$E_{\mathrm{K}} = \frac{k_B T}{z_{\mathrm{K}^+} e} \ln \left( \frac{c_{\mathrm{K}^+,e}}{c_{\mathrm{K}^+,i}} \right), \tag{6}$$

where $k_B$, $T$, $z_{\mathrm{K}^+}$ and $e$ are parameters specified in the paper, and $c_{\mathrm{K}^+,i}$ is the intracellular $\mathrm{K}^+$ concentration. Moreover, $c_{\mathrm{K}^+,e}$ is the extracellular $\mathrm{K}^+$ concentration measured in the extracellular point $(L_i+L_m+L_e/2, 0, 0)$ at the end of the simulation used to find the resting state. On the remaining part of the outer boundary of the computational domain, we apply the Neumann boundary condition

$$\varepsilon \nabla \phi \cdot \mathbf{n} = 0. \tag{7}$$

In addition, we apply no-flux Neumann boundary conditions for the ionic concentrations on the entire boundary of the computational domain

$$D_k \nabla c_k \cdot \mathbf{n} = 0, \text{ for } k = \{\mathrm{Na}^+, \mathrm{K}^+, \mathrm{Ca}^{2+} \text{ and } \mathrm{Cl}^-\}. \tag{8}$$

Comparing the results of Figures S14–S22 to the corresponding figures for the default boundary conditions described in the paper, we observe that the results seem to be very similar for the two sets of boundary conditions. The perhaps largest difference between the two cases is that the extracellular $\mathrm{K}^+$ concentration at rest for a narrow cleft of $L_e = 5$ nm is about 1 mM higher in the simulations with Neumann boundary conditions everywhere for the concentrations than for the default boundary conditions (compare Figure 6 in the main paper and Figure S14).

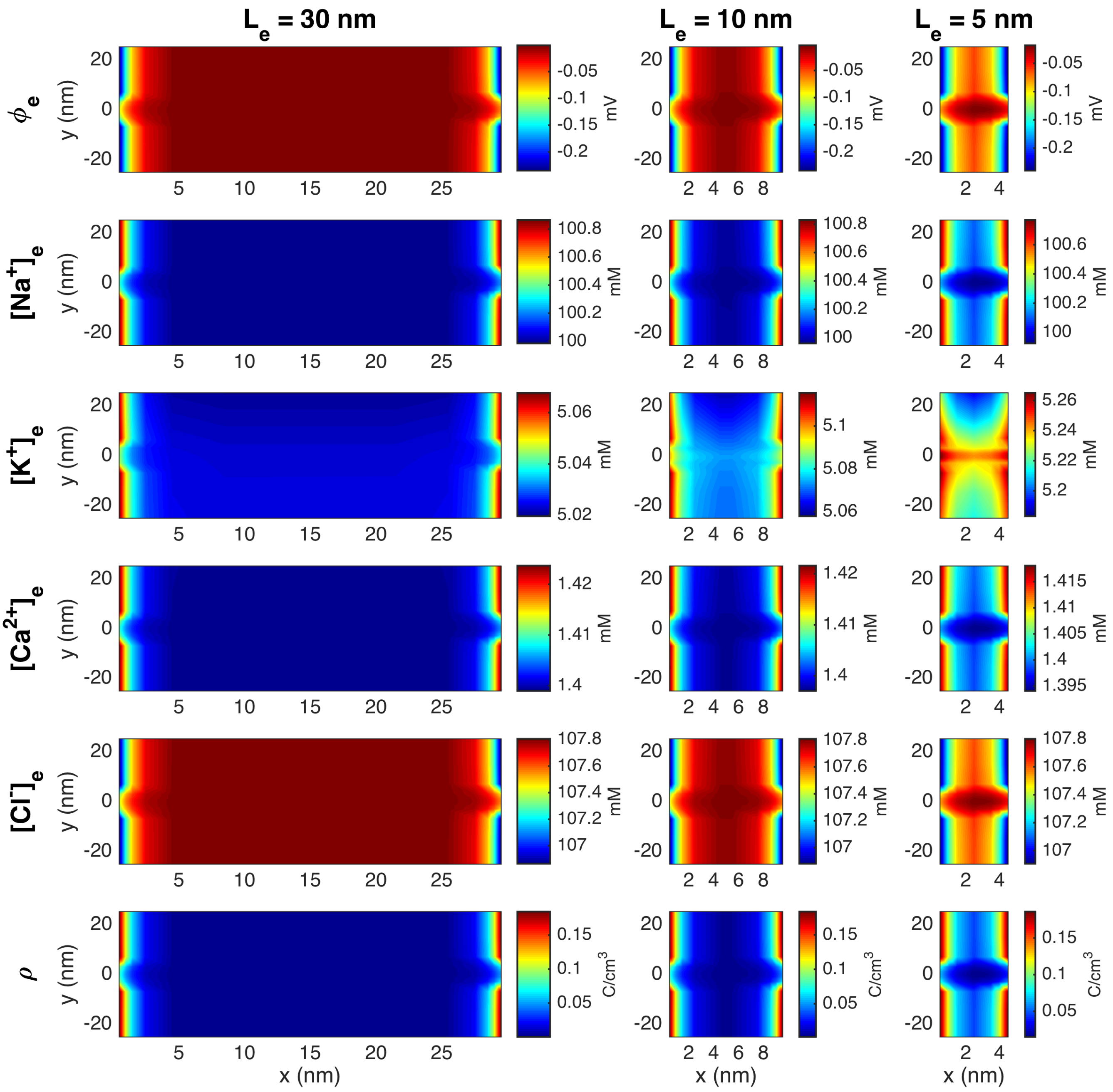

Figure S5: Stationary solution for the potential, $\phi$, the concentration of $Na^+$, $K^+$, $Ca^{2+}$, and $Cl^-$ ions, and the charge density, $\rho$, in the extracellular space between the two cells in simulations with open $K^+$ channels, but closed $Na^+$ channels. The figure corresponds to Figure 6 in the paper, but with a different profile for the initial conditions and $\rho_0$ in the ion channel clusters. Specifically, the initial condition in the channel clusters is a equal to the intracellular concentration in the half of the channel cluster that is closest to the intracellular space and equal to the extracellular concentration in the half that is closest to the extracellular space.

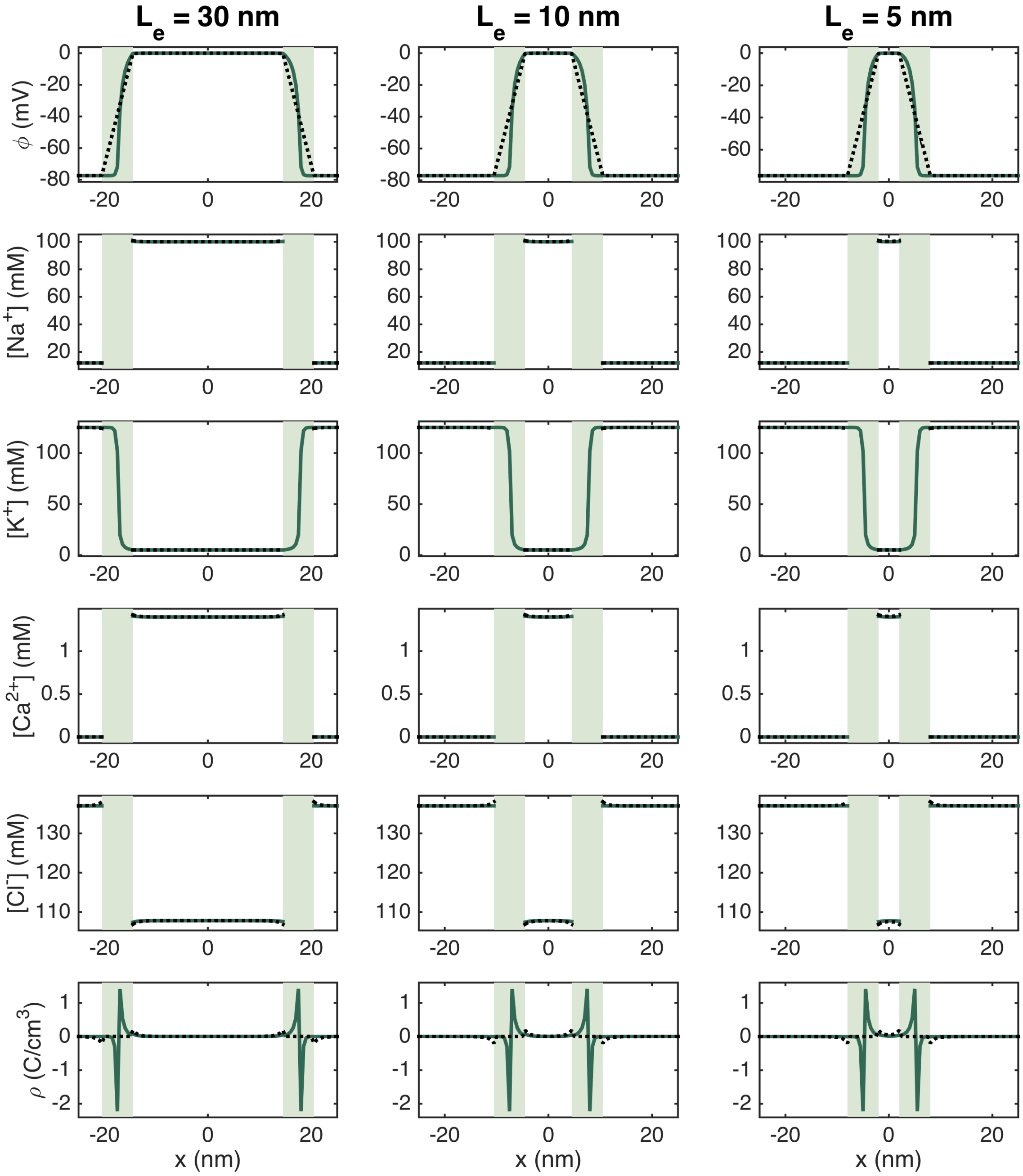

Figure S6: Stationary solution of the potential, $\phi$, the concentration of $Na^+$, $K^+$, $Ca^{2+}$, and $Cl^-$ ions, and the charge density, $\rho$, along lines in the $x$-direction for open $K^+$ channels and closed $Na^+$ channels. The full green line represents the solution along a line crossing through the $K^+$ channels and the dotted black line represents the solution along a line about 100 nm below the $K^+$ channel cluster. The light green areas mark the membrane. The figure corresponds to Figure 7 in the paper, but with a different profile for the initial conditions and $\rho_0$ in the ion channel clusters. Specifically, the initial condition in the channel clusters is a equal to the intracellular concentration in the half of the channel cluster that is closest to the intracellular space and equal to the extracellular concentration in the half that is closest to the extracellular space.

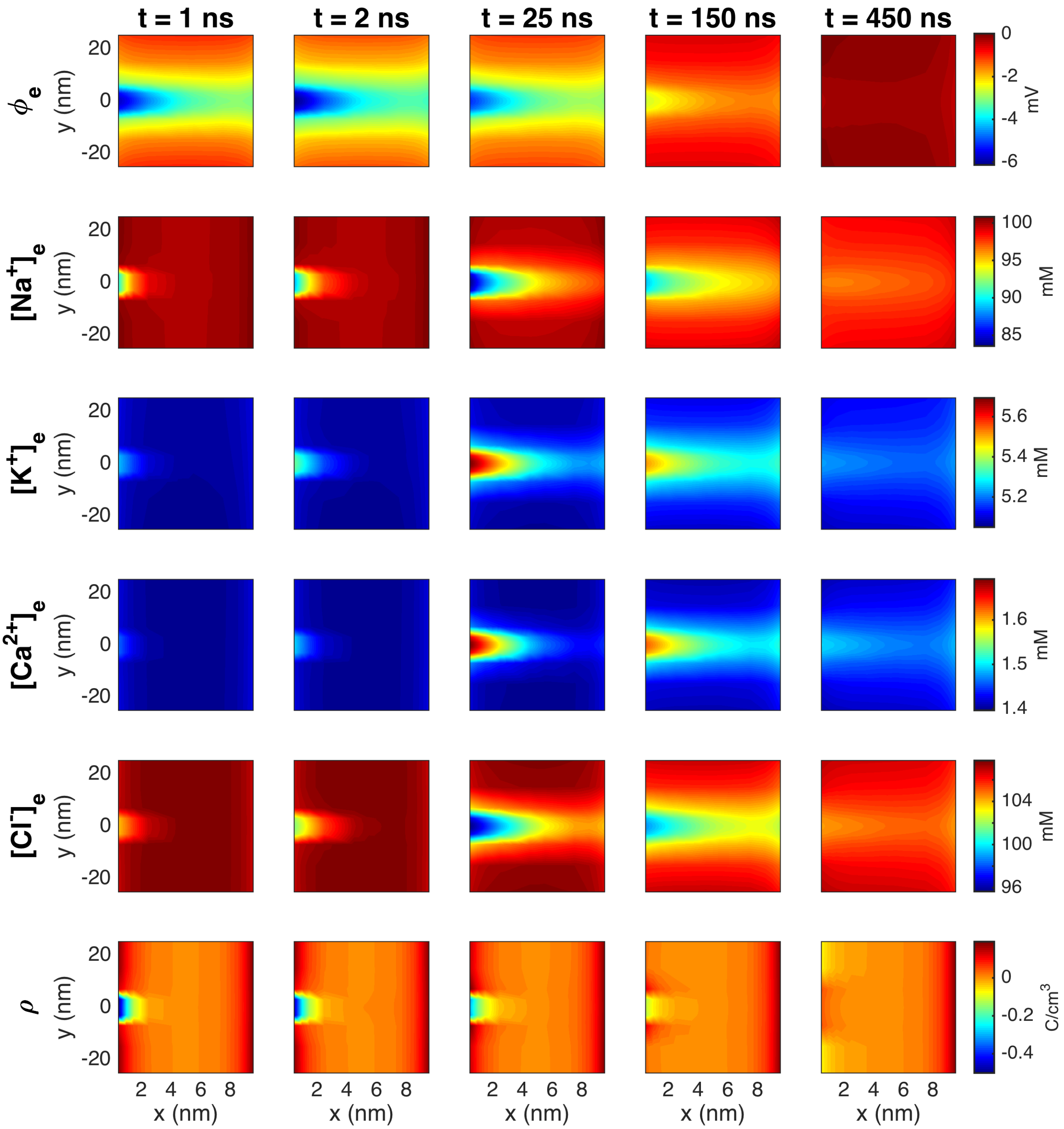

Figure S7: The PNP model solution in the extracellular space between the two cells in a simulation with $L_e = 10$ nm and an $Na^+$ channel cluster of 196 channels on the membrane of the left cell. The figure corresponds to Figure 8 in the paper, but with a different profile for the initial conditions and $\rho_0$ in the ion channel clusters. Specifically, the initial condition in the channel clusters is a equal to the intracellular concentration in the half of the channel cluster that is closest to the intracellular space and equal to the extracellular concentration in the half that is closest to the extracellular space.

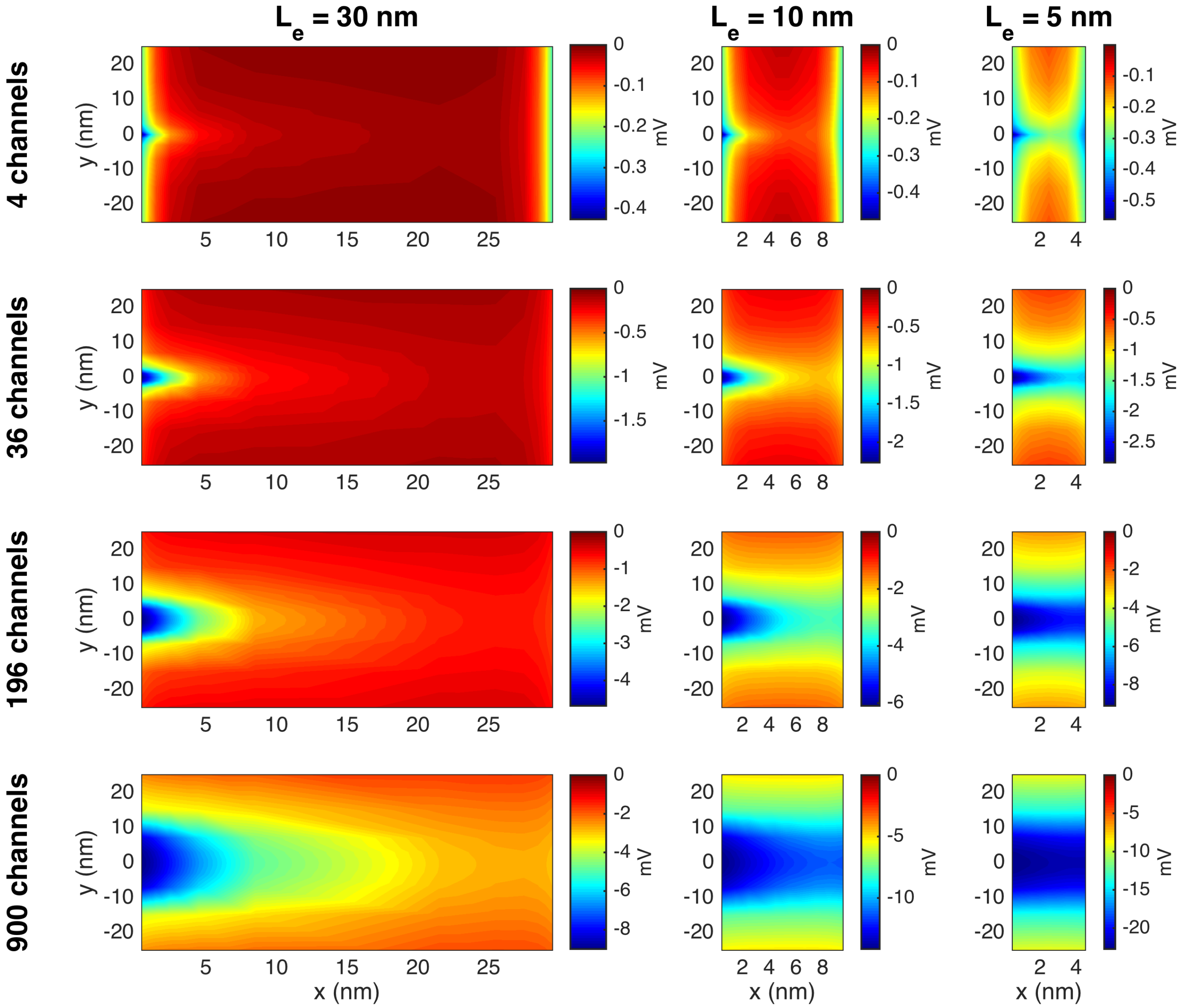

Figure S8: The potential, $\phi$, in the extracellular space between the two cells in simulations with open $Na^+$ channel clusters on the membrane of the left cell at the point in time when the most negative potential occurs. The figure corresponds to Figure 9 in the paper, but with a different profile for the initial conditions and $\rho_0$ in the ion channel clusters. Specifically, the initial condition in the channel clusters is a equal to the intracellular concentration in the half of the channel cluster that is closest to the intracellular space and equal to the extracellular concentration in the half that is closest to the extracellular space.

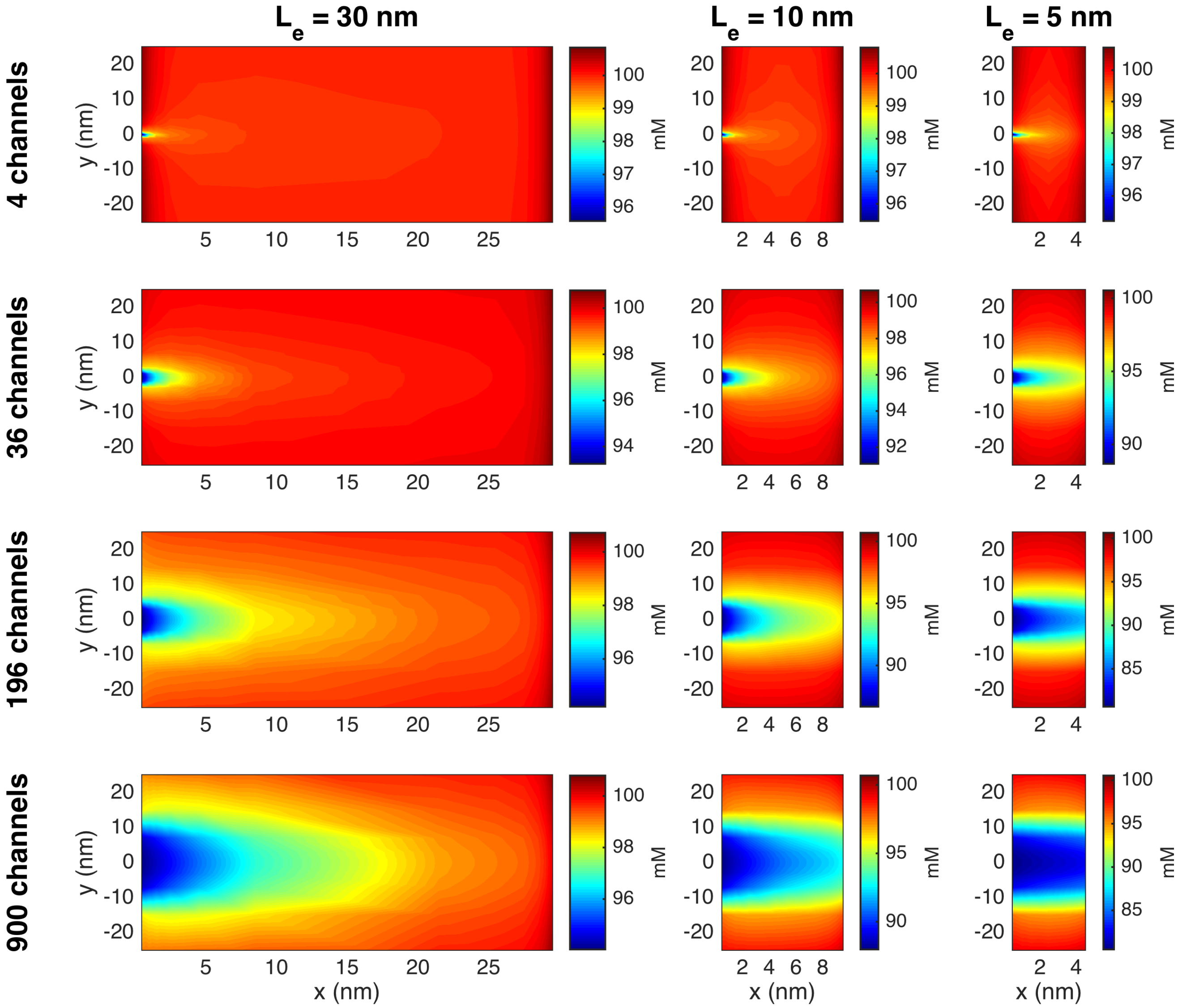

Figure S9: The $Na^+$ concentration in the extracellular space between the two cells in simulations with open $Na^+$ channel clusters on the membrane of the left cell at the point in time when the larges deviation from rest occurs. The figure corresponds to Figure 10 in the paper, but with a different profile for the initial conditions and $\rho_0$ in the ion channel clusters. Specifically, the initial condition in the channel clusters is a equal to the intracellular concentration in the half of the channel cluster that is closest to the intracellular space and equal to the extracellular concentration in the half that is closest to the extracellular space.

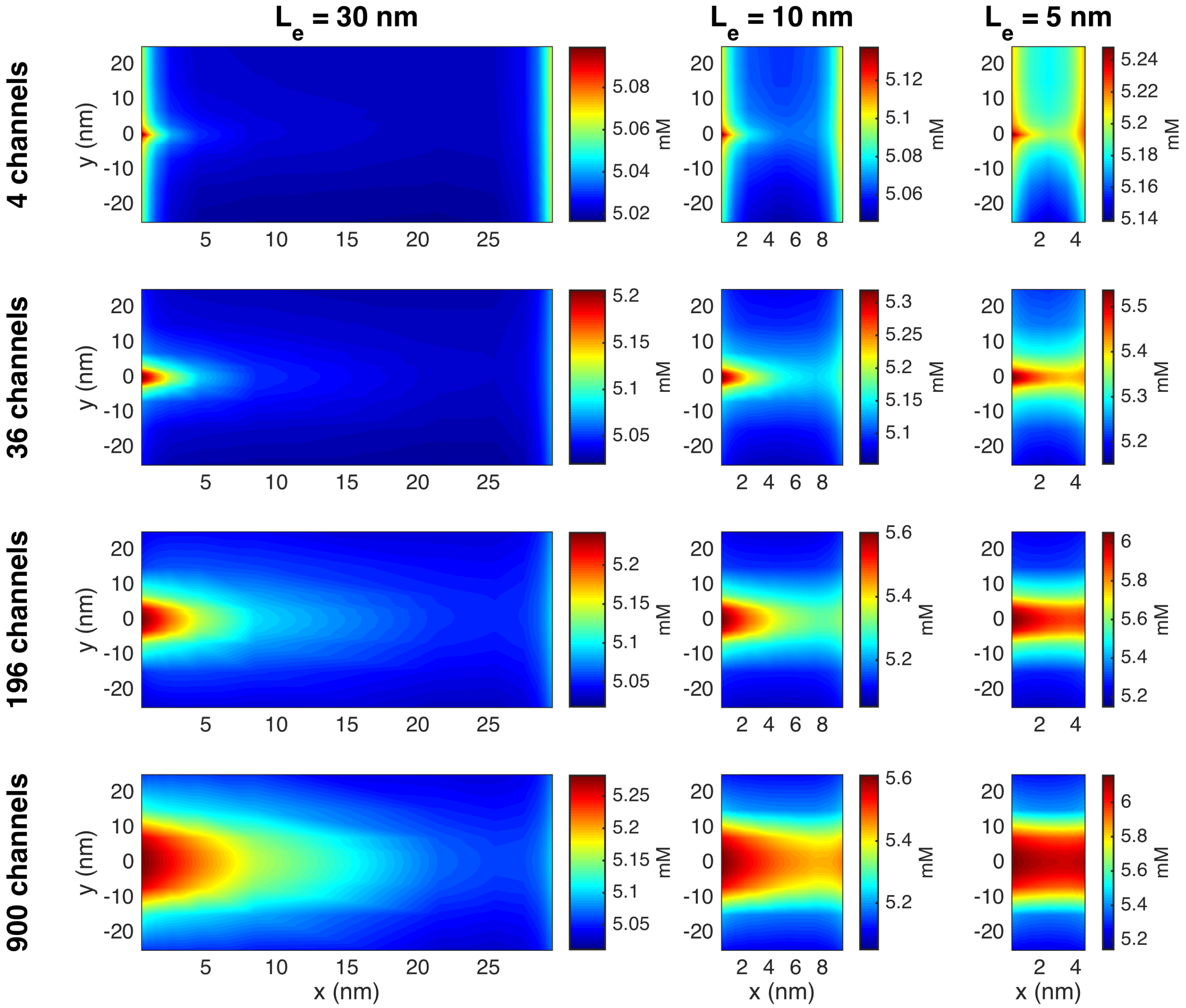

Figure S10: The $K^+$ concentration in the extracellular space between the two cells in simulations with open $Na^+$ channel clusters on the membrane of the left cell at the point in time when the larges deviation from rest occurs. The figure corresponds to Figure 11 in the paper, but with a different profile for the initial conditions and $\rho_0$ in the ion channel clusters. Specifically, the initial condition in the channel clusters is a equal to the intracellular concentration in the half of the channel cluster that is closest to the intracellular space and equal to the extracellular concentration in the half that is closest to the extracellular space.

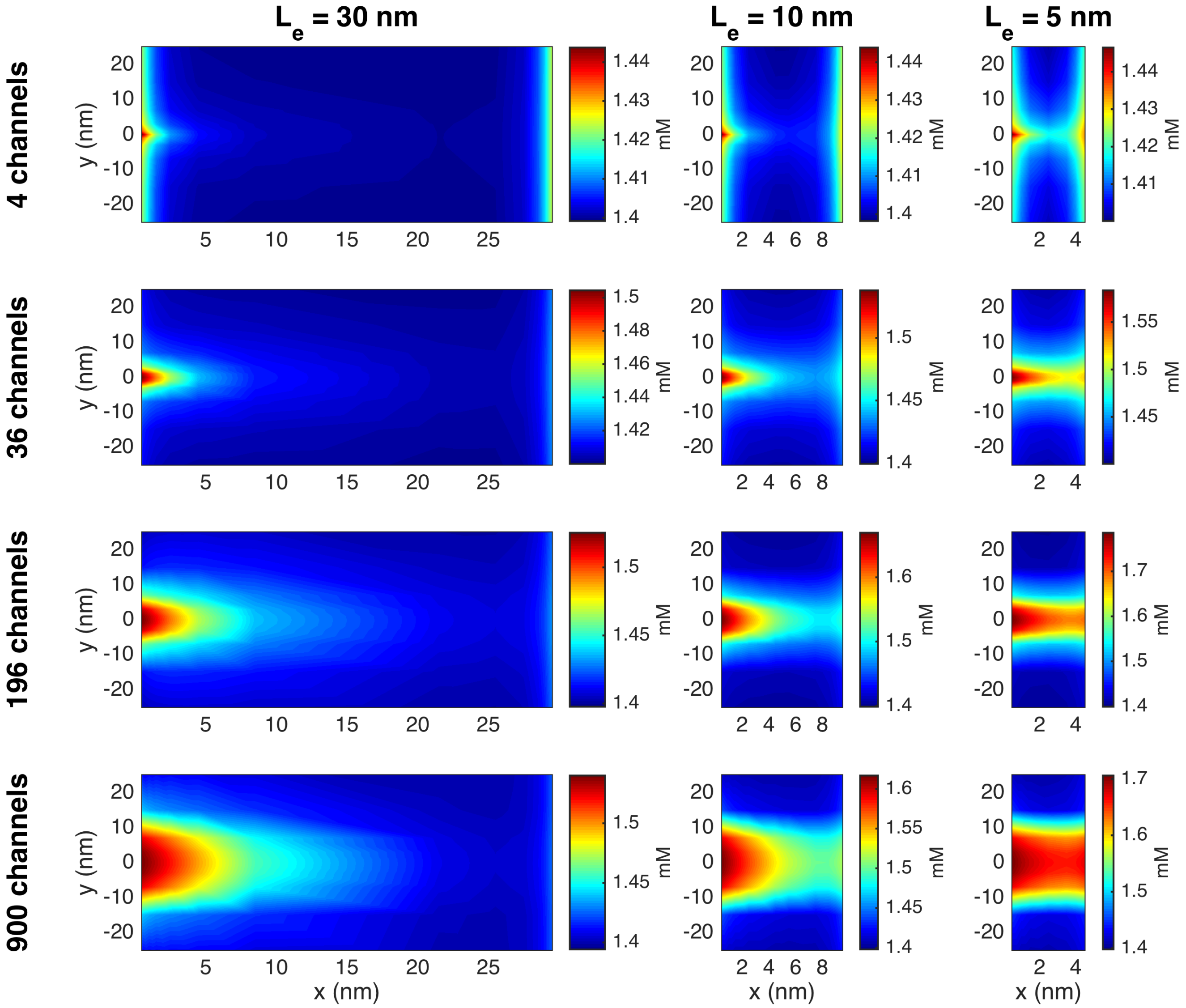

Figure S11: The $Ca^{2+}$ concentration in the extracellular space between the two cells in simulations with open $Na^{+}$ channel clusters on the membrane of the left cell at the point in time when the larges deviation from rest occurs. The figure corresponds to Figure S3, but with a different profile for the initial conditions and $\rho_0$ in the ion channel clusters. Specifically, the initial condition in the channel clusters is a equal to the intracellular concentration in the half of the channel cluster that is closest to the intracellular space and equal to the extracellular concentration in the half that is closest to the extracellular space.

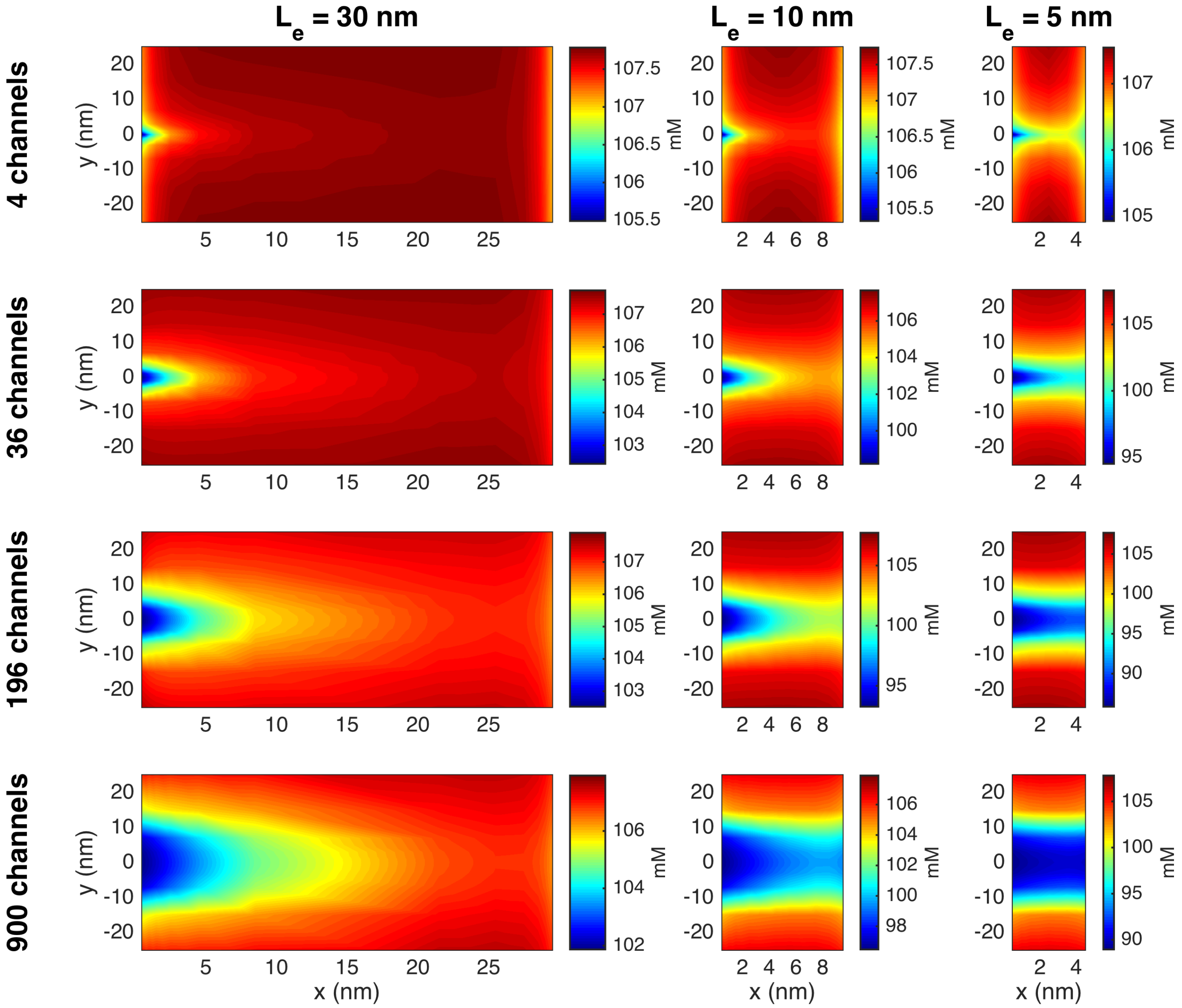

Figure S12: The $Cl^-$ concentration in the extracellular space between the two cells in simulations with open $Na^+$ channel clusters on the membrane of the left cell at the point in time when the larges deviation from rest occurs. The figure corresponds to Figure S4, but with a different profile for the initial conditions and $\rho_0$ in the ion channel clusters. Specifically, the initial condition in the channel clusters is a equal to the intracellular concentration in the half of the channel cluster that is closest to the intracellular space and equal to the extracellular concentration in the half that is closest to the extracellular space.

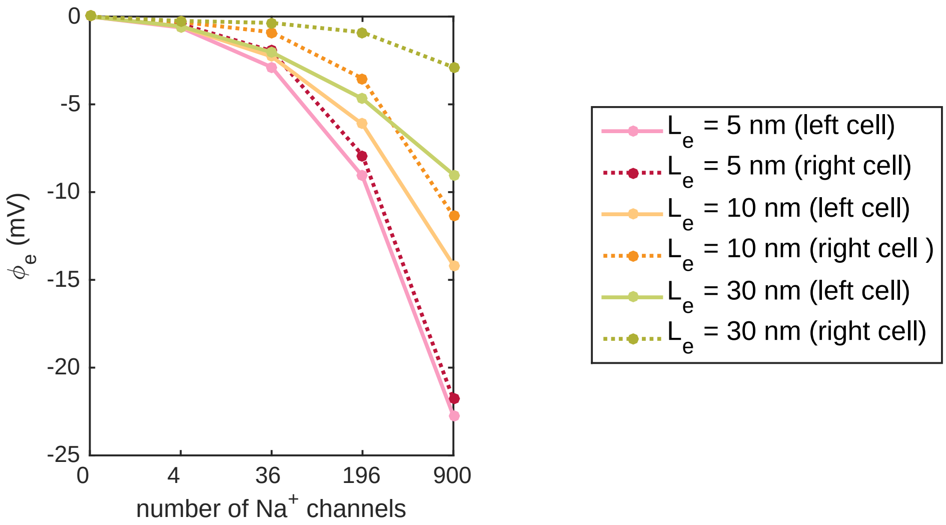

Figure S13: Most negative extracellular potential outside of the $Na^+$ channel clusters in simulations with different widths of the extracellular space between the cells, $L_e$, and different sizes of the $Na^+$ channel clusters. The figure corresponds to Figure 12 in the paper, but with a different profile for the initial conditions and $\rho_0$ in the ion channel clusters. Specifically, the initial condition in the channel clusters is a equal to the intracellular concentration in the half of the channel cluster that is closest to the intracellular space and equal to the extracellular concentration in the half that is closest to the extracellular space.

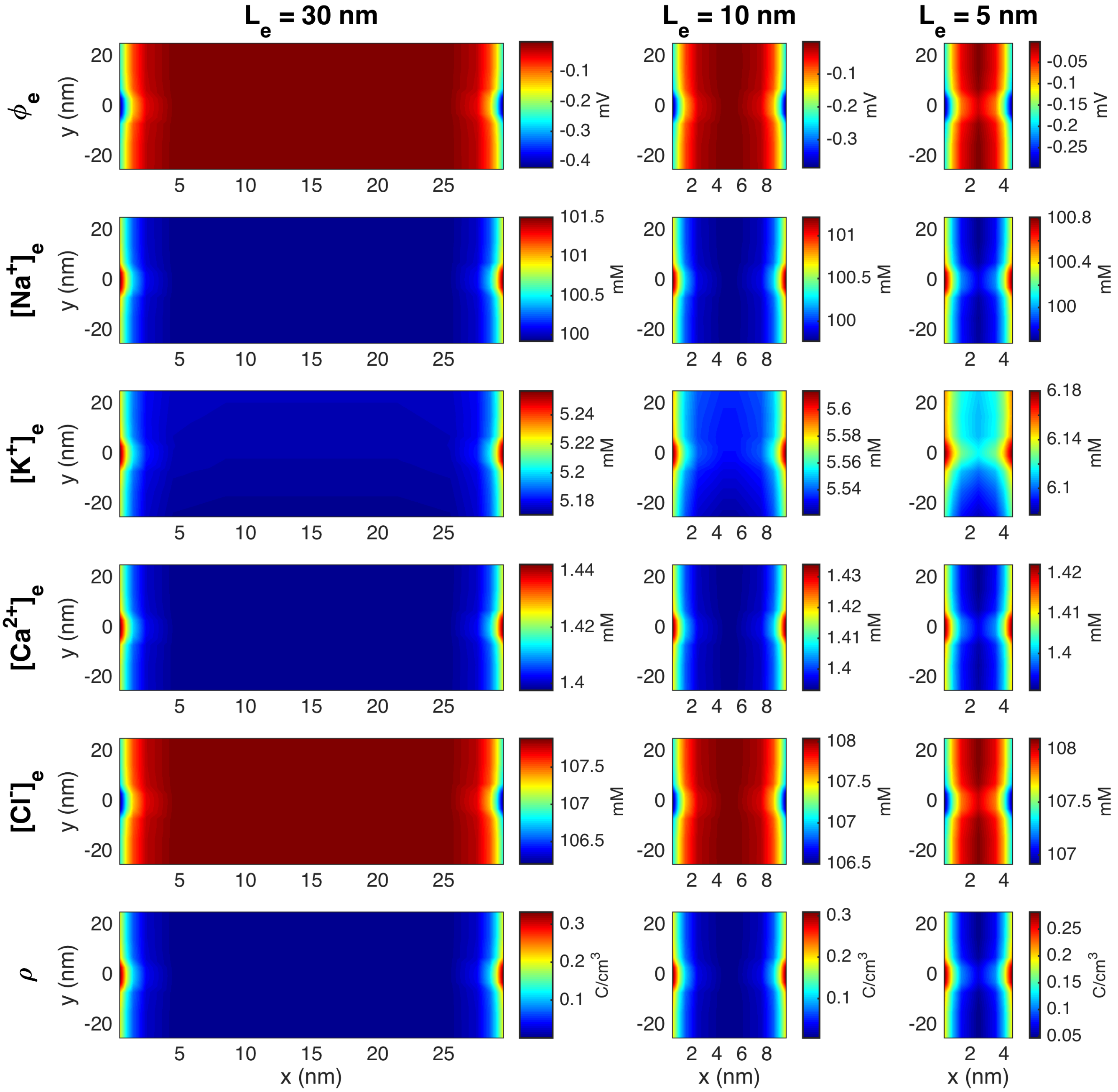

Figure S14: Stationary solution of the potential, $\phi$, the concentration of $Na^+$, $K^+$, $Ca^{2+}$, and $Cl^-$ ions, and the charge density, $\rho$, in the extracellular space between the two cells in simulations with open $K^+$ channels, but closed $Na^+$ channels. The figure corresponds to Figure 6 in the paper, but with a different set of boundary conditions applied. Specifically, no-flux Neumann boundary conditions are applied for the ionic concentrations on all outer boundaries of the computational domain.

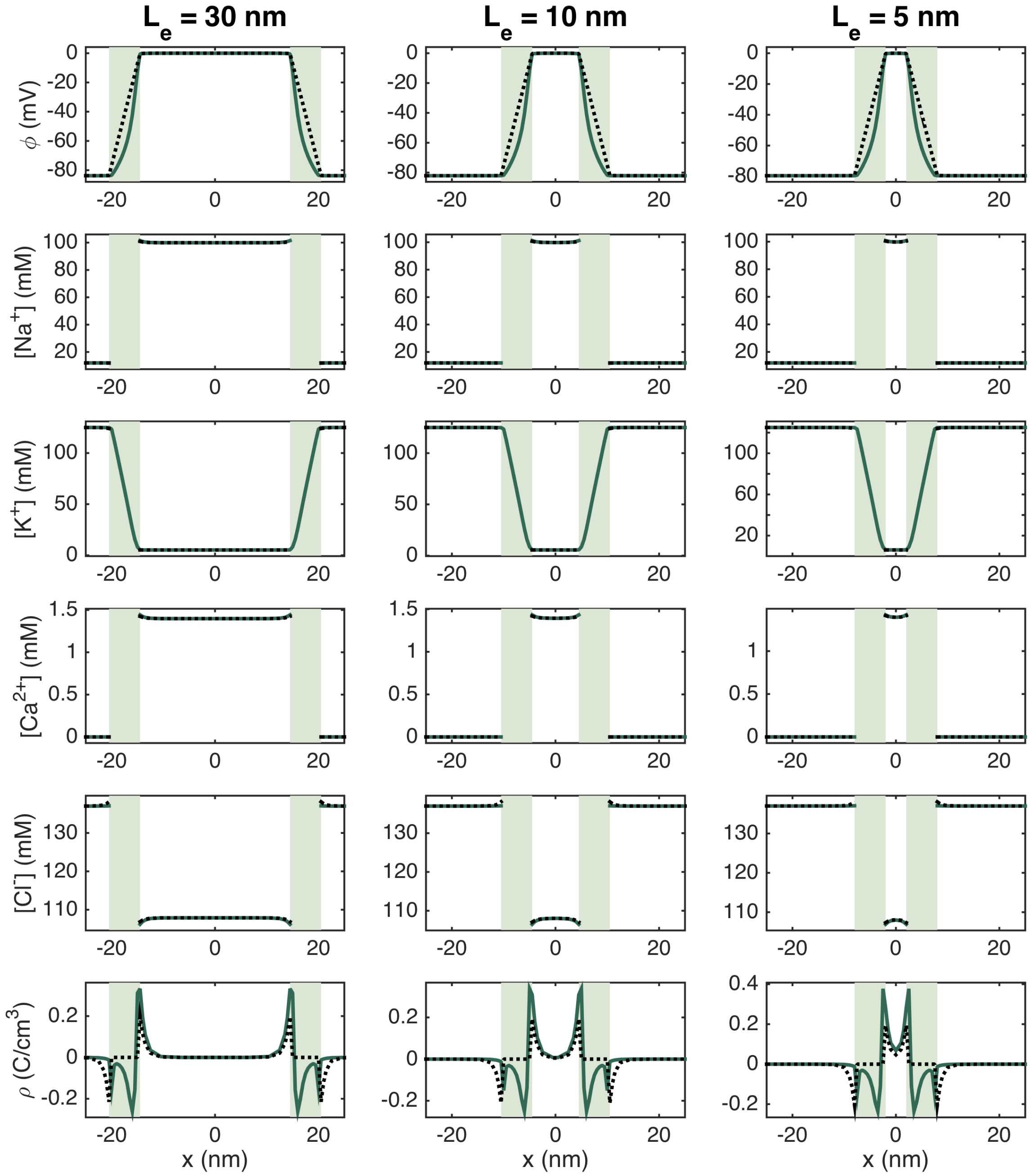

Figure S15: Stationary solution of the potential, $\phi$, the concentration of $Na^+$, $K^+$, $Ca^{2+}$, and $Cl^-$ ions, and the charge density, $\rho$, along lines in the $x$-direction for open $K^+$ channels and closed $Na^+$ channels. The full green line represents the solution along a line crossing through the $K^+$ channels and the dotted black line represents the solution along a line about 100 nm below the $K^+$ channel cluster. The light green areas mark the membrane. The figure corresponds to Figure 7 in the paper, but with a different set of boundary conditions applied. Specifically, no-flux Neumann boundary conditions are applied for the ionic concentrations on all outer boundaries of the computational domain.

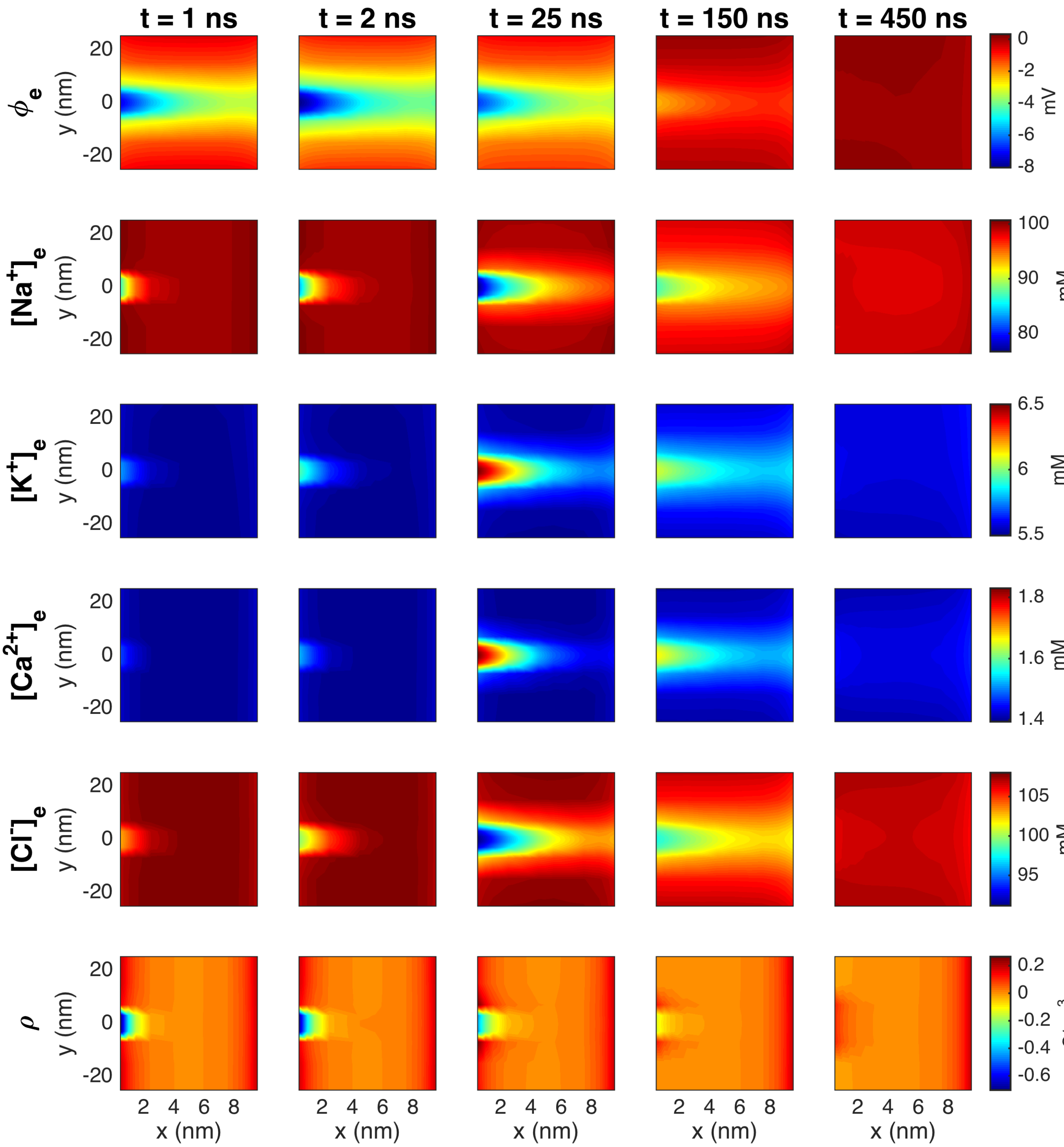

Figure S16: The PNP model solution in the extracellular space between the two cells in a simulation with $L_e = 10$ nm and an $Na^+$ channel cluster of 196 channels on the membrane of the left cell. The figure corresponds to Figure 8 in the paper, but with a different set of boundary conditions applied. Specifically, no-flux Neumann boundary conditions are applied for the ionic concentrations on all outer boundaries of the computational domain.

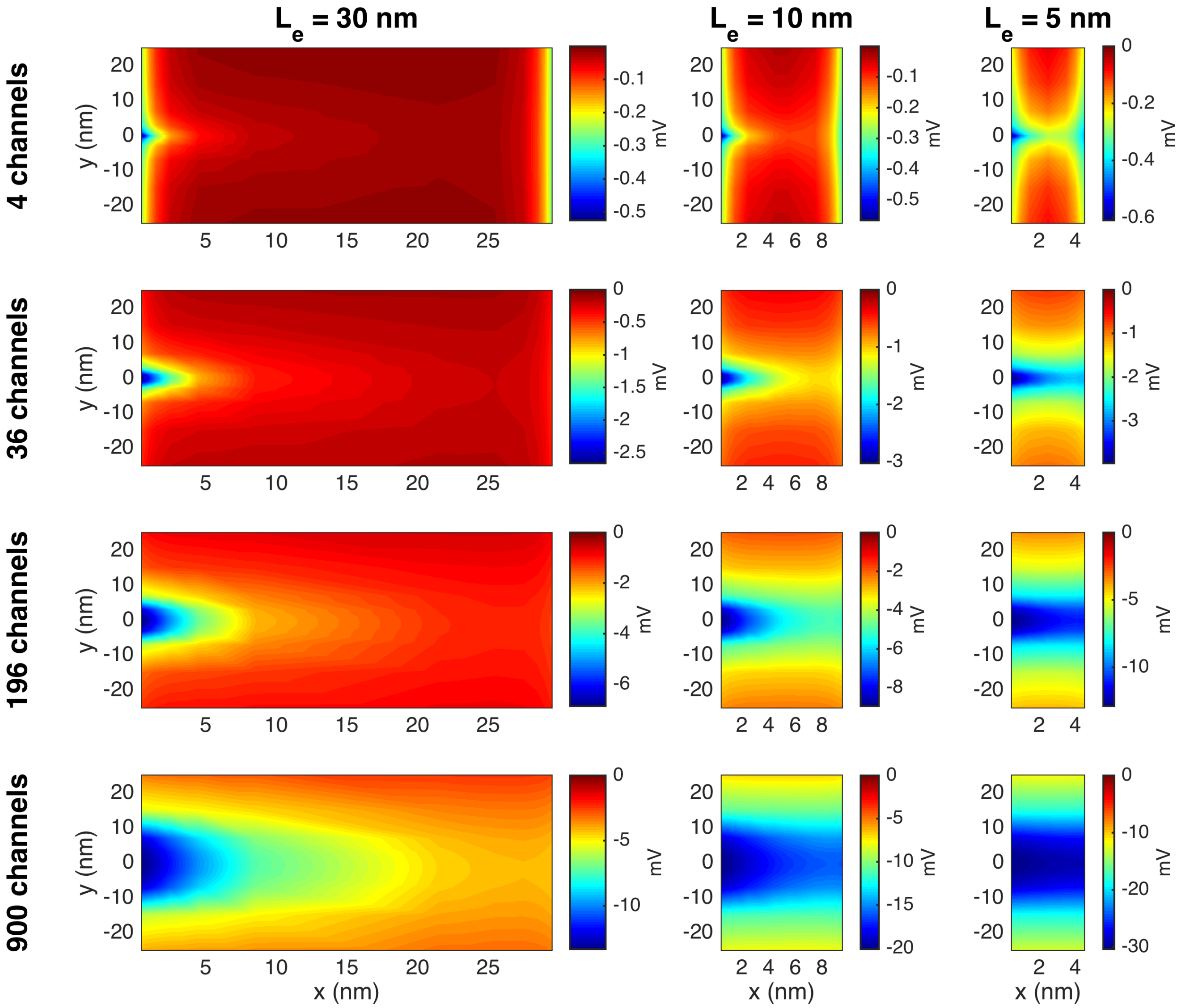

Figure S17: The potential, $\phi$, in the extracellular space between the two cells in simulations with open $Na^+$ channel clusters on the membrane of the left cell at the point in time when the most negative potential occurs. The figure corresponds to Figure 9 in the paper, but with a different set of boundary conditions applied. Specifically, no-flux Neumann boundary conditions are applied for the ionic concentrations on all outer boundaries of the computational domain.

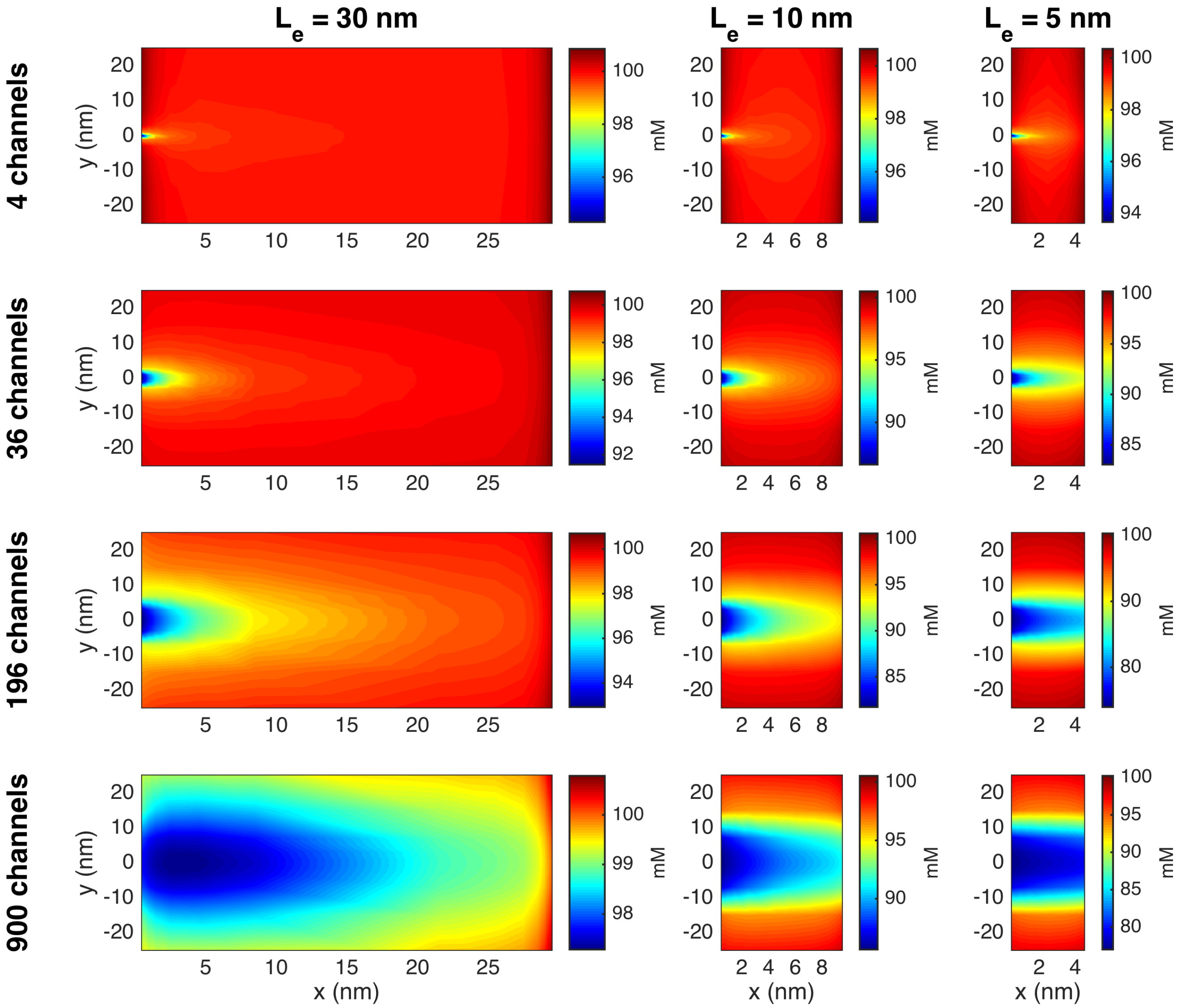

Figure S18: The $Na^+$ concentration in the extracellular space between the two cells in simulations with open $Na^+$ channel clusters on the membrane of the left cell at the point in time when the larges deviation from rest occurs. The figure corresponds to Figure 10 in the paper, but with a different set of boundary conditions applied. Specifically, no-flux Neumann boundary conditions are applied for the ionic concentrations on all outer boundaries of the computational domain.

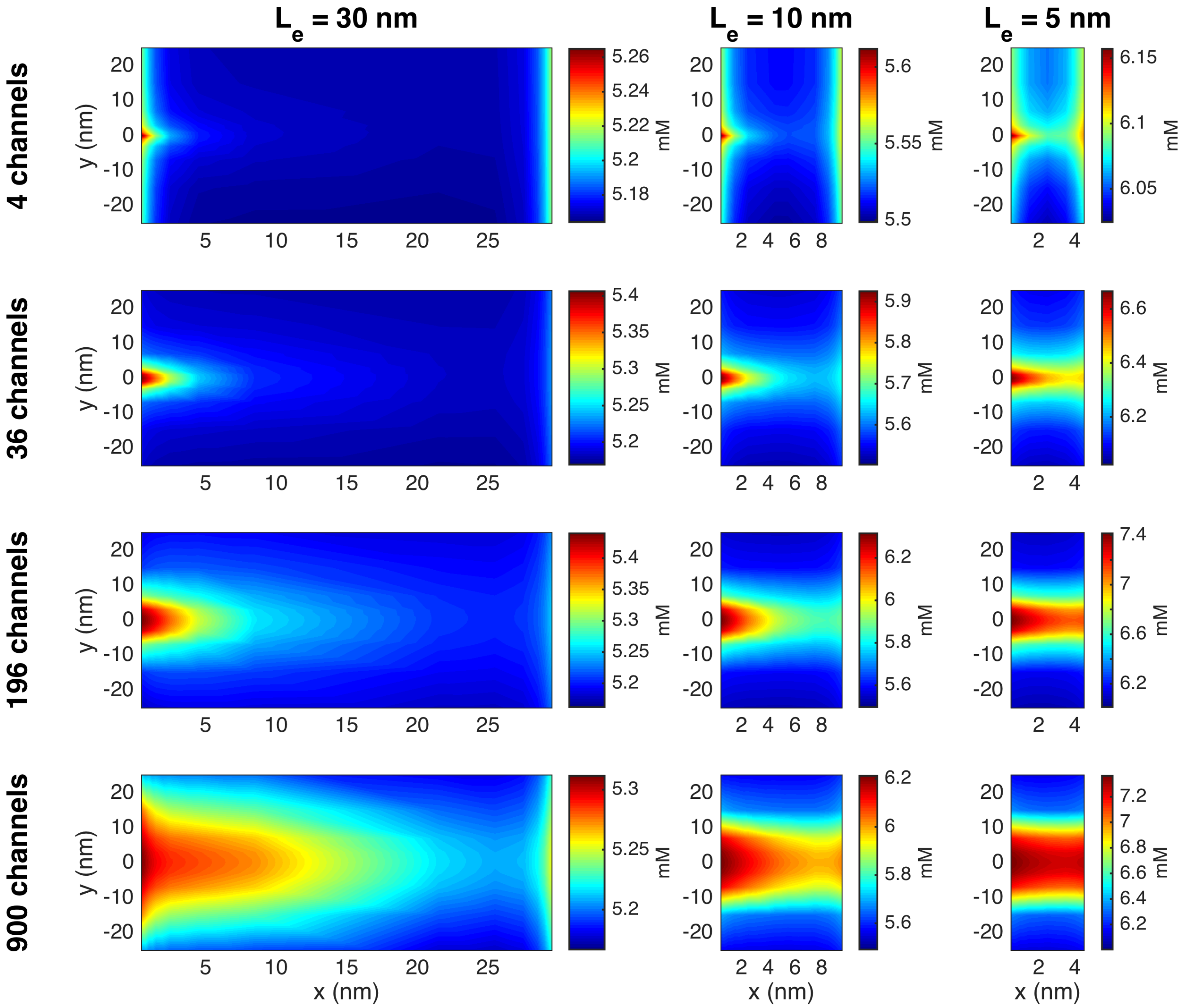

Figure S19: The $K^+$ concentration in the extracellular space between the two cells in simulations with open $Na^+$ channel clusters on the membrane of the left cell at the point in time when the larges deviation from rest occurs. The figure corresponds to Figure 11 in the paper, but with a different set of boundary conditions applied. Specifically, no-flux Neumann boundary conditions are applied for the ionic concentrations on all outer boundaries of the computational domain.

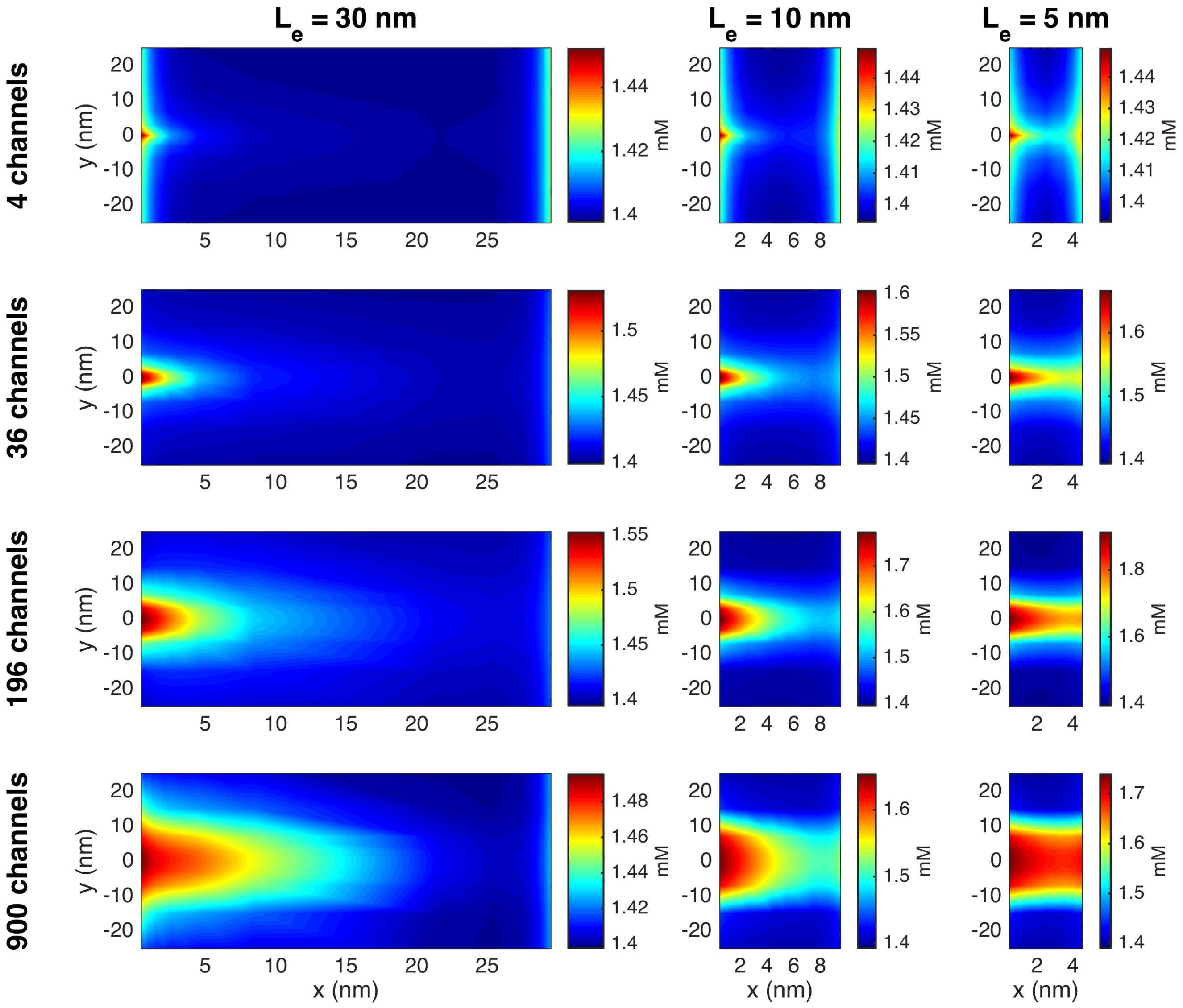

Figure S20: The $Ca^{2+}$ concentration in the extracellular space between the two cells in simulations with open $Na^+$ channel clusters on the membrane of the left cell at the point in time when the larges deviation from rest occurs. The figure corresponds to Figure S3, but with a different set of boundary conditions applied. Specifically, no-flux Neumann boundary conditions are applied for the ionic concentrations on all outer boundaries of the computational domain.

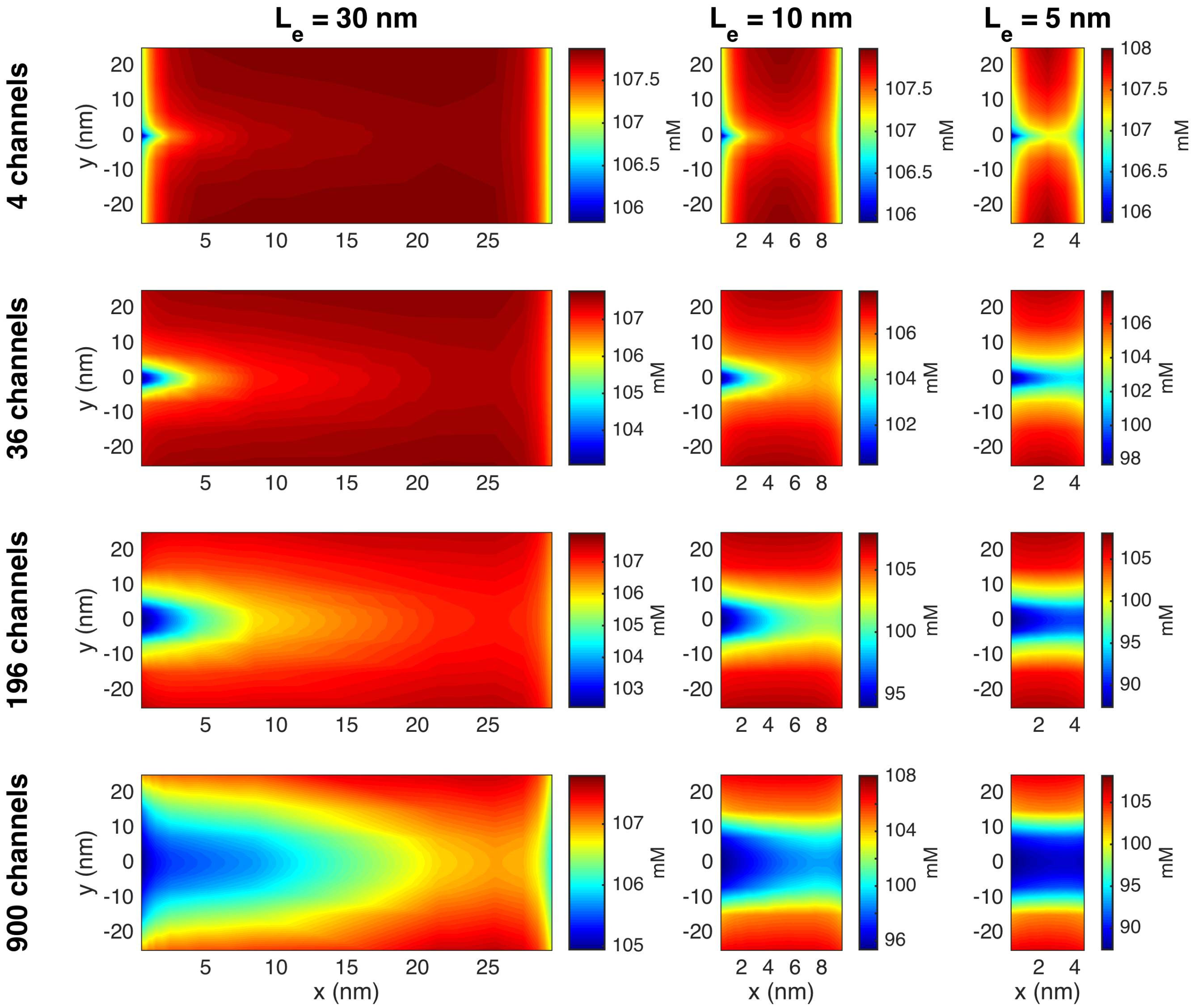

Figure S21: The $Cl^-$ concentration in the extracellular space between the two cells in simulations with open $Na^+$ channel clusters on the membrane of the left cell at the point in time when the larges deviation from rest occurs. The figure corresponds to Figure S4, but with a different set of boundary conditions applied. Specifically, no-flux Neumann boundary conditions are applied for the ionic concentrations on all outer boundaries of the computational domain.

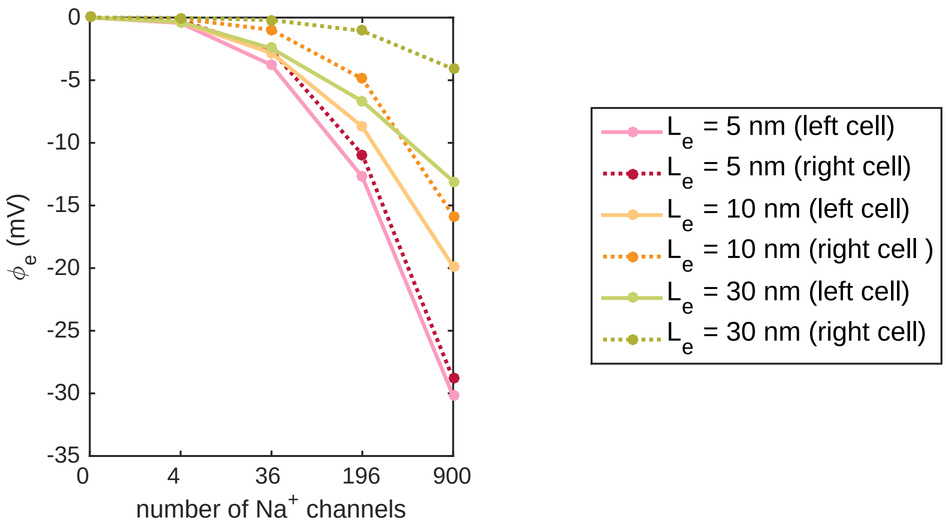

Figure S22: Most negative extracellular potential outside of the $Na^+$ channel clusters in simulations with different widths of the extracellular space between the cells, $L_e$, and different sizes of the $Na^+$ channel clusters. The figure corresponds to Figure 12 in the main paper, but with a different set of boundary conditions applied. Specifically, no-flux Neumann boundary conditions are applied for the ionic concentrations on all outer boundaries of the computational domain.
